# Genetically-encoded discovery and development of peptide-macrocycle imaging agents for PD-L1

**DOI:** 10.64898/2026.09.11.750985

**Authors:** Ratmir Derda, Vincent Albert, Matthew L. Michnik, Gary Bao, KC Tara Bahadur, William Kirby, Haley Irwin, Kristen Maiorana, James Walker, Govinda Sharma, Rima Mistry, Abdelrhman Zakaria, Danial Yazdan, Jiahui Wang, Martin Truksa, Ye Cai, Jenilee Woodfield, Man-jot Kaur, Zoe O’Gara, Andreas Dorian, Melinda Wuest, Cody Bergman, John S. Klassen, Hyun W. Kim, Thomas K. Sawyer, Todd McMullen, Frank Wuest

## Abstract

The unique cell surface composition of tumor cells forms the molecular basis for many targeting and cell-based therapies. Here, we describe the development of novel peptide-based targeting agents for Programmed Death Ligand 1 (PD-L1). Molecular imaging by peptide agents, coupled with therapeutic intervention using the same modality, represents a critical advancement in cancer management. Whole-body PET imaging of PD-L1 expression offers a superior alternative to traditional immunohistochemistry, making PD-L1 radiodiagnostic imaging a highly sought-after modality. PD-L1 targeting modalities developed for clinical imaging to date can be divided into antibodies, protein domains, and small macrocyclic peptides with fewer than 20 amino acids. The latter modalities can address many challenges seen in antibody-based targeting vectors. All potent PD-L1 targeting peptide modalities reported to date rely extensively on non-canonical amino acids (ncAAs). Here, we report a comprehensive structure-activity relationship (SAR) analysis of a family of macrocycles discovered from an Sx2Cx8Cx2 phage-display library composed entirely of natural amino acids (x represents 19 natural amino acids excluding Cys). Using >10,000 variants in “focused” phage-display libraries, we optimized these macrocycles to achieve single-digit-nanomolar potency in protein- and cell-based assays. En route to this optimization, the activity of 216 synthetic macrocycles towards PD-L1 was measured in five distinct assays; two leads have been evaluated by imaging in tumor xenografts in mice, and the X-ray structure of one advanced lead in complex with PD-L1 has been determined at 2.78 Å resolution. This publication demonstrates the development potential of PD-L1-targeting macrocycles that do not require extensive incorporation of ncAAs and the democratization of discovery by mapping the optimization path to single-digit-nanomolar assets for targeted radiopharmaceuticals via canonical phage-display technology.

## Introduction

The unique cell surface composition of tumor cells forms the molecular basis for all antibody (Ab)-based targeting, antibody-drug conjugates, and cell-based therapies (e.g., CAR-T)^1,2^. Originally envisioned as “magic bullets”, directed therapies continue to suffer from many of the same challenges as conventional therapies, including poor uptake, tumor heterogeneity, and resistance^3^. Molecular imaging by peptide agents, coupled with therapeutic intervention using the same modality, represents a critical advancement in cancer management^3^. Designed to both diagnose and treat, targeted radiotherapeutics (TRPs) include the two FDA-approved drugs Lutathera® and Pluvicto®, as well as over 25 peptide-based TRPs in Phase I-III clinical trials. In diagnostic mode, TRPs can predict efficacy and stratify patient populations before administering the therapeutic version of the same modality^4,5^. Diagnostic imaging involves peptides linked to “imaging isotopes” (e.g., positron or gamma emitters for PET or SPECT). When the same peptide delivery vector is linked to “therapeutic isotopes” (i.e., α- or β-emitting isotopes), it enables targeted therapeutic intervention. Genetically encoded technologies are leading tools for the discovery and development of peptide vectors for a broad class of extracellular receptors. This publication describes the application of genetically encoded technology for the development of novel peptide-based targeting agents for Programmed Death Ligand 1 (PD-L1).

Increased PD-L1 expression in a patient’s tumor is associated with a higher likelihood of responding favorably to PD-1:PD-L1 blockade^6,7^. The standard-of-care for PD-L1 status remains tumor tissue immunohistochemistry (IHC) from resected specimens or biopsies^8^. IHC is limited to biopsy samples, which are often fragmented and do not represent the entire tumor or intertumor variability; PD-L1 analysis in biopsy samples can therefore lead to false-negative results and misdiagnosis^7^. A PD-L1 radiodiagnostic PET scan is a highly sought-after modality, especially for metastatic disease, where target expression exhibits distinct heterogeneity across different lesions and over time. Whole-body autoradiography of PD-L1 expression offers a superior alternative to traditional IHC, as it accounts for tumor heterogeneity in PD-L1 expression and can map response over time^9^. Imaging agents that bind to PD-L1 on cancer cells capture this variability and hold promise as improved predictors of treatment outcomes in human imaging trials. Indeed, imaging studies with ^89^Zr-atezolizumab (anti-PD-L1 IgG1), adnectin ^18^F-BMS-986192, and ^89^Zr-nivolumab (anti-PD-1 IgG4) in non-small cell lung cancer (NSCLC), triple-negative breast cancer, and bladder cancer correlated with improved overall treatment outcomes^10,11^. However, other clinical trials have shown poor correlation between treatment outcomes and radiolabeled PD-L1 imaging using antibodies. These include trials on squamous cell carcinoma of the head and neck (SCCHN), where imaging of patients with ^89^Zr-durvalumab (n=13) yielded the best tumor-to-blood pool ratio but showed no correlation with response to durvalumab treatment^12^. In a trial with NSCLC patients (n=12), the uptake of ^89^Zr-pembrolizumab was higher in responding patients than in non-responders, but the difference was not statistically significant^4^.

The non-specific uptake of large IgG molecules in tumors due to enhanced permeability and retention (EPR) effects may limit the effectiveness of PD-L1 imaging with antibodies^13^. EPR-driven uptake may be reduced in small protein domains, and a rich repertoire of mini-domains has been developed to target PD-L1 (see comprehensive review^14^). Examples include BMS-986192, discovered by mRNA display screening^15^, and FN3_hPD-L1_, discovered by yeast display screening^16^ of ∼100 amino acids (aa) compound libraries derived from 10^th^ type III domain of human fibronectin (10Fn3)^15^ (Fig. 1a). Other ligands include nanobodies NM-01^17^, Nb109, APN09, K2 (120 aa), and KN035 (128 aa), derived from phage-display libraries from immunized camels or alpacas^14^; affibodies M1 (59 aa),^18^ PDA (74 aa)^19^, and Z_PD-L1_ (61 aa)^20^ (Fig. 1a). Cysteine-rich protein MODP-1 (47 aa, 6 Cys)^21^ was developed by truncation and optimization of the sequence of native PD-1 ligand (Fig. 1a). A yeast display screen of a computationally designed library yielded a Cys-rich protein, PDL1B1G1 (49 aa, 6 Cys, K_D_=40 nM), which, after affinity maturation in yeast display, yielded PDL1B1G2 with K_D_=0.2 nM (Fig. 1a)^22^. The smallest Cys-rich domain (27 aa, 6 Cys) was recently developed by Chuanliu Wu and co-workers: the authors discovered the lead dmp1 with K_D_=2.14 μM by phage display and optimized it by yeast display to yield the dmp9 lead (Fig. 1a), with K_D_=0.3 nM for binding to purified PD-L1 and EC_50_=6.5 nM for binding to PD-L1^+^ cells^23^.

**Figure 1.**
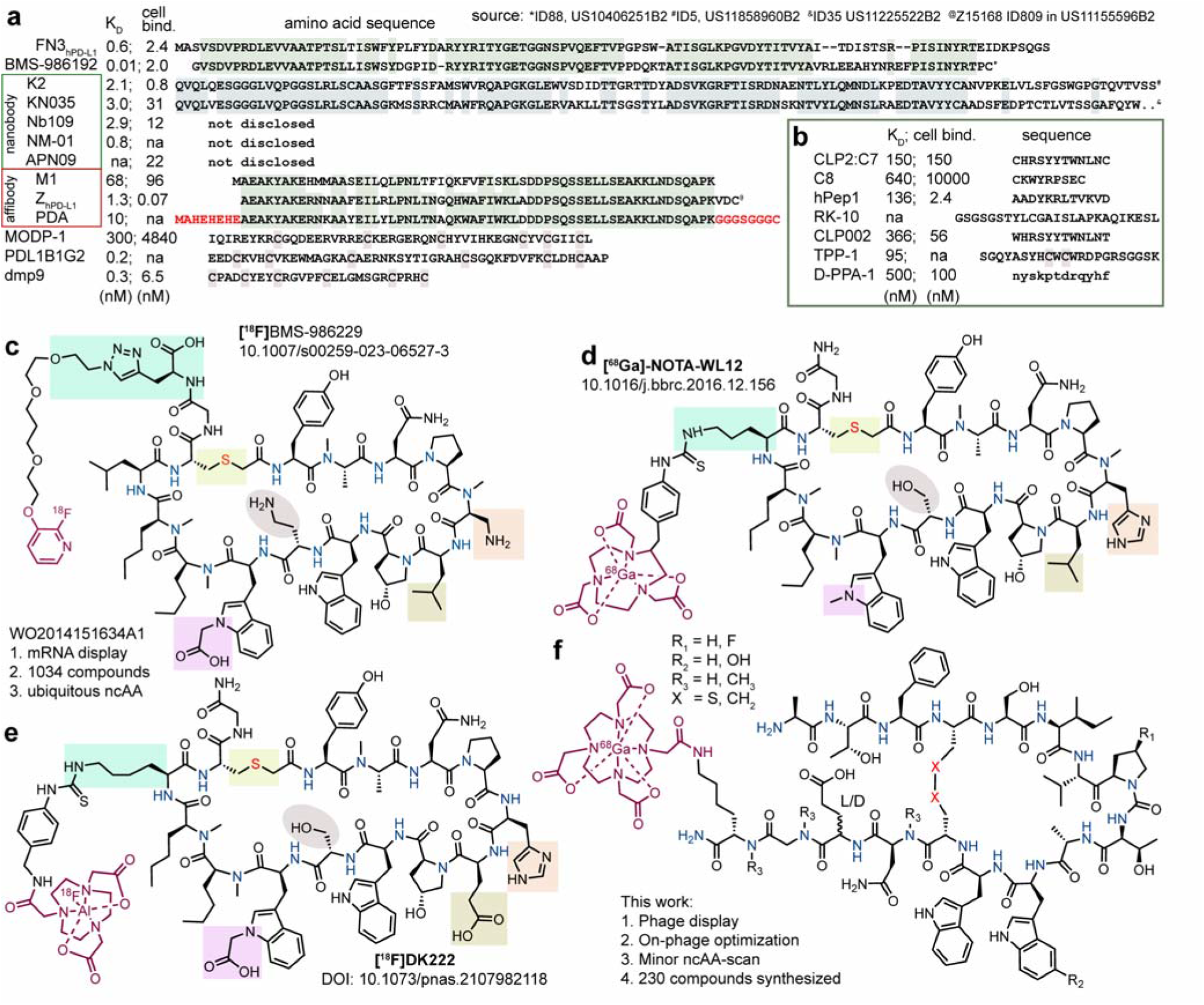
Examples of Targeted Radiopharmaceutical (TRP) macrocyclic peptides developed for PD-L1 imaging. **a**, Sequences of protein domains reported to bind to PD-L1, alongside their reported K_D_ values in binding to PD-L1 and cell-binding activities. Similar sequences are grouped together. Sequences of adnectin BMS-986192, nanobodies K2 and KN035 and Merck affibody Z_PD-L1_ were inferred from the indicated patent sources. **b**, Sequences of linear and disulfide-constrained peptides reported to bind to PD-L1, alongside their reported K_D_ values for binding to purified PD-L1 and cell-binding activities. **c-f**, Macrocyclic peptides that bind to PD-L1 and originate from the same IP portfolio WO2014151634A1; shared and divergent features are color-coded. **f**, The macrocyclic peptide family targeting PD-L1 developed in this work has a distinct topology and amino acid sequence. It contains an aromatic F-W-W triad with a spatial arrangement resembling the Y-W-w’ triad in BMS-family macrocycles, where w’ is an ncAA derivative of W; highlighted are non-canonical amino acids identified to improve the properties of these macrocycles or non-canonical linkers that do not interfere with binding (X=S or CH_2_)

Mini-protein domains targeting PD-L1 show promise in clinical imaging^14^. For example, tumor uptake of ^18^F-BMS-986192 in patients with advanced NSCLC (n=13) correlated with response to anti-PD-1 therapy^11^, and in patients with metastatic melanoma (n=8)^24^, successfully predicted an immune check-point inhibitor treatment-induced reduction in lesion volume. Imaging with nanobody ^99^Tc-mNM01 in NSCLC patients (n=16) correlated with PD-L1 immunohistochemistry results^17^. The smaller size of these domains compared with antibodies allowed PET scans to be performed just one hour after tracer injection, rather than multiple days later, as is required for antibody-based TRPs. This single-visit examination is significantly more convenient for patients, reduces radiation exposure, and enables multiple imaging sessions within a short time frame—particularly critical for patients with metastatic disease who require immediate treatment^24^.

Small peptide-based imaging modalities address many challenges seen in antibody-based TRPs^25^. To date, PD-L1 imaging in humans has been performed with the macrocycles ^18^F-BMS-986229^25^, ^68^Ga-NOTA-WL12^26^, and ^18^F-DK222^27^ (Fig. 1c-e), all three of which originate from the same intellectual-property portfolio (WO2014151634A1), developed via mRNA display by Bristol Myers Squibb (BMS) and referred to in this publication as “BMS-macrocycles”. Other macrocyclic peptides in the BMS-macrocycle family include PCP2^9^, ^68^Ga-DOTA-P6^28^, NJMP ^29^, PG-1^30^, iPD-L1^31^, and pAC65^32^. Up to 50% of the amino acids in these compounds are non-canonical amino acids (ncAAs). There are no reports of cyclic peptides made entirely from natural amino acids with low-nanomolar activity reminiscent of BMS-macrocycles. Several disulfide-constrained peptides exist but lag in performance by 2-3 orders of magnitude. A cyclic 12-mer peptide CLP-2:C7 (CHRSYYTWNLNC, Fig. 1b) was reported to have IC_50_ around 150 nM and similar cell-binding potency^33^. Screening of phage-display libraries identified disulfide-constrained “C8” with K_D_=640 nM (Fig. 1b); the authors observe 60% blocking of Fc-PD-1 binding to CHO PD-L1^+^ cells at 10 μM of C8^34^. A series of linear peptides have also been reported: Examples include hPep1 (12 aa, EC_50_=136 nM), RK-10 (24 aa, K_D_ not reported),^35^ CLP002 (12 aa, K_D_=366 nM), TPP-1 (23 aa, K_D_=95 nM), and the linear 12 aa D-peptide D-PPA-1 with K_D_=500 nM^14^.

A comprehensive survey of the literature suggests that all small peptides or macrocycles outside the BMS family lag in performance compared to BMS-macrocycles. It is not clear whether potent responses exhibited by BMS-macrocycles are dependent on the presence of ncAAs, N-methylation, or a thioether bridge historically employed in mRNA display and recently in phage display^36^. Our report demonstrates that none of these non-canonical features are necessary for low nanomolar binding to PD-L1. In this report, we describe a family of disulfide-constrained peptide macrocycles composed primarily of natural amino acids that achieve potency analogous to that of ncAA-rich macrocycles in the BMS family (Fig. 1f). Optimization driven by phage display yielded multiple leads with EC_50_=1-2 nM in cell-engagement assays without relying on a plurality of ncAAs and complex multicyclic topologies. The outcomes from this publication offer several important observations, including the development potential of macrocycles devoid of ncAAs and the democratization of discovery by mapping the optimization path to single-digit-nanomolar assets via canonical phage-display technology.

## Results

### Optimization and structure-activity relationships using focused phage libraries

We have previously described a series of calibrated phage-display discovery campaigns that identified PD-L1-binding macrocycles starting from >10^10^ diverse peptides displayed on phage^37^. The published screen discovered a family of peptides with Cys-flanked octapeptide W**C**xLVPEAxW**C** motif, and a series of substitutions around the conserved core improved its potency^37^. Here we performed an extensive optimization campaign and built the structure-activity relationship (SAR) using >10,000 variants of the VPxA-macrocycles displayed in five iterations of “focused” phage-display libraries (Fig. 2).

**Figure 2.**
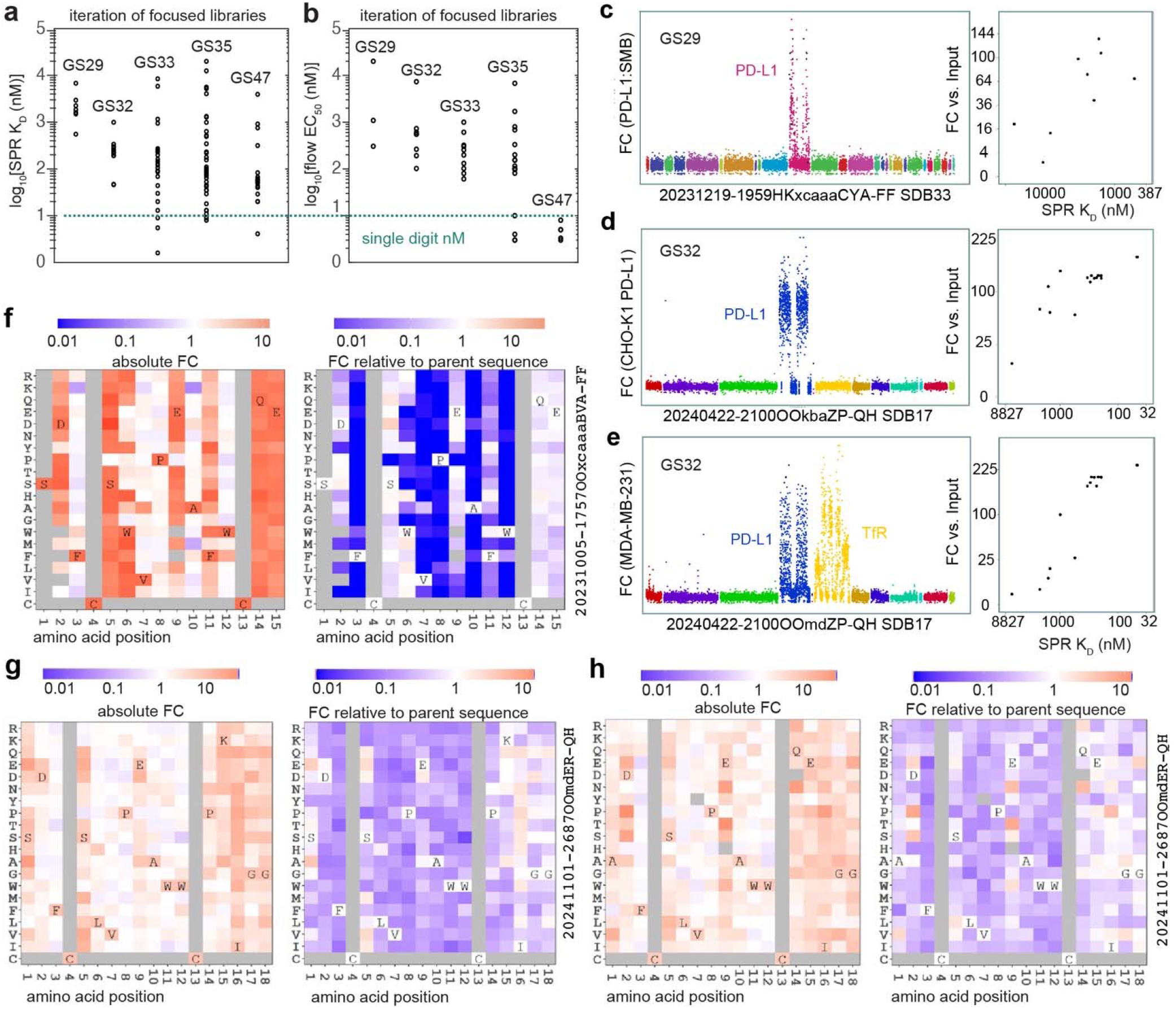
Optimization of PD-L1 binding macrocycles using focused libraries. **a-b,** Five major iterations of focused libraries were designed; peptides nominated from each library were synthesized and measured by SPR (**a**) or flow cytometry (**b**). The plots show the distribution of K_D_ measured by SPR (**a**) and EC_50_ measured by flow-cytometry (**b**) for compounds nominated from these focused libraries. The third-generation library (GS33) contained peptides with validated single-digit-nanomolar K_D_ values (**a**) but only mid-nanomolar EC_50_ values in cell-binding assays (**b**). The fourth and fifth iterations (GS35 and GS47) identified multiple candidates with single-digit-nanomolar EC_50_ values in cell binding assays. **c,** Manhattan plot describing panning of the first-iteration focused library GS29 against purified PD-L1. GS29 contained 1062 sequences nominated from the PD-L1 screen (burgundy) and ∼10,000 other peptides from 21 other discovery campaigns for 21 other targets^37^. Nine peptides from the burgundy segment were synthesized, and their affinities were measured by SPR (left). The left and right plots share a Y-axis, which describes the fold-change (FC) enrichment of each peptide determined by differential enrichment (DE) analysis of NGS data. **d-e**, Two Manhattan plots similar to **c** describe panning of the second-generation focused library GS32 against PD-L1^+^ CHO cells (**d**) and MDA-MB-231 cells (**e**). DE-analysis of NGS data determined the enrichment (FC) of every peptide in GS32 relative to the input; each dot represents a unique peptide sequence. Eleven populations represent 11 families of sequences; the navy-colored family contains PD-L1-binding sequences and positional scans of these sequences, whereas the yellow family contains sequences that bind to the human transferrin receptor (huTfR). The latter sequences bind to human MDA-MB-231 cells but do not bind to CHO cells, which do not express huTfR. Comparison of FC values with K_D_ values measured by SPR shows a better correlation in experiments employing MDA-MB-231 cells with moderate levels of PD-L1 expression than in experiments employing PD-L1^+^ CHO cells with high levels of PD-L1 expression. **f-h,** Heat maps describing the absolute and relative FC values from deep mutational scans (DMS) of three peptides from two focused libraries. **f,** The DMS of the parent sequence SDFCSWVPEAFWCQE (**8191**) measured in GS29 illustrates specific AA changes associated with a loss of peptide affinity (blue squares) or a gain in peptide affinity (warm colors). **g,** The expanded DMS of advanced leads from the PKI-family suggest favourable changes at AA positions 1, 5, 8, 16-18 and no further possibilities for improvement at positions 6-8 and 10-13. **h,** The DMS of the QE-family offers similar information but reveals different positional preferences of this family. Each DMS contains the file name of the NGS dataset from which it originated

We validated the results from focused libraries by measuring the activity of 216 synthetic compounds in four assays (Supplementary Fig. 1). The focused phage-displayed libraries were constructed from SC1966-HF92:GenTitan Hi-Fi 92k pool, 12-50K defined DNA sequences of 147 bp each (GenScript). This DNA, when cloned into phagemid or phage vectors^38^ equipped with silent DNA-barcodes^39^, produced a library of 12-50K peptide sequences (Supplementary Fig. 2). Five focused libraries GS29, GS32, GS33, GS35 and GS47 were produced (Fig. 2a-b) and panned against purified PD-L1 protein, PD-L1^+^ CHO or MDA-MB-231 cells (Fig. 2c-e, Supplementary Fig. 2-7). We calibrated these focused libraries by comparing the fold-change (FC) enrichment observed in the focused library with the affinities of synthetic peptides (Fig. 2c-e, Supplementary Fig. 3-7). Enrichment information provided by the focused libraries made it possible to systematically improve the lead compounds.

The focused libraries contained four classes of sequences. Class 0 comprised sequences not related to PD-L1 selection; these sequences critically offered a “baseline response”^37^ (for example, non-burgundy sequences in Fig. 2c or non-navy sequences in Fig. 2d). Class I were random sequences identified in screening of billion-scale phage-displayed libraries. Class II were near-neighbour derivatives— mutations, deletions, insertions—of the sequences with validated performance (Fig. 2f-h). Class III were sequences that integrated multiple observations in near-neighbour derivatives. Class IV came from affinity maturation libraries (AffMat, Supplementary Fig. 8-9). Sequences nominated from AffMat were re-cloned into focused libraries for re-testing alongside other sequences. An iterative production of five focused libraries (Fig. 2a-b) made it possible to drive the affinity and cell-binding capacity of the optimized peptides. Specifically, K_D_ of the hits progressed from the low-micromolar to the single-digit-nanomolar range after three rounds of optimisation guided by focused libraries: the K_D_ values of three macrocycles from focused library GS33 were in the single-digit-nanomolar range (Fig. 2a). Similarly, the EC_50_ values of these hits in cell-based assay progressed from the micromolar range for early hits to the single-digit-nanomolar range after four rounds of optimisation guided by focused libraries: the EC_50_ of three synthetic macrocycles from focused library GS35 were in single-digit-nanomolar range (Fig. 2b).

A focused library denoted as GS29 contained class 0, I and II sequences. GS29 contained 1062 sequences that originated from PD-L1 screen (burgundy segment in Fig. 2c), and 9,000 peptides from 21 discovery campaigns for 21 other protein targets^37^: the latter sequences were class 0 or “baselines”. An example of a Class I sequence is the SDFCSWVPEAFWCQE (**8191**) discovered in three-round phage-display selection^37^. Sequence **8191** had EC_50_=10 μM in cell-binding assay, and K_D_=2-3 μM for purified PD-L1. GS29 contained 216 Class II sequences located within one Hamming distance (H^1^) of **8191**: each of the 12 underlined amino acids in SDFCSWVPEAFWCQE was systematically changed to 18 other amino acids, excluding Cys. The H^1^ heat-map of this deep mutational scan (DMS) in Fig. 2f summarizes the absolute fold change enrichment factors (FC) and relative FC of these 216 peptides compared to the FC of the parent SDFCSWVPEAFWCQE peptide. In the relative FC-DMS plot, every dark-blue position represents an apparent loss-of-function of the mutant peptide displayed on phage (Fig. 2f). DMS suggested that positions 3, 8, and 10 had an absolute requirement for F, P, and A with a >10-fold drop in FC for any other amino acid at these positions. Positions 7 and 12 appeared to yield only a minor penalty for V7A or W12F substitutions. Changes to positions 1 and 14-15 appeared to offer 10-12 net-neutral possibilities for substitution, whereas mutations in positions 5-6 and 11 offered suggestions for improvement (warm shades on the heatmap, Fig. 2f). Analysis of the affinities of the synthetic peptides confirmed many predictions from DMS and GS29 (Supplementary Fig. 3-4 and “QE-family” below).

The value of the DMS heatmap for SDFCSWVPEAFWCQE (**8191**, Fig. 2f) is determined by the relationship between FC enrichment factors in GS29 and the affinities observed for synthetic peptides. The synthesis of peptides retroactively confirmed a correlation between the fold-change enrichment factors (FC) observed in focused library GS29 and EC_50_, K_D_, and IC_50_ values (Supplementary Fig. 3-4). Similar calibration was performed for library GS32 (Supplementary Fig. 5-7). We noted that under certain conditions, such as panning focused libraries on cells that express a high density of PD-L1, the calibration was “flat”: the ability to discriminate mid-nanomolar from micromolar binders was poor, whereas decreasing the density at which the target was displayed improved calibration (Fig. 2d, 3; Supplementary Fig. 2b, 3b, 4i, 6a-e). A surprising observation was that the format of display—multivalent phage or monovalent phagemid—had only a minor effect (Fig. 3; Supplementary Fig. 6 vs. 7). In conclusion, the use of targets with low display density is generally beneficial, whereas the use of monovalent display offers only modest benefit. Data on the reproducibility of calibrations for libraries GS29 and GS32 versus SPR-K_D_ across different library and target batches and experiments conducted by different users are available in Supplementary Fig. 2-6; expansion of this analysis to FC-EC_50_-K_D_-IC_50_ correlations across multiple libraries and target formats (proteins, cells) is beyond the scope of this report and will be presented in subsequent reports.

**Figure 3.**
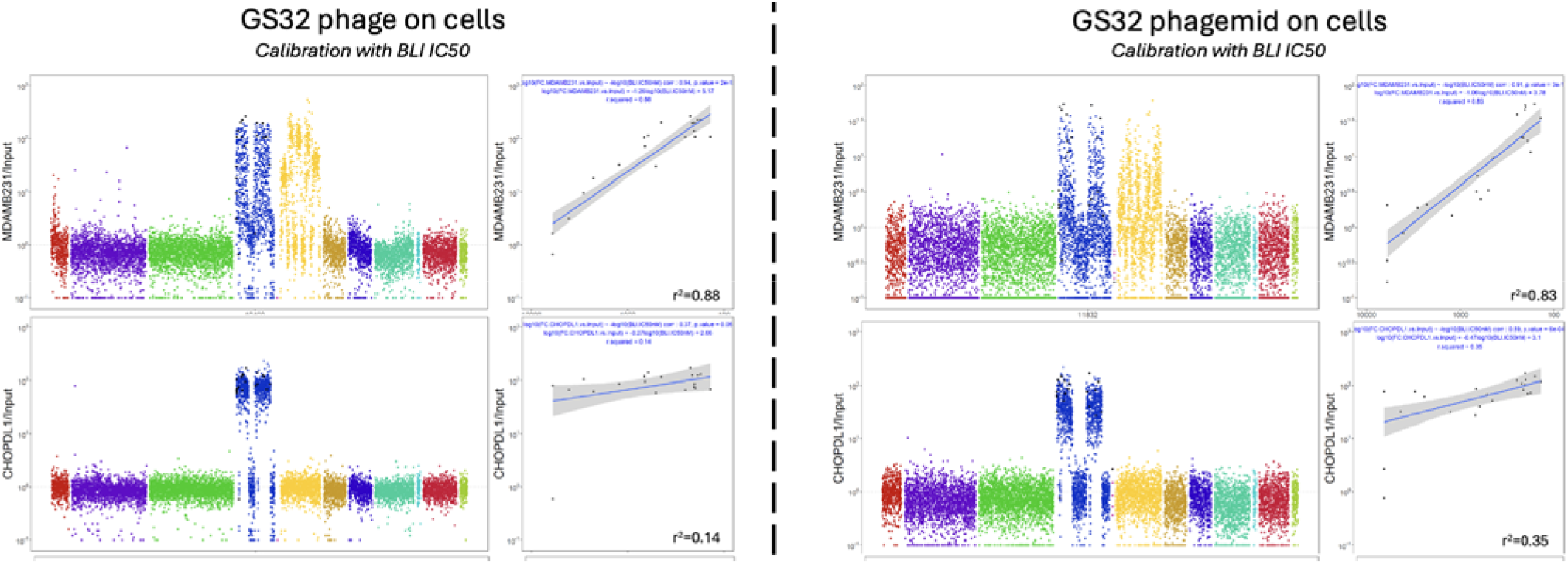
Example of calibration of focused library GS32 in two conditions and two formats. Cell-based assays suggest that panning the MDA-MB-231 cancer cell line, which has a lower density of PD-L1 expression, is a better predictor for IC_50_ than panning PD-L1^+^ CHO cells expressing a high density of the same protein. When target density is high, as in the case of an overexpressing cell line, the change from phage to phagemid appears to offer little to no improvement. In contrast, both phagemid and phage screens offer satisfactory correlation when panning MDA-MB-231 cells that express a modest level of PD-L1. Each unique dot on the Manhattan plot represents a unique peptide sequence. Eleven colors in the Manhattan plot represent 11 families of sequences, of which only one, the navy-colored family, originates from the PD-L1 screen. The ellow family contains peptides that bind to the human transferrin receptor (huTfR). These sequences bind to human MDA-MB-231 cells but do not bind to CHO cells, which do not express huTfR. The experiment in this figure was performed on the same day by the same user. An analysis of reproducibility in experiments conducted by different users, with different libraries and batches of cells, is presented in Supplementary Fig. 6-7.

**Figure 4.**
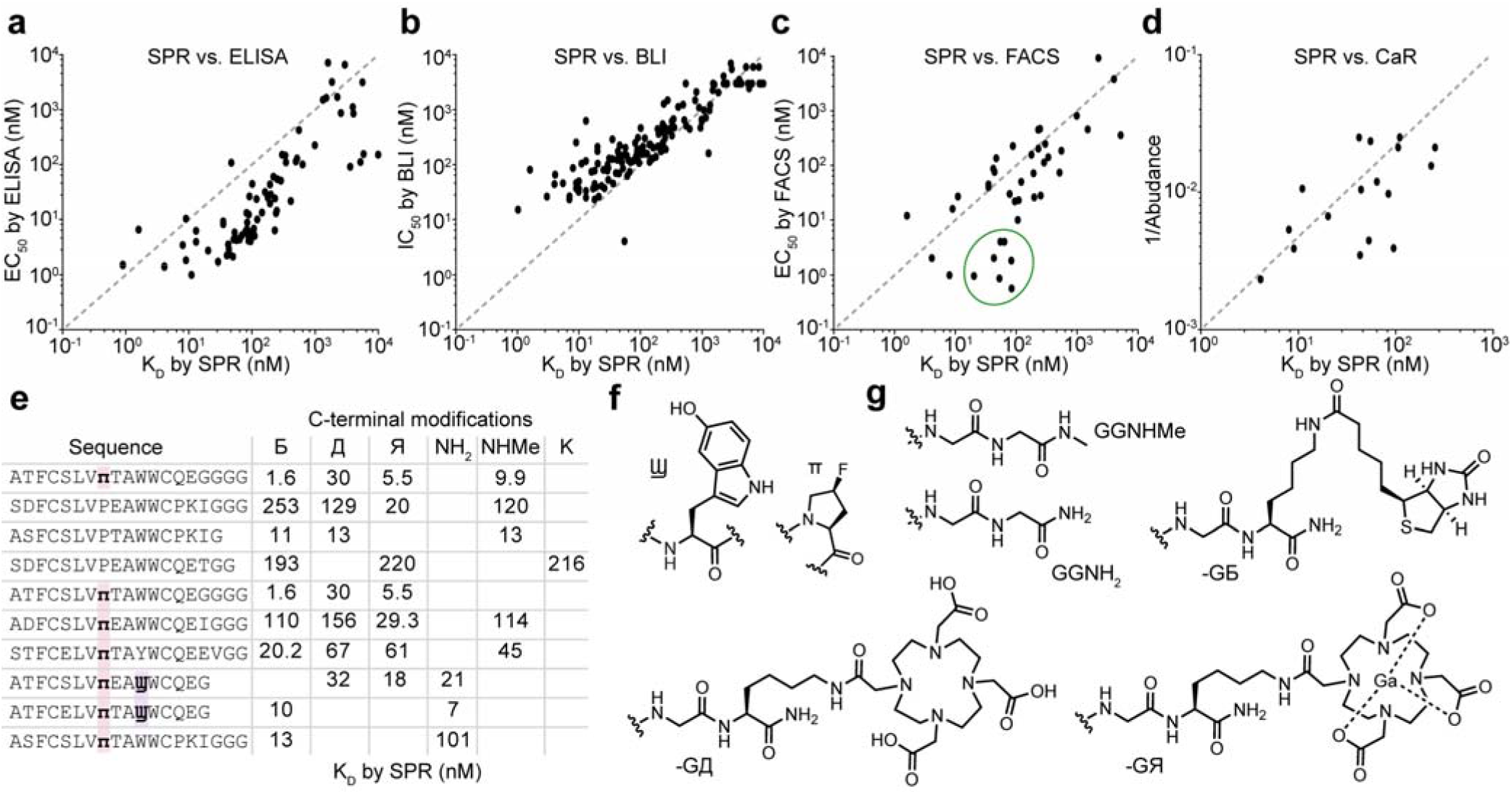
Analysis of the measurements and influence of C-terminal modifications. **a-d,** Correlations between K_D_ values measured by SPR and EC_50_ values measured by ELISA (**a**), IC_50_ values measured by BLI (**b**), EC_50_ values measured in cell-binding assays measured by FACS (**c**), and abundance assays measured by catch-and-release (**d**). The line represents a hypothetical 1:1 correlation. **e,** Summary of 33 K_D_ values measured for ten peptides synthesized with six different C-terminal modifications. **f**, Structures of the unnatural amino acids. **g**, Structures of the C-terminal modifications. Analysis in **e-g** is non-exhaustive because only 33 of 60 plausible combinations were synthesized; empty cells represent compounds that were not synthesized.

**Figure 5.**
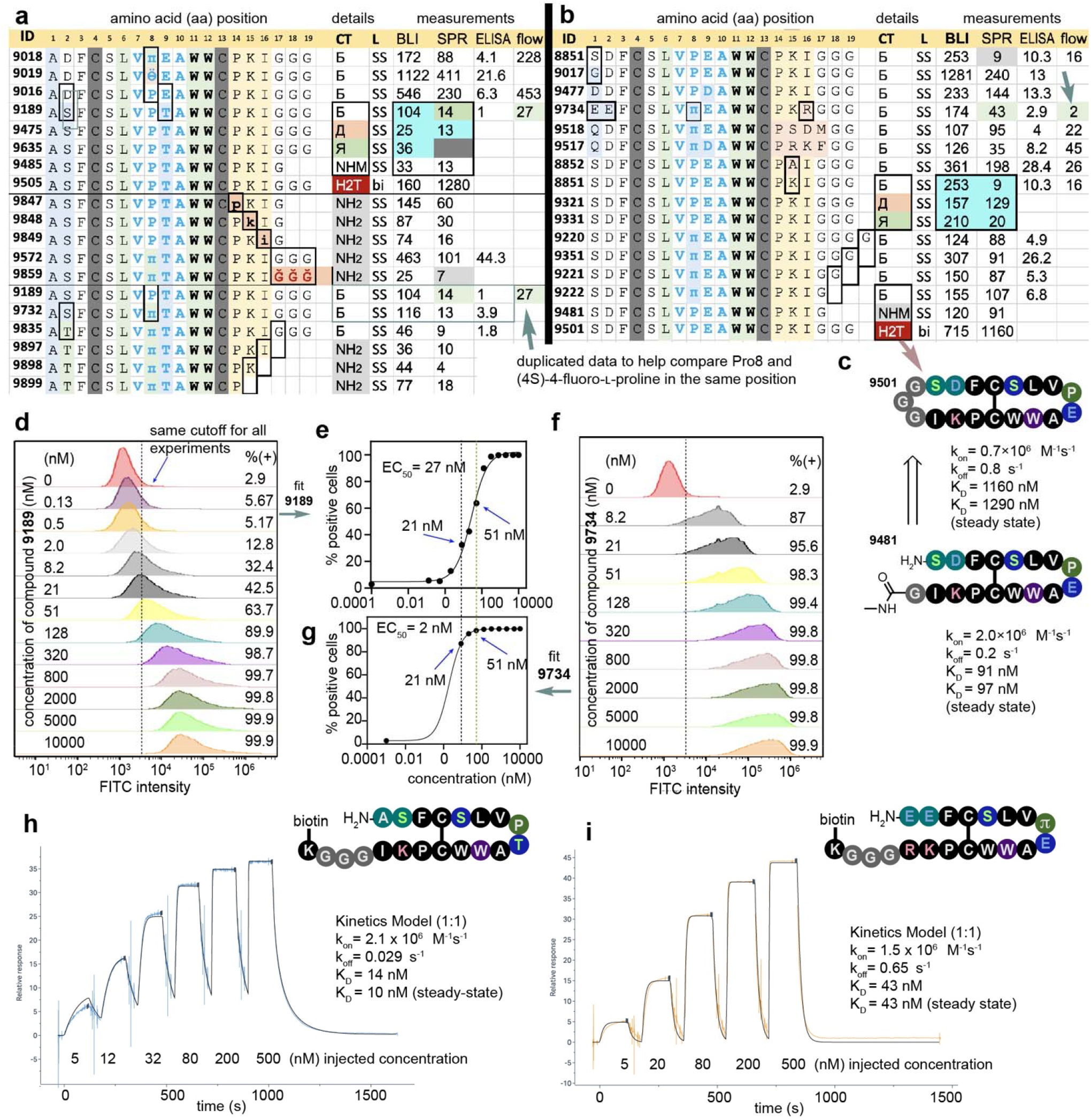
SAR in the PKI family. **a-b,** Summary of the K_D_ measured by SPR, IC_50_ measured by BLI, binding measured by ELISA, and cell-binding measured by flow-cytometry; all values are reported in nanomolar units. Error bars are omitted for clarity and are available in the supporting information. Compounds related by a change in a single amino acid are grouped, and the changes are highlighted by black rectangles. The identities of the C-terminal modifications (CT) are shown in Fig. 4g; the linker designation (L) is SS for disulfide, whereas “bi” refers to a bicycle in which one ring is closed via a disulfide and the other via head-to-tail cyclization, forming an amide between N- and C-termini. **c,** Illustration of bicyclization for compound **9501**, which can be viewed as a derivative of compound **9481**, together with a detailed summary of the on- and off-rates of these compounds. **d,** Visualization of the dose-response titration of biotinylated compound **9189** with PD-L1^+^ CHO cells, yielding an EC_50_ of 27 nM (**e**). **f-g,** The same dose-response titration for compound **9734** estimated an EC_50_ of 2 nM. **h-i**, SPR curves for these compounds.

**Figure 6.**
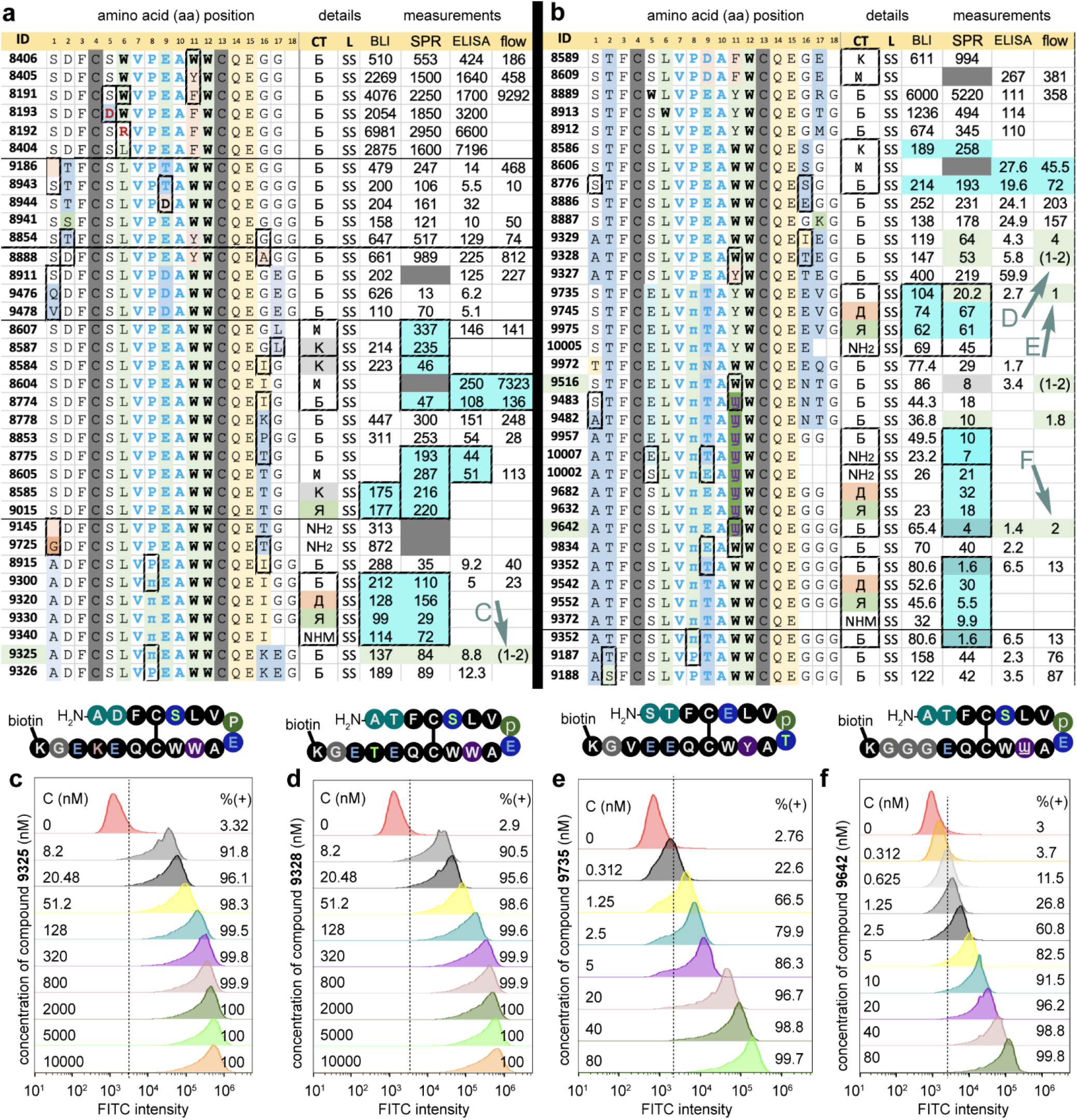
SAR in the QE-family. **a-b**, Tables summarize K_D_ values measured by SPR, IC_50_ values measured by BLI, EC_50_ values measured by ELISA and flow cytometry; all values are reported in nanomolar units. Error bars are omitted for clarity, but representative raw data are available in the Supporting Information. Compounds are grouped to facilitate comparison of amino acid changes (black rectangles). Abbreviations for linkers (L) and C-terminal modification (CT) are defined in Figs. 4 and 5. Measurement values for compounds with identical sequences but different C-terminal modifications (CT) are highlighted in blue. Green highlights denote compounds with EC_50_=1-2 nM as measured by flow cytometry. Note that many of these compounds have only mid-nanomolar K_D_ values as measured by SPR. **c-f**, Flow-cytometry traces for **9325**, **9328**, **9735**, and **9642** (compounds with green arrows in panels **a** and **b**). Compounds **9325**, **9328**, and **9735** with SPR K_D_ values of 84, 53, and 20 nM, respectively, but potent EC_50_ values of 1-2 nM, contain EKE, ETE, and EEV sequences, suggesting a beneficial effect of a negatively charged, hydrophilic, or zwitterionic C-terminus on cell binding.

The calibrated DMS data from GS29 and GS32 libraries identified productive changes at 3-4 distinct positions of the SDFCSWVPEAFWCQE (**8191**), yielding two families of leads with single-to double-digit nanomolar potency. One family based around the sequence **A**DFCS**L**VPEA**W**WCQE**I** is referred to below as the QE-family, and the other, SDFCS**L**VPEA**W**WC**PKI**, is referred to as the PKI-family. These families exhibited not only differences in the underlined positions but also distinct DMS preferences (Fig. 2g-h). Beyond optimization guided by DMS, we constructed four AffMat libraries displayed on a monovalent phagemid vector, three for QE-family and one for PKI-family (Supplementary Fig. 8a-b). We panned a mixture of these four AffMat libraries against three types of cells: (i) CHO cells expressing high levels of PD-L1; (ii) PD-L1-negative CHO cells as a control; and (iii) MDA-MB-231 cells with moderate levels of PD-L1 expression (Supplementary Fig. 8c-e). Calibrated differential enrichment (DE) analysis of the NGS data versus spike enrichment baseline is described in Supplementary Fig. 8f-h. Mapping of the affinity of peptides originating from the analysis of AffMat onto DE-NGS analysis of AffMat is described in Supplementary Fig. 9. These AffMat-derived sequences are bundled in the downstream discussion of SAR; they can be identified as sequences with multiple changes in the N- and C-terminal regions (Supplementary Fig. 9i-k).

**Figure 7.**
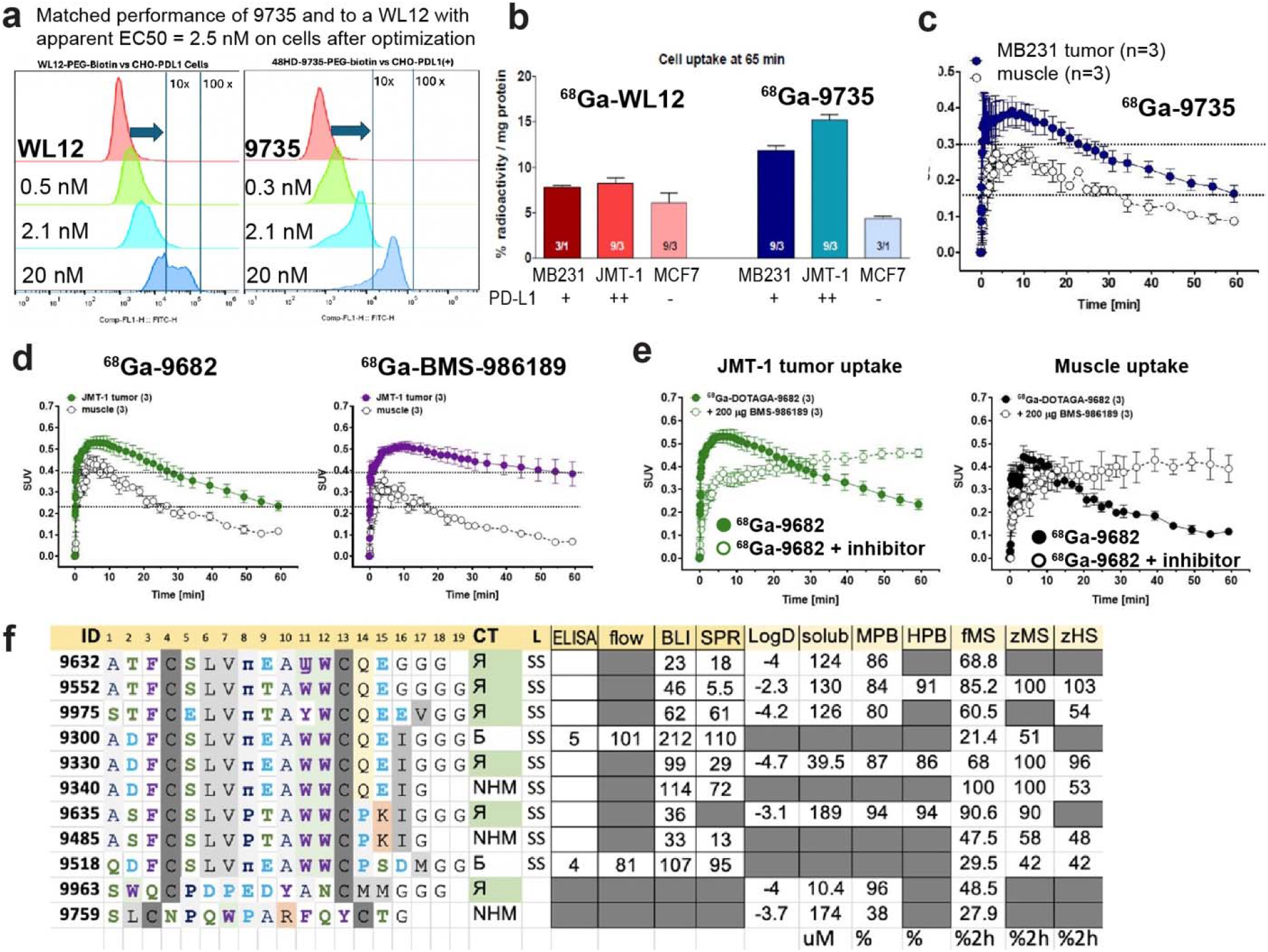
Advanced PD-L1 binding leads. **a,** Dose-response titration of the PD-L1-binding macrocycle **9735** with PD-L1^+^ CHO cells, as measured by flow cytometry; the EC_50_ matches that of **WL-12**^26^ in the same assay. **b,** Cell uptake of ^68^Ga-**9735** *in vitro* exceeds the uptake of ^68^Ga-**WL-12** in the same assay. **c,** *In vivo* PET contrast of ^68^Ga-**9735** shows a suboptimal SUV in the 0.3-0.4 range. **d,** The optimized sequence ^68^Ga-**9682** exhibits an SUV>0.5, similar to that of the best-in-class clinical asset ^68^Ga-**BMS-986189**. Signal deterioration after 30 mins is currently a focus of optimization. **e,** *in vivo* uptake of the PD-L1 binding lead ^68^Ga-**9682** in JMT-1 tumors is specifically inhibited by unlabeled **BMS-986189**, demonstrating that tumor labeling by ^68^Ga-**9682** is specific. **f,** Stability of eleven macrocyclic peptides in fresh mouse serum (fMS), previously frozen mouse serum (zMS), and previously frozen human serum (zHS). Numbers indicate the percentage of intact peptide remaining after 2 h of incubation. Both N- and C-terminal modifications and amino acid sequence influence the stability of the compounds. For seven peptides, we also measured LogD, kinetic solubility (micromolar units), and plasma protein binding in mouse serum (MPB) and human serum (HPB). The latter entries represent the percentage of plasma-bound macrocycle as determined by dialysis. PD-L1 binding data are reproduced from Figs. 5-6 for continuity. For structure of the C-terminal modifications, see Fig 4g.

**Figure 8.**
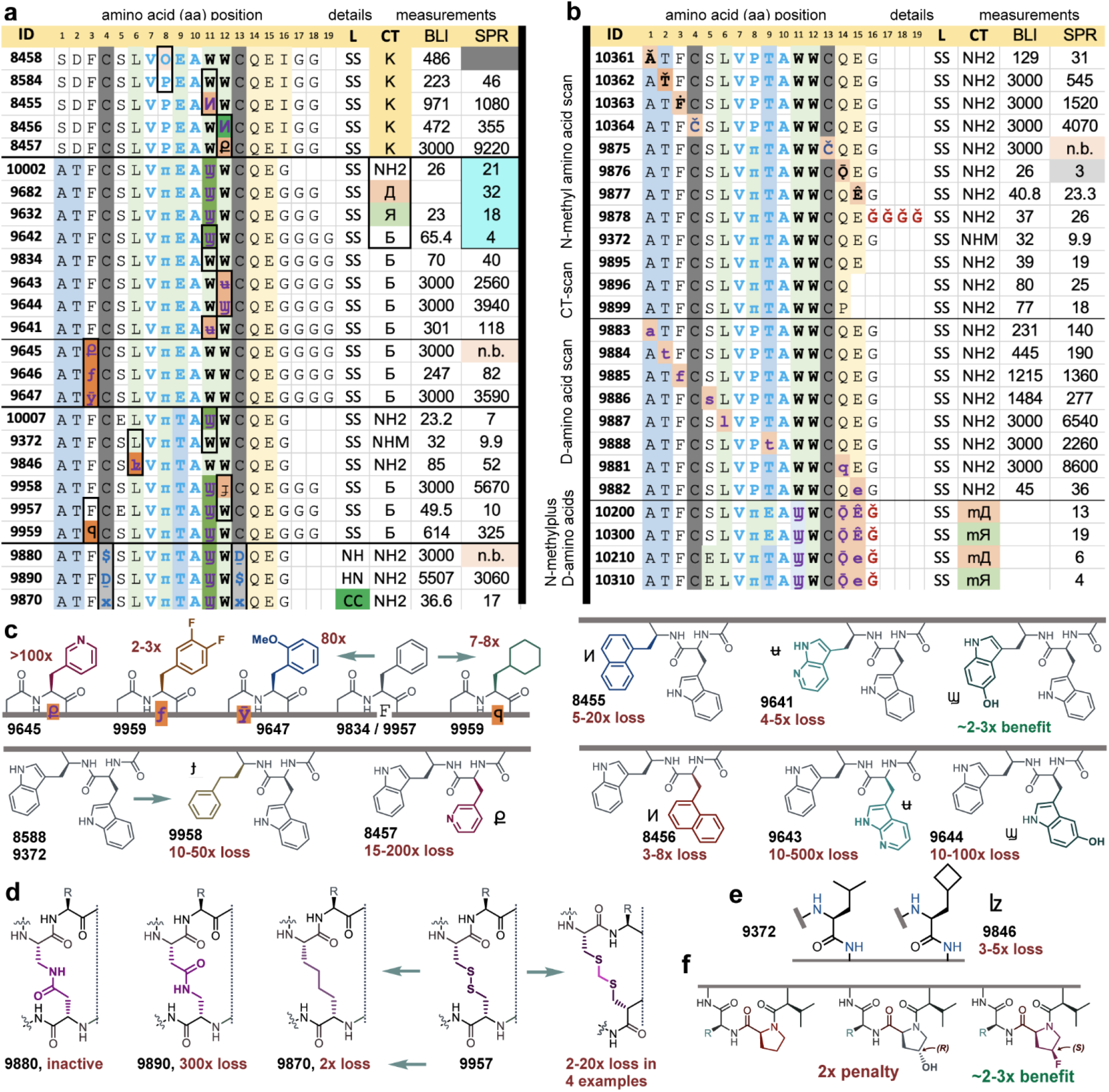
Scanning of lead macrocycles with ncAAs and N-methylated amino acids. **a,** Scanning of positions 3, 6, 8, 11, and 12 with ncAA analogs whose structures are depicted in panels c and e. As in Figs. 6 and 7, peptides with only one position changed are grouped together, and the changes are highlighted. **b,** Systematic replacement of positions by N-methylated amino acids, denoted with a caron (e.g., **c**→č), and D-amino acids, denoted by lowercase letters (e.g., A=L-Ala, whereas a=D-Ala), as well as C-terminal truncations and combinations of multiple ncAAs at the C-terminus. **c**, Structures of ncAA analogs of Phe and Trp. Compound numbers are in bold black, and observed losses in BLI and SPR assays relative to the parental sequences are shown in burgundy. **d**, Structures of non-disulfide bridges and associated compound numbers and losses. **e**, Structure of Leu isostere. **f**, Structures of proline analogs from selected sequences. The full list of proline analogs and compounds is available in Supplementary Fig. 13.

### Validation of synthesized macrocycles: the role of peptide modifications in their activity

During the optimization process, the phage-display driven discovery and measurement of the properties of synthetic peptides were iterated over several cycles. Overall, we synthesized 216 macrocycles with diverse C-terminal modifications (Fig. 4, Supplementary Fig. 1). The affinity of 216 synthetic macro-cycles towards PD-L1 was measured in five distinct assays (Fig. 4a-d). **Assay 1** employed ELISA to measure binding of macrocycles with C-terminal biotin to PD-L1 immobilized on the surface. ELISA yielded EC_50_ values ranging from 1 nM to 20 μM (Fig. 4a). **Assay 2** employed surface plasmon resonance (SPR) to measure the kinetics of binding of soluble peptide macrocycles to PD-L1 immobilized on the surface of a CM5 chip (Supplementary Fig. 1c). **Assay 3** employed a biolayer interferometry (BLI) instrument to measure the ability of soluble peptide macrocycles to block the interactions between surface-immobilized PD-L1 and its native ligand PD-1 present at a constant concentration (Supplementary Fig. 1d). We noted that EC_50_ values from ELISA were consistently lower than K_D_ values measured by SPR, and they tapered off at single-digit-nanomolar EC_50_ values (Fig. 4a). The dynamic range of inhibition assay (IC_50_) was between 50 nM and 6000 nM due to the format of the assay (Fig. 4b). Overall, SPR K_D_ values are considered the standard in this manuscript, but both ELISA and BLI were equally useful. For example, ELISA was also used to show that VPxA-macrocycles, like BMS-macrocycles, bind only to human PD-L1 and not to mouse PD-L1 (Supplementary Fig. 1a). We further used ELISA to show that binding of macrocycles is decreased significantly when N-glycans are removed from PD-L1 by PNGase F whereas treatment by enzymes that trim the terminal glycans has no effect on binding (Supplementary Fig. 1a). BLI inhibition assays have also been used to show that compounds developed in this manuscript bind to the same PD-L1 binding site as WL12 (Supplementary Fig. 1e). **Assay 4** employed flow cytometry and dose-response titration of macrocycles with C-terminal biotin to PD-L1^+^ cells incubated on ice, followed by the readout of the signal with fluorescently labeled streptavidin (Fig. 4c, Supplementary Fig. 1f-i). Measurement of the mean fluorescent intensity (MFI) of the labeled population by flow cytometry (FACS) can be used to estimate the concentration that led to 50% of the maximum MFI value (MC_50_, Supplementary Fig. 1g). We noted the instability of MFI-derived MC_50_ values both for VPxA-macrocycles and BMS-macrocycles because MFI values drifted continuously as concentration increased (Supplementary Fig. 1f-g). As stable estimates of affinity in cell-based assays, we used EC_50_ values, which were estimated as the concentration at which 50% of the cell population exhibited fluorescence above that of the control population stained only with fluorescently labeled streptavidin (Supplementary Fig. 1h-i). The EC_50_ values were 1-2 nM for WL-12 and BMS-986229 (Supplementary Fig. 1h-i), whereas MC_50_ values ranged from 30 nM for WL-12 (Supplementary Fig. 1h-i) to 200-400 nM for BMS-986229 due to the drift in MFI at high concentrations (Supplementary Fig. 1i). In a series of related compounds, MC_50_ correlated with EC_50_ (Supplementary Fig. 5e); however, we generally abandoned MC_50_ in favour of more stable EC_50_ values. The last assay, **Assay 5**, employed catch-and-release native mass spectrometry (CaR-nMS). The CaR method is a rapid, sensitive, label- and immobilization-free assay that enables the simultaneous screening of compound libraries against water-soluble protein targets and the detection and affinity-based ranking of both high- and low-affinity interactions.^40–42^ In CaR-nMS, proteins are incubated with potential ligands, and intact protein-ligand complexes are transferred to the gas phase by electrospray ionization. Complex ions are then isolated and collisionally-activated (heated) to release bound ligands, which are identified by their mass-to-charge ratios (m/z). Only ligands released from isolated complexes are detected; non-binding library components are excluded, facilitating binder identification in complex mixtures. Here, we extend CaR-nMS to peptide-library screening and evaluate its ability to identify protein-binding peptides from a multi-component library (Fig. 4d).

The chemically and functionally diverse collection of macrocycles also provides a reference set for benchmarking emerging technologies for the measurement of peptide-protein interactions. As an example, a small subset of peptides (**9895** and **8585**) was evaluated using the label-free visual interference colour assay (VICA) (Supplementary Fig. 10).

The modest correlation between EC_50_ measured by FACS and K_D_ measured by SPR was not surprising (Fig. 4c), because measurements on purified protein often do not translate directly to receptor binding on cells. A surprising subset of compounds exhibited EC_50_=1-2 nM in FACS, while the SPR K_D_ of the same compounds was in the 10-100 nM range (green circle in Fig. 4c). We examine these compounds in detail in the subsequent sections. A subset of compounds was tested in microscopy-based internalization assays (Supplementary Fig. 11), in dosimetry assays to test the uptake of radioactive peptides *in vitro*, and for homing to tumor xenografts in mouse models (see sections below).

From 216 macrocycles, 53+4 were synthesized as C-terminal amides or N-methyl amides. The activity of these peptides was measured by SPR and BLI inhibition assays, but such peptides were not suitable for ELISA or cell-binding assays. To enable such assays, other peptides contained assay-specific C-terminal modifications: 114 were synthesized with C-terminal Lys(ε-biotin) for ELISA and cell-based measurements, 8+8 were synthesized with Lys(ε-DOTA) either with or without chelated gallium metal (Fig. 4g). A small subset of peptides contained designer modifications for end-to-tail cyclization (Fig. 5c) or dimerization. Retroactive analysis of the synthesized series identified ten sets of compounds that had identical amino acid sequences with three or more divergent modifications at the C-terminus (Fig. 4e). Analysis of this set made it possible to assess the effect of the C-terminal modification on the performance of the peptides in one uniform assay (SPR in Fig. 4e and other assays in Supplementary Fig. 12). In these sets, we observed that addition of Lys(ε-biotin) to the C-terminus resulted in minor attenuation of the K_D_ measured by SPR but, in rare cases, improved binding by a factor of 2-4 compared with the peptide without Lys(ε-biotin). Addition of Lys(ε-DOTA) frequently resulted in a 2-10-fold loss in binding. A change from DOTA to DOTA:Ga (gallium chelated by DOTA) often reverted any damage incurred by the introduction of DOTA (Fig. 4e). These changes are mirrored in other assays and can be tracked, for example, by comparing compounds **8851**, **9189**, **9330**, **9975**, **10300**, **10310** with compounds **9552**, **9331**, **9635**, **9015** in Supplementary Fig. 12. We tested if changes in linker length between the peptide and biotin from Gly_4_ to Gly can influence the outcome of the assays, but we observed no significant differences in potency (see **9220**, **9351**, **9221**, **9222** in Supplementary Fig. 12). The retroactive analysis uncovered a satisfactory tolerance of most peptides in VPxA to C-terminal modifications: most modifications yielded less than an order-of-magnitude change in binding potency. Fine-grained analysis of the data in Supplementary Fig. 12 shows that the C-terminal modifications produce assay-dependent outcomes. One must be mindful that SAR is assay dependent: for example, for many variants SPR-K_D_ values remain unchanged while cell-binding EC_50_ changes. These observations suggest that SAR discussion should strive to employ data from one type of assay and peptides with identical C-termini.

### Summary of SAR in two families. Family 1: PKI-family

In the macrocycles from the PKI-family, substitution of Pro8 (**9016**) with (4S)-4-fluoro--proline (**9018**) improved K_D_ and IC_50_ by 3-fold; conversely, substitution with (4R)-4-fluoro--proline (**9019**) produced a 2-fold loss (Fig. 5a, Supplementary Fig. 13). These observations were made early in the optimization process and suggested that the pucker of the proline ring is important for affinity. In some peptide sequences, C-terminal addition of a chelator (DOTA, **9475**), Lys(ε-biotin) (**9189**), or replacement of a Gly-Gly tail entirely with C-terminal N-methylamide (NHMe, **9485**) had only a minor effect on K_D_ values measured by SPR. The same invariance of K_D_ was observed in **9835** with GGGK-Biotin tail and **9897** with no tail (i.e., C-terminal Ile-amide, Fig. 5a). The invariance of K_D_ was not universal, and in two sequences a change in the C-terminal tail had a nearly 10-fold effect on K_D_: compare the (**9572**, **9732**) and (**8851**, **9321**, **9331**) sets in Fig. 5a-b.

A series of peptides (**9734**, **9517**, **9518**, **9477**, **9017**) at the top of Fig. 5b represents the results of simultaneous optimization of both N- and C-terminal lariats by affinity maturation using phagemid libraries and PD-L1^+^ cells as bait (Supplementary Fig. 8-9). None of the matured peptides exhibited improvement in K_D_ as measured by SPR when compared to the parent peptide N-terminal ASF or ATF and C-terminal PKI. However, macrocycle **9734** exhibited EC_50_ = 2 nM for macrocycle-cell interactions as measured by flow cytometry (Fig. 5f-g). Cell binding exhibited a 10-15-fold improvement compared with parent compound **8851** (Fig. 5b) or **9189** (Fig. 5d-e). The compounds **9734** and **9189** exhibited diametrically opposing behaviour in SPR and flow-cytometry: **9189** was 4-fold more potent in SPR than **9734** (Fig. 5h-i), but **9734** was 10-15-fold more potent than **9189** in binding to PD-L1^+^ CHO cells (Fig. 5d-f). At a similar concentration of 8.2 nM, compound **9189** labeled 32% of the population, whereas compound **9734** labeled 87% of the population (dotted lines in Fig. 5e-g). We attribute this enhancement to the presence of a charged EE:KR pair at the N and C-terminal lariats and confirm this observation in the other family.

The incorporation of D-amino acids in the advanced lead (**9189**) led to a progressively increased penalty for inversion of stereochemistry at Ile16, Lys15 and Pro14 (**9847**, **9848**, **9849**, Fig. 5a, middle). Comparison of **9572**, **9732**, and **9859** showed that replacement of Gly3 with Sar3, where Sar = sarcosine or N-methylglycine, had a favourable effect on K_D_ and yielded one of the most potent compounds in the series. Ironically, the most potent compound with K_D_=4 nM resulted from simple C-terminal truncations up to Lys15, whereas removal of Lys15 incurred a penalty (**9897**, **9898**, **9899** in Fig. 5a, bottom). The importance of Lys15 was further confirmed by a nearly 20-fold penalty to K_D_ upon K15A mutation in the **8851**-**8852** pair (Fig. 5b). A moderate tolerance of PKI-family to the composition of lariat tails prompted us to connect the N- and C-termini to form a head-to-tail macrocycle with an additional disulfide bridge (Fig. 5c and H2T-bi entries **9501** and **9505** in Fig. 5a-b). Both ring closures worsened the K_D_ to ∼1 μM but maintained the binding mode: both bicycles exhibited inhibitory potencies in the mid-to-high-nanomolar range (Fig. 5a-b). Subsequent X-ray structure explained this loss by showing a significant distance between N and C-termini. The sensitivity of the PKI-family to the C-terminal alteration represents both a hurdle and an opportunity in the future optimization of strategies for integration of metal chelators; however, a low single-digit potency of truncated or N-methylated tails offers a promising path for further development.

### QE-family

DMS of the core of the QE-family suggested the importance of W12, tolerance of W11 to substitutions with other aromatic amino acids (AAs), and the possibility of replacing W6 with Leu, Met, or Ile (Fig 2f). Measurement of the properties of synthetic peptides confirmed that W6L substitution has only a modest impact on activity (Fig. 6a). This change was retained in all members of QE-family to avoid plurality of W in the same peptide. The W_11_W_12_ dyad in many peptides was altered to Y_11_W_12_, resulting in a 3-10-fold loss in potency. It is convenient to refer to the first six sequences in Fig. 6a (**8406**-**8404**) as “first-generation parent sequences”, with K_D_ values in the 500-2500 nM range. These sequences emanated from multi-round selection followed by validation in focused library GS29 (Fig. 2c,f). The next five sequences emerged from DMS optimization in focused library GS32 (Fig. 2d-e), giving rise to K_D_ in the 100-250 nM range, as well as improved cell-binding potency. Activity of these sequences is attenuated by altering E, T, D in position 9 and S, T, D in position 2 (Fig. 6a). The remaining 20 sequences (**8607**-**9326**) in Fig. 6a represent systematic alteration of the C-terminus in positions 15-16 as suggested by positional scans in focused libraries and affinity maturation on PD-L1^+^ cells (Supplementary Fig. 8-9). These changes led to progressive additive improvement, giving rise to up to a 20-fold increase in potency compared with the first-generation parent sequences. When C-terminal changes were combined with S1A change at the N-terminus, some compounds exhibited remarkable single digit nanomolar potency in cell-binding assays (e.g., **9325**, Fig. 6a, c), albeit only a modest K_D_ = 84 nM as measured by SPR.

**Figure 9.**
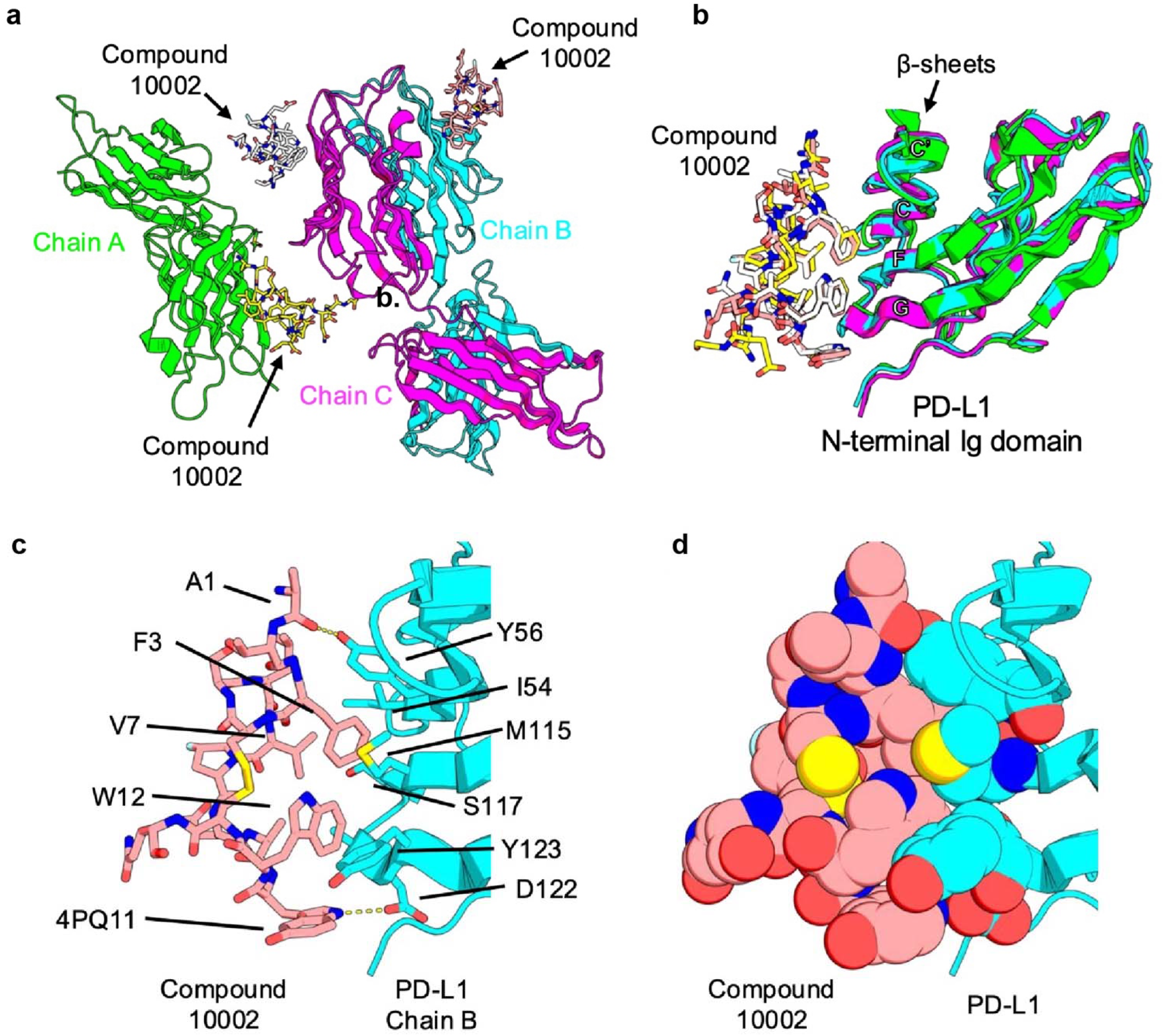
Crystal structure of human PD-L1 in complex with compound **10002**. **a,** Asymmetric unit of the PD-L1:**10002** crystal structure, showing three PD-L1 molecules, each bound to one molecule of compound **10002** at the N-terminal Ig domain. **b,** Superposition of the three N-terminal PD-L1 domains and their corresponding bound **10002** molecules shown in panel **a**, demonstrating the consistency of the binding mode. **c,** Close-up view of the molecular interactions between PD-L1 and compound **10002**, with hydrogen bonds depicted as yellow dashed lines. **d,** Sphere representation of compound **10002** and the interacting PD-L1 residues, highlighting the packing interactions at the binding interface.

Two ncAAs were found favourable in the QE-family. The substitution of Pro9 with (4S)-4-fluoro--proline, denoted as π (**9187**:**9352** pair, Fig. 6b), resulted in a 20-fold improvement in SPR K_D_ and a 5-fold improvement in EC_50_ in cell-based assays. As a result, P9π change was used in 50% of all members of the QE-family. A quarter of the compounds in these series contain a W dyad, where is 5-hydroxy-Trp. This ncAA appeared to be a generally favourable substitution in some compounds or net-neutral in others. Peptides in the PKI- and QE-families exhibited sensitivity to structural changes at the C-terminus. Specifically, the addition of the DOTA chelator often decreased the K_D_ of the peptides. Examples are the SPR K_D_ differences seen in **9735**:**9745**, **9352**:**9542**, **9682**:**9642** pairs (Fig. 6b). However, chelation of Ga by DOTA usually restored the binding affinity. Examples are the SPR K_D_ differences in pairs like **9320**:**9330**, **9552**:**9542**, and **9682**:**9632**. Inhibitory values measured by BLI parallel these observations, but the changes are dampened, possibly due to the limited dynamic range of the BLI assay in the 50-100 nM region.

The N-terminus was important in the QE-family. Both A1 and S1 were associated with higher-affinity compounds compared with the truncated peptide (**9187**, **8943**, **9186**). Fig. 6b describes concerted changes in the N-terminus and positions 5, 9, Trp11, and the C-terminal lariat. The role of C-terminal sequences in the QE-family is illustrated by two observations. First, the C-terminal tail is not essential, and compounds **9642** and **10007**, which lack diversity beyond the QE-sequence, represent local optimization minima with single-digit SPR K_D_ and low-double-digit cell-binding potency. A second observation is that compounds with acidic residues in the C-terminal tail exhibit outstanding cell binding potency as low as 1 nM, and yet they exhibit only modest SPR K_D._ Fig. 6c-f show flow-cytometry dose-response titration curves for all compounds. The enhanced cell binding potency appears to be correlated with the introduction of specific charged residues in the C-terminal region, even though these residues show no improvement in K_D_ as measured by SPR.

The additional charges in N and C-terminal lariats improve interactions with the PD-L1 receptor embedded into the glycocalyx of live cells but have no effect on interactions with standalone purified PD-L1 receptor. As of now, we do not understand the mechanistic origin of the divergence between K_D_ measured on pure protein and cell-based EC_50_ values. The magnitude of this divergence is substantial and should be further investigated because it illustrates that the cell-binding benefits of peptide targeting vectors are not always related to the potency measured on purified proteins. The best SPR K_D_ values of the leads in the QE-family matched the K_D_ values of many BMS-family macrocycles, which exhibit single-digit to low-double-digit nanomolar K_D_ values as determined by SPR using purified PD-L1^14^; however, this family has not yet reached the potency of the best-in-class BMS-986229. A biotinylated variant of BMS-986229 exhibited K_D_=30 pM in SPR (Supplementary Fig. 1c); interestingly the same molecules exhibited EC_50_=3 nM in cell-based assays (Supplementary Fig. 1i). Similarly, disulfide-rich peptide dmp10 reported by Chuanliu Wu and co-workers has K_D_=0.3 nM in binding to purified PD-L1 but EC_50_=6.5 nM in binding to PD-L1^+^ cells^23^. While their performance in binding to purified PD-L1 does not match that of these comparators, the cell-binding performance of QE-family peptides matches that of WL12 (Supplementary Fig. 1f-h), dmp10 (see Figure 3 in the original publication^23^), and a biotinylated variant of BMS-986229 (Supplementary Fig. 1i). The recurrent observation of potent cell-binders in the QE- and PKI-families with performance matched to best-in-class PD-L1 binders confirms the ability of AffMat selection on PD-L1^+^ cells to identify potent cell binders. Inspired by these observations, we tested two peptides: **9682**, a derivative of **9642** (Fig. 6f) and **9745**, a derivative of **9735** (Fig. 6e), in PET imaging in mouse tumor xenografts.

### Testing of PD-L1 Targeting Macrocycles *in vitro* and *in vivo* using PET

We evaluated peptides using 3-4-month-old female NIH-III Nude mice (Charles River Laboratories, Saint-Constant, QC, Canada) with tumor xenografts established by subcutaneous inoculation of either 5x10^6^ JMT-1 or MDA-MB-231 cells. Preliminary *in vivo* PET imaging data were collected for the two lead compounds. The first generation lead **9735** conjugated with a DOTA chelator was radiolabeled with ^68^Ga to yield **^68^Ga-DOTA-9735.** This compound exhibited a significantly higher *in vitro* cell uptake into both JMT-1 and MDA-MB-231 cells (15.2 ± 0.6 and 11.8 ± 0.5 % radioactivity/mg protein; both n=9/3) versus the clinically tested radiotracer **^68^Ga-DOTA-WL-12** (8.2 ± 0.6 [n=3/1] and 7.8 ± 0.2 % radioactivity/mg protein; n=9/3; Fig. 7b), amounting to ∼85% higher uptake in JMT-1 cells and ∼50% more uptake in MDA-MB231 cells. However*, in vivo*, **^68^Ga-DOTA-9735** exhibited only moderate MDA-MB-231 tumor uptake, with substantial washout over time as seen in the time-activity curves (Fig. 7c). Mean SUV values amounted to 0.39 ± 0.04 after ∼10 min dropping to 0.16 ± 0.02 (n=3) after 60 min post injection. Although ^68^Ga-labeled **BMS-986189** has previously been used for PET imaging in MDA-MB-231 xenografts, we observed higher tumor uptake in JMT-1 tumors, with almost no washout: mean SUV values were 0.51 ± 0.02 after 10 min and 0.38 ± 0.06 (n=3) after 60 min post-injection (Fig. 7d). In the same batch of animals, we also tested **^68^Ga-DOTA-9682**, which resulted in a similar initial JMT-1 tumor uptake followed by some washout over time: mean SUV values were 0.53 ± 0.03 after 10 min and 0.24 ± 0.02 (n=3) after 60 min post-injection (Fig. 7d). Specificity of target binding was confirmed by *in vivo* blocking using unlabeled **BMS-986189** (8 mg/kg iv; Fig. 7e) resulting in 45% decrease of JMT-1 tumor uptake at 10 min post injection. However, after 60 min this was reversed and radiotracer tumor uptake was increased by 133%. The same was observed in muscle tissue. This finding was in line with the observations that these compounds share the same PD-L1 binding site (Supplementary Fig. 1e).

Despite this success, we noted substantial tumor tissue washout of ∼55% of **^68^Ga-DOTA-9682** from JMT-1 tumor tissue. In contrast, **^68^Ga-DOTA-BMS-986189** showed only ∼25% washout from the tumor tissue (Fig. 7d). This difference in pharmacokinetic profile may be attributed to (i) potential degradation of **^68^Ga-DOTA-9682** *in vitro* and (ii) the suboptimal off-rate (k_off_) of **9682** compared with **BMS-986189**. To test the role of degradation in serum *in vitro*, we measured the stability of **9632 (^nat^Ga-DOTA-9682)** and observed that only 69% of **9632** remains intact after 2 h of incubation in fresh mouse serum (fMS, Fig. 7f). In the same assays, six other peptides yielded between 20-100% of intact peptides after 2 h of incubation (Fig. 7f). For example, PKI-macrocycles **9635** and QE-macrocycle **9340** exhibited no degradation after 2 h of incubation, illustrating that stability in serum can be achieved even if the peptide is composed of natural amino acids and has a disulfide bridge. Significant changes in the degradation rates of QE-macrocycles **9300**, **9330**, and **9340**, which have identical amino acid sequences and different C-terminal modifications, illustrated a critical role of the C-terminus in this process. The most stable compound, **9340**, lacks a tri-glycine linker. In the PKI-family, comparison of compounds **9485** and **9635** shows the opposite trend, in which removal of the tri-Gly linker from the C-terminus leads to increased degradation (Fig. 7f). This exploration of a set of 11 macrocycles did not definitively identify the SAR for degradation, but the results suggested that N-methyl-scans, D-scans, linker scans, and local ncAA scans in the vicinity of the termini might be beneficial for improving the stability of these peptides. This stability study further highlighted a substantial serum stability of disulfide constrained macrocycles based 100% on natural AA (**9635**) or with only one conformation-constraining ncAA (e.g., **9330** and **9340** have (4S)-4-fluoro--proline). As a result, the hits emanating from phage-display campaigns present a great starting point for fine tuning of pharmacokinetic properties.

Measurement of LogD, plasma binding, and solubility of selected macrocycles did not uncover any obvious liabilities. The relatively high hydrophilicity of the peptide might be one of the contributing factors to rapid clearance. However, 80-90% of the peptides were bound to mouse plasma (MPB, Fig. 7f), and they exhibited similar human plasma-binding properties (HPB, Fig. 7f). The combination of high hydrophilicity and relatively high plasma binding for QE-family of PD-L1 binding macrocycles presents an interesting combination for downstream optimization of *in vivo* targeting. Two substitutions, E9T and 5-hydroxyTrp to Trp at position 11, as well as an extra glycine, changed LogD from -4 to -2.3 (**9632** and **9552** in Fig. 7f), illustrating that LogD in the QE-family can be fine-tuned without affecting other parameters.

### SAR of modification with non-canonical amino acids

DMS and follow-up synthesis confirmed that the substitution at position 11 from the parent phenylalanine (**8404**) to tryptophan (**8587**) led to an improvement, and this position can tolerate tyrosine and 5-hydroxy-Trp. However, other ncAA replacements in this position were detrimental to binding. Changing Trp11 in peptide **9834** to 7-AzoTryptophan (**9643**) or 1-naphthyl-alanine (**8455**) resulted in a 3-5-fold penalty (Fig. 8a,c). All tested ncAA changes to Trp12 were detrimental; in peptide **9834,** changing Trp12 to 5-HydroxyTryptophan (**9643**) and 7-AzoTryptophan (**9644**) resulted in up to 500-fold loss of activity (Fig. 8a,c). Similarly detrimental were changes of Trp12 in **8774** to 1-naphthyl (**8456**) or 3-pyridyl (**8457**). Fig. 8a,c shows that even subtle steric or electronic changes to the side chain of Phe4 were detrimental to binding. The sensitivity to ncAA substitution at positions 12 and 4 aligned with the DMS observations (Fig. 2f, g, h): positions 12 and 4 were exquisitely specific for Trp and Phe, with no tolerance for any other canonical amino acids.

Hybrids of L and D-amino acids are known to be stable to proteolytic degradation and the introduction of even a single D-amino acid can protect against degradation in the vicinity of that ncAA.^20, 21^ To identify permissive positions for D-amino acids, we replaced each amino acid in the ATFCSLVPTAWWCQEG peptide with its D-amino-acid counterpart. This D-scan identified a strongly conserved core spanning amino acids 1-13. The surprising outcome was the sensitivity of terminal Ala1 and Gln13 to the inversion of stereochemistry. In contrast to dramatic loss of activity upon D-inversion of Gln13, there was tolerance of neighbouring Glu14 to the D-inversion. This observation aligns with only a minor loss of activity upon truncation of Glu14 (**9895**, **9896**, Fig. 8b). Inversion of Leu6 resulted in the most profound loss of binding. Even a seemingly minor replacement of Leu6 to cyclobutyl alanine resulted in a 5-fold reduction in binding (compare **9846** and **9372**, Fig. 8b,e). These observations suggest that Leu6 is both a problem and an opportunity: replacement of Leu6 with an ncAA is not trivial, but the role of this residue is profound; hence, there may be an ncAA replacement at Leu6 that may result in a substantially enhanced binding.

An N-methyl-amino-acid scan (Fig. 8b) showed that every amino acid in the C-terminal tail starting from Gln13 can be replaced with its N-methyl analog with no observable loss of activity (**9876-9878**, Fig. 8b). In contrast, N-methylation of any Cys completely abolished the activity (**9875**, **10364**). While N-methylation in the N-terminal lariat was often detrimental (**10362**, **10363)**, the penalty due to N-methylation of Ala1 appeared to be minor (**10361**, Fig. 8b). We sought to identify simple chemical modifications to the N-terminus to reduce exopeptidase degradation. However, modification of the N-terminus with N-acetyl (**9811**), pyroglutamate (**9815**) or acylamide (**9816**), N-tosyl, Ser-derived oxaloyl and hydrazides derived from this oxaloyl were detrimental to affinity (Supplementary Fig. 14). An attractive solution to N-terminal degradation would be ligation of the DOTA chelator to the N-terminus, reminiscent of N-terminal ligation of chelators in Lutathera®. We tested whether N-terminal modification of ∼1000 peptides in a focused library could identify sequences that tolerate N-terminal modification by DOTA chelators. All tested sequences exhibited reduced enrichment upon N-terminal modification (Supplementary Fig. 15a). We confirmed with synthetic compounds that placing DOTA (Supplementary Fig. 15b) or biotin (Supplementary Fig. 15c) at the N-terminus indeed led to a 3-20-fold penalty in activity. Decreased PD-L1 binding upon stereoinversion of Ala1, the minor penalty upon N-methylation, and the larger penalties upon acetylation, alkylation, and ligation of chelators suggest a critical role of the N-terminal residues in recognition of PD-L1.

We observed sensitivity of the hits to changes in the disulfide bridge. We performed extensive testing of chemically modified peptides in focused libraries. Modification of focused libraries by dichloroacetone, meta-dibromoxylene, and other cross-linkers reproducibly resulted in loss of enrichment in all ring-expanded sequences (Supplementary Fig. 16b). A one-carbon insertion into the S-S bridge led to a 3-10-fold loss of potency (Fig. 8d, Supplementary Fig. 16b). Its replacement by amide-bond-based bridges completely abolished activity (Fig. 8d). The most productive change to the disulfide bridge was its replacement by an all-carbon bridge (Fig. 8d), which resulted in only a minor penalty, albeit only in one compound tested so far. Removal of oxidation-prone sulfur atoms from the preclinical lead compounds is very promising because it can improve radiostability.

We made several attempts to change the topology of the hit sequences from monocyclic to bicyclic. Simple linking of the N- and C-termini could, in theory, increase potency and/or reduce proteolytic degradation, but instead this cyclization led to a severe decrease in binding (Fig. 5c). Inspired by PCSK9 inhibitor enlicitide (MK-0616) from Merck^25, 26^, we attempted to link side chain residues that were predicted to be in spatial proximity by molecular modeling studies, but the resulting bicycle incurred a 20x loss in potency (Supplementary Fig. 17). Inspired by Chuanliu Wu and co-workers,^23^ we integrated a pair of CxC motifs at the N- and C-terminal lariats in place of Cys (Supplementary Fig. 18). However, after testing of hundreds of variants in focused libraries, most were completely inactive and indistinguishable from the random baseline (Supplementary Fig. 16b-d). Focused library studies suggested that tetracysteine peptide GCSDFCSLVPEAWWCPKIC could be optimized to yield modestly effective binders (Supplementary Fig. 18e-f), but this direction was not further pursued in this study. The difficulties incurred in topological reshaping in the absence of structural information prompted us to pursue X-ray crystallography. Further optimization of bicyclic structures will be possible using the X-ray structure of the complex (Fig. 9).

Building on these observations, we designed compounds **10210** and **10310** to combine N-methyl-Gln13 with D-Glu14 and N-methylglycine (sarcosine) in the regions separating the macrocycle from the DOTA chelator (Fig. 8b, bottom). Similar compounds **10200** and **10300** combine N-methyl-Gln13, N-methylGlu14 and N-methyl-Gly15 (sarcosine) in the regions separating the macrocycle from the N-methyl-Lys16 attachment of the DOTA chelator. The SPR K_D_ of compounds containing this D- and poly-N-methylated C-terminal tail was driven to the lowest single-digit-nanomolar value. We scheduled these peptides for testing in PET imaging in mouse xenografts *in vivo,* and results of these investigations will be reported in subsequent publications.

### Preliminary investigation of the binding mode using X-ray crystallography

We made multiple attempts to solve X-ray structures of QE- and PKI-family macrocycles, but we were unable to repurpose any reported crystallization conditions to obtain crystals of sufficient quality. In the late stages of preparation of this manuscript, prolonged screening of conditions yielded the first X-ray structure of compound **10002** in complex with human PD-L1 protein at 2.78 Å resolution (Fig. 9a). A detailed structural analysis will be presented in subsequent reports; however, preliminary analysis indicates that compound **10002** adopts an extended macrocyclic conformation and binds along the solvent-exposed face of the PD-L1 Ig domain, interacting with the G, F, C, and C′ β-strands (Fig. 9b). The binding interface is stabilized by two key hydrogen-bonding interactions: one between the A1 carbonyl oxygen of compound **10002** and the Y56 hydroxyl group of PD-L1, and a second between the 4PQ11 indole N–H of compound **10002** and a D122 carboxylate oxygen of PD-L1 (Fig. 9c). In addition to these hydrogen bonds, the complex is further stabilized by extensive van der Waals interactions involving compound **10002** residues F3, V7, 4PQ11, and W12, which pack against PD-L1 residues I54, Y56, M115, S117, and Y123 (Fig 9d). The observations from the structure shed light on multiple SAR trends observed in the previous sections. The structure explains the reasons for the failure of multicyclic and head-to-tail constrained designs and paves the way for structure-guided optimization of these macrocycles toward picomolar activity, improved stability, and other performance characteristics required for first-in-human evaluation of this novel family of macrocycles.

## Discussion

Natural peptide hormones are common sources for therapeutic peptides^43^ and imaging probes^44^. The approved drug Lutathera® is a derivative of the somatostatin hormone. Other receptors with natural peptide ligands, such as bombesin^45^, oxytocin^46^, gastrin^47^, ghrelin^48,49^, vasoactive intestinal peptide^50,51^, melanocortin 1^52–54^, neuropeptide Y^55,56^, neurotensin^57^, and glucagon-like peptide-1 (GLP-1)^58,59^, are currently being investigated in preclinical studies and clinical trials. However, not every extracellular target has a natural peptide ligand. Many receptors lack a natural peptide counterpart; instead, their natural ligands are large proteins (e.g., growth factors) or glycans. For such receptors, targeting vectors must be discovered *de novo*. High-throughput screening (HTS) has successfully discovered ligands for TRP targets that are “druggable” by traditional small molecules; examples include the clinically used FAP-binding diagnostics ^68^Ga-FAPI and ^18^F-FAPI, which are based on the small molecule FAPI-04^60,61^. However, addressing “non-druggable targets” with binding interfaces that span large surface areas is challenging for HTS and small molecules^62^. The contemporary approach to undruggable targets involves *de novo* selection of peptide macrocycles originating from billion-scale phage-, yeast-, bacterial-, and mRNA-display libraries. Specifically, mRNA display is a powerful method for the discovery of peptide macrocycles, including Merck’s first-in-class oral PCSK9 inhibitor MK-0616 (Phase 3 trial)^63,64^ and FDA-approved Zilucoplan® by UCB Pharma.^65^ In the TRP field, the PD-L1-binding macrocycle BMS-986229 successfully completed pilot first-in-human imaging of PD-L1 in esophagogastric cancer (NCT04161781)^25,66^. While many receptors have been targeted by only a few modalities, PD-L1 is an example of a target for which almost every modality has been developed, including small molecules^67,68^, peptide macrocycles, mini-protein domains (Fig. 1), and even aptamers^69^. As a result, PD-L1 represents an important case for comparison among these modalities.

Iterative optimization of the micromolar hit SDFCSWVPEAFWCQE (**8191**) in focused phage-display libraries, combined with fine-tuning of sequences with ncAAs, yielded promising single-digit-nanomolar leads in which C-terminal degradation of the peptide sequence should be suppressed. Liabilities associated with the N-terminus and disulfide bridge remain to be addressed; however, both X-ray analysis and SAR identified productive avenues for this optimization. Future exploration should combine an all-carbon-bridge peptide with D-amino acids and N-methyl amino acids in the C-terminal lariats and test a small library of N-terminal modifications to identify those that enhance binding while diminishing N-terminal degradation.

Throughout this manuscript, we build on the benefits of cell-based panning of phage libraries. Identification of single-digit-nanomolar binders to PD-L1^+^ cells was driven not by optimization of K_D_ in protein-binding assays, but by selection of phage-displayed libraries against PD-L1^+^ cells. A fundamental limitation of mRNA display is its incompatibility with cell-based and *in vivo* screening due to degradation of the RNA message. The phage-display platform is ideally suited for screening on cells *in vitro*^70–75^ and *in vivo*^76–78^, thus addressing a fundamental limitation of mRNA display. Stored inside the M13 virion, the DNA message is protected from degradation *in vivo*^76–79^. Building on this capacity, it should also be possible to map the interactions of multiple PD-L1-binding macrocycles with tumors and normal organs *in vivo (publication in preparation)*. Simultaneous *in vivo* profiling of 50-100 PD-L1-binding macrocycles decoded by next-generation sequencing could offer a valuable addition to TRP development. This publication offers a path towards the development of a new class of PD-L1-binding macrocycles; it also provides an important calibration dataset for further development of the phage-display platform and its application to the discovery of potent peptide-based vectors for TRPs and their optimization using purified proteins *in vitro*, cell-based assays, and potentially, *in vivo*.

## Acknowledgements

This work was supported by Canadian Institutes of Health Research funding (to R.D. and F.W.). J.Walker. acknowledges a Mitacs postdoctoral fellowship, supported by Alberta Innovates AICE-Concepts grant 232404986

## Author Contributions

R.D., conception, design, and writing; V.A., conception, design, and medicinal chemistry; M.L.M., data organization; G.B., computational and bioinformatic analyses; T.B.K.C. and M.K., biophysical measurements; W.K., H.I., K.M., G.S., J.Wang, R.M., and Z.O., panning; R.M. and Z.O., design of panning procedures; J.Walker, D.Y., and M.T., data integration; A.Z., CaR-nMS experiments; Y.C., peptide synthesis; J.Woodfield, *in vitro* radiotracer uptake studies; A.D., peptide medicinal chemistry and synthesis; M.W. and C.B., PET imaging; J.S.K., supervision and design of CaR-nMS studies; H.W.K. and T.K.S., X-ray crystallography; T.M., VICA; F.W., supervision of PET imaging.

## Data Availability

The authors declare that the data supporting the findings of this study are available within the paper and its Supplementary Information file. Should any raw data files be needed in another format they are available from the corresponding author upon reasonable request.

