## Supplemental Information for "Genetically-encoded discovery and development of peptide-macrocycle imaging agents for PD-L1"

### Materials and Methods

**Focused library production**

Library preparation procedure was similar to the protocol in our previous publication^1^. In short: PCR was used to convert a ssDNA library (GenScript) to produce double-stranded (dsDNA) as follows:

1. 10x Taq buffer 12 μL

2. 10 mM dNTPs 2 μL

3. Taq Polymerase 2 μL

4. GSX/IDT Oligo FW (10 μM) 3 μL

5. GSX/IDT Oligo RV (10 μM) 3 μL

6. ssDNA library Oligo 2 μL

7. MgCl 2 (25 mM) 8 μL

8. Nuclease free water 85 μL

Cycling was performed using the following thermocycler settings:

a) 95 °C 60 s

b) 95 °C for 20 s

c) 54 °C 20 s

d) 72 °C 15 s

e) repeat b)-d) for 35 cycles

f) 72 °C 30 s

The PCR reaction was purified using Macherey-Nagel Gel and PCR clean up kit using an adapted protocol. The phage vector was digested overnight with KpnI HF (NEB Cat# R3142S) and EagI HF (NEB Cat# R3505S). The insert PCR fragment was then digested with KpnI HF and EagI HF, purified using Macherey-Nagel PCR clean up kit and ligated into the cut vector using T4 Ligase. The ligation products were then transformed into electrocompetent E. coli 10G prlA4 cells, and the transformants were grown overnight on E. coli ER2738 prlA4 cells to allow for phage production. Phage cultures were then centrifuged to remove cells and debris, and then the phage was precipitated by PEG precipitation, and resuspended in PBS. Concentration of phage library was determined through qPCR.

### Multi-Round Selection by Phage Display

All reagents used in the preparation of buffers and media were purchased from Thermo Fisher Scientific or Sigma-Aldrich, unless otherwise stated. Panning procedures were performed in 3 rounds. For Round 1 protein was immobilized on plate. Specifically, solution of 1 μg of His-Avi-PDL1 (ACROBiosystems, PD1-H82E5) in 100 μL of PBS was added to each well of Ni:NTA plate (Thermo Fisher Scientific, 15342). After slight agitation for 1 hour at room temperature, the solution was discarded. Blocking buffer (CMHBS, 1 mM CaCl_2_, 150 mM NaCl, 50 mM HEPES, 1 mM MgCl_2_, 1% BSA, 0.1 mM D-biotin) was added to each well and incubated with slight agitation for 1 hour at room temperature. Naïve phage library was depleted on NiNTA and streptavidin beads. Specifically, 100 μL of phage-display library (10^12^ pfu/mL), in blocking buffer was incubated for one hour with suspension (30 μL) of streptavidin beads (Thermo Fisher Scientific, 11205/6) 1 hour at room temperature. The library was then incubated with NiNTA beads (Thermo Fisher Scientific, 10104D). Blocked library was added to blocked PD-L1 coated well and incubated at 4 °C overnight. The following day, the library was removed, and the wells were washed 3 times with 300 μL of protein wash buffer (CMHBS, 0.1% Tween 20). Elution was performed with 200 μL of elution buffer (0.2 M glycine-HCl, pH 2.2). Following a 9-minute incubation with agitation, the solution was promptly aspirated and transferred to the new tube containing 30 μL of neutralization buffer (1 M Tris-HCl, pH 9). 200 μL of eluted phage was used for amplification (see amplification procedure).

Following amplification, libraries were mixed to yield an input for Round 2, which was depleted as in Round 1. For Rounds 2 and 3, 1 μg His-Avi-PDL1 (in 100 μL PBS) was immobilized on 10 μL of streptavidin-coated magnetic beads. Specifically, beads are washed 3 times with PBS to remove the storage buffer, then resuspended in protein solution and rotated for 1 hour at room temperature. For Round 3, blank streptavidin-coated magnetic beads (control) were included. The input for Round 3 (~10^11^ pfu/mL) was prepared by mixing the R2 amplification and GS29 library, which were mixed in a 1:1 ratio according to qPCR analysis. Mixed library was depleted as in Round 1. The protein/bead solution, input library, and other reagents were added to a KingFisher 96 Deep-well plate (Thermo Fisher Scientific, 95040450) as follows:

Row A: 900 μL of 1x CMHBS, 100 μL protein/bead solution

Row B: reserved for KingFisher 12-tip comb (Thermo Fisher Scientific, 97003500)

Row C: 1 mL 1x CMHBS

Row D: Blocking buffer

Row E: phage library (10^11^ or 10^10^ pfu/mL in blocking buffer)

Row F-H: wash buffer

Row 2A-C: wash buffer (for Round 3 panning)

The following steps were performed using a KingfisherTM Duo Prime Purification System with a magnetic comb to transfer the beads. The program is as follows:

a) Collect comb from row B
b) Collect beads from row A on comb
c) Wash beads in row C – 30 seconds
d) Block beads in row D – 1 hour
e) Bind phage in row E – 1.5 hours
f) Wash beads in row F – 1 minute
g) Wash beads in row G – 1 minute
h) Wash beads in row H – 1 minute

For Round 3:

i) Wash beads in row 2A – 1 minute
j) Wash beads in row 2B – 1 minute
k) Wash beads in row 2C – 1 minute

At the end of the program, the contents of each well from the final row were transferred to individual microcentrifuge tubes placed on a magnetic rack to capture the beads. For Round 2, the supernatant was discarded, the beads were resuspended in elution buffer (200 μL) and rotated for 9 min. The supernatant was then removed from the beads and combined with 30 μL of neutralization buffer in a fresh tube. The neutralized R2 output was amplified as in Round 1. For Round 3, the supernatant was discarded, the beads were resuspended in 30 μL of nuclease-free water and the samples were boiled for 10 minutes at 95°C to release phage ssDNA. The supernatant containing the ssDNA was removed from the beads and employed in qPCR analysis and PCR-based preparation of DNA for NGS.

### Amplification of Phage Libraries

In a 15 mL culture tube, 1.5 mL of 2YT was inoculated with 15 μL of ER2738 (Biosearch Technologies, 60522-2) overnight culture and shaken for 1 hour, 37 °C, 200 rpm. 200 μL (protein panning) or 860 μL (cell panning) of eluted phage was added to the above mixture and amplified for 3 or 4 hours at 37 °C, 200 rpm, respectively. The amplified culture was centrifuged for 15 minutes at 4000xg, 4 °C. The supernatant was transferred to a microfuge tube with 1/5 volume of PEG/NaCl. Phage was incubated at 4 °C until completely cooled (Round 1 protein) or overnight (Round 2 protein, cell panning). It was then centrifuged at 10,000xg for 10 minutes, 4°C. The pellet was resuspended in 500 µL of PBS, and stored at 4 °C. If necessary, a second PEG purification/precipitation was performed by transferring the suspension to a new tube, with 1/5 volume of PEG/NaCl (~200 µL) and incubating for 30 minutes to 1 hour on ice before proceeding with centrifugation at 10,000xg and resuspension in PBS.

### Panning of focused phage and phagemid libraries on cells

Cell based panning procedures were similar to procedures described in our previous publications^2^. Input libraries were prepared in incubation buffer at 10^11^ pfu/mL. If necessary, inputs were depleted on negative cells for 1 hour on ice. Cells were harvested and washed 2 times with 3 mL of cell wash buffer (CMHBS, 0.1% BSA). Cells were then resuspended in incubation buffer (CMHBS, 1% BSA) at 10^6^ cells/100 μL. Each replicate of panning was performed in FACS tube: we first added 100 μL of cells (10^6^), centrifuged the tube for 5 minutes, 500xG, 4 °C. The tube was briefly inverted to drain the supernatant. The pellet was resuspended in 100 μL of depleted library. The cell/input mixtures were incubated for 1 hour on ice, followed by a desired number of washes (n=1-6). In each wash, we added 3 mL of cell wash buffer; the suspension of cells in wash buffer was incubated on ice for 5 minutes (“5-min soak”), followed by centrifugation for 5 minutes, 500xG, 4 °C. Following the final wash, cell pellets were resuspended in 1 mL of wash buffer and used either for amplification or DNA isolation and NGS preparation as described in our previous publication^2^. Specifically, the cell suspension was centrifuged in 1.5 mL Eppendorf tube. Supernatant was discarded and the pellet was resuspended in 50 μL of 1x HF (New England Biolabs, B0518S) with 0.1 mg/mL RNase A (Thermo Scientific, EN0531), then vortexed for 1 minute at maximum speed to lyse the cells. Following centrifugation, the supernatant was added to 1 μL of 1% Proteinase K (EO0491). Samples were heated at 56 °C for 10 minutes, followed by 95°C for 3 minutes. After centrifugation, the supernatant was used for PCR amplification and qPCR analysis (see below).

### Panning of Focused Library on PD-L1 protein

His-Avi-PDL1 protein (1 μg, 0.1 μg, or 0.01 μg) were incubated with suspension of streptavidin-coated magnetic beads (10 μL) as described for multi-round selection. Blank streptavidin-coated beads were included as a control. The focused library input was prepared at 10^11^ pfu/mL in blocking buffer. The KingFisher plate was prepared and loaded as described for Round 3 multi-round screening on protein, and the samples were processed by boil elution to yield ssDNA for NGS. The focused library screening was performed similarly for both polyvalent (phage) and monovalent (phagemid) libraries with only one difference: eluted ssDNA from phagemid panning was amplified using phagemid-specific primers whereas ssDNA from phage was processed using phage-specific primers.

### Preparation of phage DNA for Next-Generation Sequencing

Input and output samples from each round in multi-round selection or from focused library screening were prepared for NGS using a 2-step PCR scheme.

Step 1 reactions were amplified as follows:

1. 15 µL CloneAmp HiFi PCR Mix (Takara, 639298)
2. 1 µL of 96 reverse primer (IDT)
3. 1 µL NF10 forward primer (IDT)

Cycling was performed using the following thermocycler settings:

a) 98 °C, 3 minutes
b) 98 °C, 10s
c) 50 °C, 20s
d) 72 °C, 20s
e) Repeat b)-d) for 25 cycles
f) 12 °C, 1 minute

2-8 µL of Step 1 reaction were added to Step 2 reactions as a template. The reactions were amplified as follows:

1. 10 µL CloneAmp Hifi PCR Mix
2. 0-6 µL nuclease free water
3. 2-8 µL Step 1 template
4. 1 µL reverse primer (TEV_RV2; IDT)
5. 1 µL forward primer (SDBF; IDT)

Cycling was performed using the following thermocycler settings:

a) 98 °C, 10s
b) 50 °C, 20s
c) 72 °C, 20s
d) Repeat b)-d) for 15 cycles
e) 12 °C, 1 minute

### Maturation library production, screen and analysis

Top performing sequences in the focused library screening were submitted to our maturation tool, which analyses the positional scan data of sequences and estimates which mutations at which positions may improve binding. These oligos, synthesized by IDT, were cloned into the monovalent M13 phagemid vector pADL-100 (Antibody Design Labs, CAT# PD100). Briefly; PCR was used to produce double stranded DNA from these Affinity Maturation oligos. The oligo was amplified using the following thermocycler settings:

a) 95 °C 60 s

b) 95 °C for 20 s

c) 56 °C 20 s

d) 72 °C 30 s

e) repeat b)-d) for 35 cycles

f) 72 °C 30 s

The PCR reaction was purified using Macherey-Nagel Gel and PCR clean up kit using an adapted

protocol. The pADL-100 phagemid vector was digested overnight with BglI (NEB Cat# R0143S). The insert PCR fragment was then digested with BglI, purified using Macherey-Nagel PCR clean up kit and ligated into the cut vector using T4 Ligase. The ligation products were then transformed into electrocompetent *E. coli* TG1 prlA4 cells, and the transformants were grown overnight in ampicillin + (100 μg/ml final concentration) 2YT growth media with 1% w/v glucose at 30°C at 200 rpm. A subculture of the amplified transformed cells was made using 1% v/v and grown in 2YT until OD_600_=0.5. At that time the M13 helper phage M13KO7 (Antibody Design Labs CAT# PH010S) was added at MOI = 10, and incubated at 37 °C with shaking at 50 rpm for 1 hour. After 1 hour, ampicillin (100 μg/ml), kanamycin (50 μg/ml), and IPTG (200 µM) were added and incubated overnight at 30 °C with shaking at 200 rpm for phage production. The following day, cultures were centrifuged to remove cell debris and phage purification was done through PEG precipitation as described above.

Screening of matured libraries on protein and cells was performed with the same protocol as the focused library outlined above.

### Focused Library Analysis

NGS reads from input and output focused-library samples were mapped to the designed peptide sequence list. Read counts for each peptide were normalized to counts per million, and enrichment was calculated as the fold change of output abundance relative to input abundance. Enrichment values were visualized as Manhattan plots, with each point representing a unique peptide and colors indicating sequence family. For positional-scan libraries, fold-change values were also compared with the corresponding parent sequence to identify substitutions that were deleterious or beneficial. Enriched variants and favorable substitution patterns were used to guide subsequent library design and peptide nomination for synthesis and validation.

### Cell culture

Chinese Hamster Ovary cells stably overexpressing PD-L1 (CHO-PDL1) were obtained from Acro Biosciences (CAT# SCCHO-ATP0TP077H). Parental CHO-K1 cells were obtained from BPS Bioscience (CAT# 60545). Both CHO lines were cultured in Ham's F-12K (Kaighn's) Medium (Thermo Fisher, Cat# 21127022) supplemented with 10% fetal bovine serum (FBS; Corning, Cat# 35-015-CV) and 1% penicillin-streptomycin (PenStrep; Thermo Fisher, Cat# 15140122). To maintain selection pressure for the CHO-PDL1 line, 2 µg/mL puromycin (Thermo Fisher, Cat# A1113802) was added to the culture medium. MDA-MB-231 human breast adenocarcinoma cells were provided by Dr. Frank Wuest (University of Alberta) and were maintained in Dulbecco's Modified Eagle Medium containing GlutaMAX (DMEM; Thermo Fisher, Cat# 10569010) supplemented with 10% FBS and 1% PenStrep.

All cell lines were maintained in a humidified incubator at 37 °C under a 5% CO₂ atmosphere and were strictly restricted to experimental use between passages 5 and 20 (P5–P20). Cells were cultured in either T75 (Thermo Fisher Cat# 156499) or T175 (Thermo Fisher Cat# 159910) tissue culture flasks, varying the vessel size based on the experimental yield required. To ensure stability across passages, PD-L1 expression was regularly verified via flow cytometry using an anti-PD-L1 antibody conjugated to Alexa488 fluorophore (R&D systems, Clone# Hu124/CAT# FAB10348G-100UG).

Upon reaching approximately 80% confluence, the culture medium was removed, and the cell monolayer was detached using TrypLE Express (Thermo Fisher, Cat# 12605028). Following complete detachment, the enzymatic activity was quenched by adding complete growth medium. Cells were collected via centrifugation, and total cell counts and viability were evaluated using a hemocytometer via trypan blue exclusion (Thermo Fisher, Cat# 15250061). Harvested cells were subsequently resuspended in staining buffer at a final density of 1×10⁶ cells/100 µL for downstream bio-panning and flow cytometry assays.

### Synthesis and Characterization of compounds

### Peptide Synthesis Method 1

Peptides 8911-8913, 10246-10247, 10302, 8455-8458, 8581-8587, 8589, 8604-8607, 8609, 8741-8748, 8778, 8812-8813, 9635, 9815-9816, 8414-8418, 9811, 9847-9849, 9881-9888, 8521, and 8523 were synthesized on a PreludeX peptide synthesizer (Gyros Protein Technologies) by standard Fmoc solid chemistry using Rink Amide AM resin. Fmoc-protected amino acids, HBTU, and Rink Amide AM resin (200-400 mesh, 0.92 mmol/g, ChemPep, Wellington FL USA). Each coupling to 200 mg of Rink Amide AM resin was performed for 1 h with 5:5:10 eq of AA:HBTU:DIPEA. Peptides were cleaved from the resin by using a TFA/EDT/TIPS/Water (89.9/2.28/4.54/2.28 v/v) or TFA/TIPS/Water (95/2.5/2.5 v/v) for peptides with C-terminal Azido-Lysine. Cleaved peptides were precipitated and washed with ice-cold diethyl ether and purified by Combiflash (C18 column, 214 and 254 nm) and lyophilized. Disulfides were made using Iodine oxidation. 200 mM I_2_ in MeOH was added dropwise to a solution of 10-20 mg of crude peptide (10-20% Acetonitrile or DMSO in water, 0.1% TFA). The reaction was then quenched with 1 drop of 1 M ascorbic acid 30 seconds after the presence of color in solution. Peptides were then purified on Waters Preparative HPLC (214 nm and 254 nm), the fractions analyzed with LCMS (214 nm), and Lyophilized.

### Peptide Synthesis Method 2

Peptides 8404-8406, 9970-9974, 9516, 9641-9647, 8885-8889, 8915-8916, 8774, 8776, 10177-10179, 9186-9189, 10815-10816, 8250, 9220-9222, 9317-9319, 9351.1, 9016-9019, 9857.1, 9858.1, 9871-9874, 10361-10364, 9859, 9875-9878 were purchased from GeneScript, MedChemExpress, Peptide Lab, or Scitides using their proprietory synthesis protocols. All peptides were provided by the suppliers with a standard quality control (HPLC trace, mass spectrometry verification).

### Peptide Synthesis Method 3

Peptides 9844, 10005, 10007, 10200, 10210, 10300, 10310, 9015, 8191-8193, 8195-8198, 9372, 9481, 9485, 9489, 9501-9502, 9505, 9509, 9514-9515, 9517-9519, 9542, 9552, 9732, 9929, 8194, 9340, 9572, 9639, 9669, 9734-9736, 8851-8854, 8941, 8943-8944, 8775, 9482-9483, 9957-9959, 8129, 9632, 9682, 9745, 9834-9835, 9846, 9880, 9890, 9895-9899, 9975, 9870, 10002, 9475-9478, 9300, 9320-9321, 9331, 9325-9331, and 9351-9352 were synthesized by WuXi AppTec using the following representative protocols:

Peptides with similar modifications to 9682, 9880, 9890, 9325 were synthesized on Sieber resin (0.10-0.50 mmol, 0.26 mmol/g), whereas peptide 9870 (carbon bridge macrocycle) was synthesized on MBHA resin (0.20 mmol, 0.26 mmol/g) and employed custom-made amino acid Int_9870 (synthesis described below). Peptides with similar modifications to 9734 and 9735 were synthesized on Rink amide MBHA resin (0.35 mmol, 0.31 mmol/g). Resin swelling was performed in DMF under nitrogen at 20–25 °C for 20–50 min, followed by Fmoc deprotection using 20% piperidine in DMF for 10–15 min with intermediate DMF washes. Coupling was carried out using HBTU or HATU with DIEA in DMF at 20–25 °C for 0.5–1 h. Peptides 9682, 9734, and 9735 used Fmoc-Lys(Dde)-OH as the coupling amino acid, whereas peptides 9870, 9880, 9890, 9325, and 9328 employed Fmoc-Gly-OH or Fmoc-Lys(biotin)-OH depending on the target sequence. In selected cases (9880 and 9890; Amide bridged peptides), Fmoc-Dap(Alloc)-OH and Fmoc-Asp(OAll)-OH were used instead of cysteines. After de-Alloc/de-OAllyl (Pd (PPh3)4 (0.20 eq), PhSiH3 (40.0 eq)), HOAt/DIC was used to cyclize amide bridge peptides with overnight agitation. Int_9870 (Scheme S.1. - S.4.) was then used in place of cysteines during peptide synthesis of 9870 following the coupling and de-Alloc/Deallyl procedures described above. Dde deprotection for peptides 9734 and 9735 was performed using 3% hydrazine hydrate in DMF for 15 min at 25 °C.

For peptide cleavage, all resins were washed with DMF and MeOH and dried under vacuum. Peptide 9682 was cleaved using 90% TFA, 2.5% TIS, 2.5% H2O, 2.5% 3-mercaptopropionic acid, and 2.5% thioanisole at 20 °C for 1.5 h. Peptide 9870 was cleaved using 90% TFA, 5% TIS, 2.5% H2O, and 2.5% 3-mercaptopropionic acid at 20 °C for 1.5 h. Peptides 9880 and 9890 were cleaved using 92.5% TFA, 2.5% TIS, 2.5% H2O, and 2.5% 3-mercaptopropionic acid at 20 °C for 1.5–2 h. Peptides 9325 and 9328 were cleaved using 92.5% TFA, 2.5% DTT, 2.5% H2O, and 2.5% TIS at 25 °C for 2 h. Peptides 9734 and 9735 were cleaved using 92.5% TFA, 2.5% TIS, 2.5% H2O, and 2.5% 3-mercaptopropionic acid at 20 °C for 2 h. In each case, the crude peptide was precipitated with cold tert-butyl methyl ether or isopropyl ether, collected by centrifugation, washed, and dried under vacuum.

Oxidative cyclization was performed for peptides 9682, 9325, 9328, 9734, and 9735 by dissolving the crude peptide in H2O/MeCN and adding I2/MeOH dropwise until a persistent yellow color was observed. Peptide 9682 was cyclized in H2O/MeCN (200/300 mL) using 0.5 M I2/MeOH at 20 °C for 30 min. Peptides 9325 and 9328 were cyclized in H2O/MeCN (50/50 mL) using 0.2 M I2/MeOH at 20 °C for 10 min, followed by quenching with aqueous sodium thiosulfate. Peptide 9734 was cyclized using 0.5 M I2/MeOH at 20 °C for 30 min, and peptide 9735 was cyclized under the same conditions at 25 °C for 30 min, followed by sodium thiosulfate quenching.

All crude peptides were purified by preparative HPLC under TFA conditions (mobile phase A: 0.075% TFA in H2O; mobile phase B: acetonitrile). Peptide 9682 was isolated as a purple solid, peptides 9870, 9325, 9734, and 9735 were isolated as white or off-white solids, and peptides 9880 and 9890 were isolated as white solids. Final yields ranged from 1.15% to 3.62% for the compounds described here, with purities of 93.8–99.4% after purification. Identity and purity were confirmed by LCMS and analytical HPLC using compound-specific methods.

### Synthesis of the Int_9870 for synthesis of carbon-bridged macrocycle

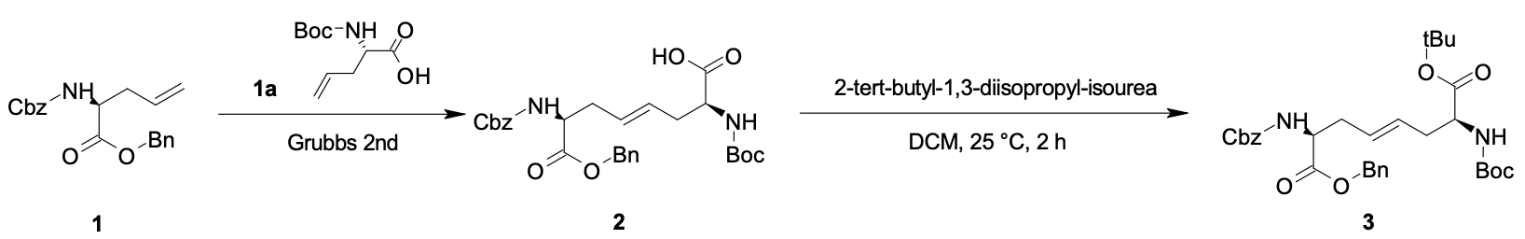

Scheme 1: Step 1 and 2 of carbon bridge building block synthesis

To a solution of compound 1 (5.50 g, 15.3 mmol, 1.00 eq) and compound 1a (3.95 g, 18.4 mmol, 1.20 eq) in DCM (165 mL) was added benzylidene-[1,3-bis(2,4,6-trimethylphenyl)imidazolidin-2-ylidene]dichloro-ruthenium;tricyclohexylphosphane (650 mg, 765 μmol, 0.05 eq) under N2 atmosphere, and the reaction mixture was stirred at 40 °C for 4 h (Scheme S.1.). The reaction mixture was concentrated, and the residue was purified by prep-HPLC (TFA condition) to afford compound 2 (2.00 g, 3.45 mmol, 22.5% yield, 90.9% purity) as a brown oil, which was confirmed by LCMS and 1H NMR. Compound 2 (2.00 g, 3.45 mmol, 1.00 eq) was then treated with 2-tert-butyl-1,3-diisopropylisourea (4.15 g, 20.7 mmol, 6.00 eq) in DCM (10 mL) at 25 °C for 2 h (Scheme 1). After filtration and concentration, the residue was purified by column chromatography to give compound 3 (1.65 g, 2.83 mmol, 82.0% yield) as a yellow oil, confirmed by 1H NMR and LCMS.

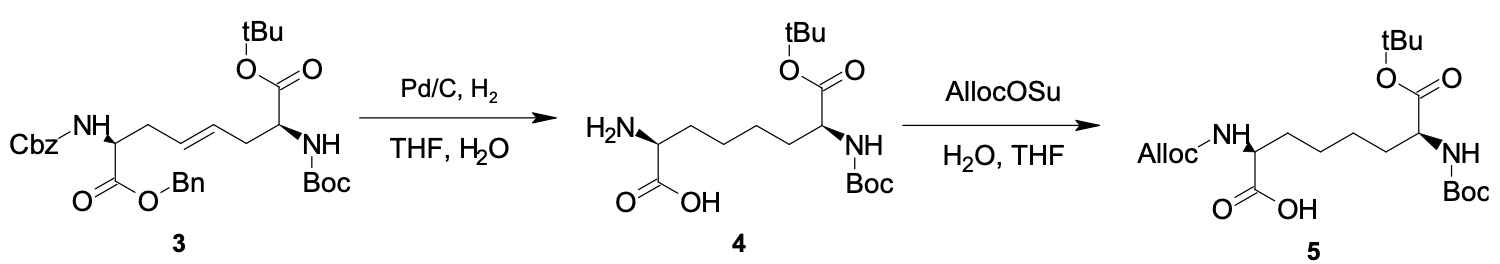

### Scheme 2: Step 3 and 4 of carbon bridge building block synthesis

Compound 3 (1.65 g, 2.83 mmol, 1.00 eq) was hydrogenated in THF (17 mL) in the presence of Pd/C (165 mg, 10%) under H2 (50 psi) at 40 °C for 12 h. The reaction mixture was filtered to afford compound 4 (1.02 g, crude) as a colorless liquid, which was carried directly into the next step without further purification (Scheme S.2.). Compound 4 (1.02 g, crude) was then treated with NaHCO3 (476 mg, 5.66 mmol, 2.00 eq) and allyl (2,5-dioxopyrrolidin-1-yl) carbonate (564 mg, 2.83 mmol, 1.00 eq) in THF/H2O (17 mL/17 mL) at 25 °C for 1 h. After acidification and extraction, the crude product was obtained as compound 5 (1.25 g, crude) as a yellow oil (Scheme S.2.). Compound 5 (1.25 g, crude) was coupled with compound 6 (510 mg, 4.21 mmol, 1.50 eq) in the presence of K_2_CO_3_ (777 mg, 5.62 mmol, 2.00 eq) in ACN (13 mL) at 40 °C for 12 h. The reaction mixture was purified by column chromatography to furnish compound 7 (1.16 g, 2.39 mmol, 85.1% yield) as a colorless oil (Scheme 3).

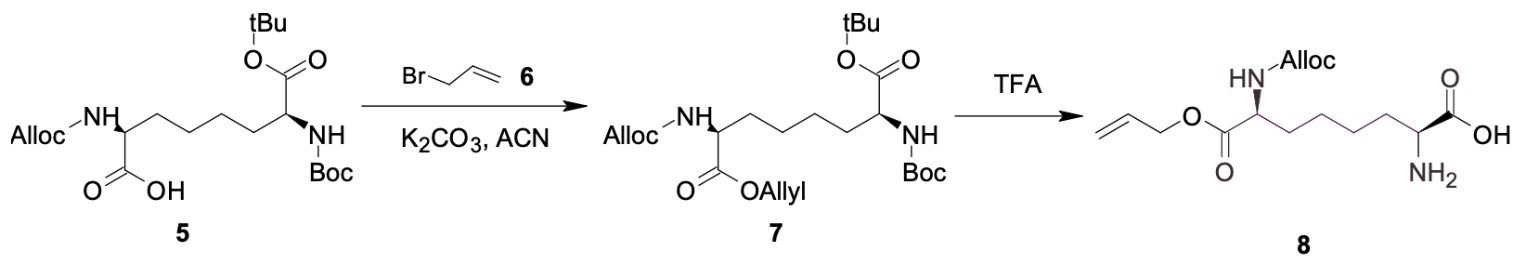

### Scheme 3: Step 5 and 6 of carbon bridge building block synthesis

Compound 7 (1.16 g, 2.39 mmol, 1.00 eq) was deprotected with TFA in DCM (18.5 g, 12.0 mL, 67.7 eq) at 25 °C for 12 h to afford compound 8 (1.05 g, crude, TFA salt) as a yellow oil (Scheme S.3.). Finally, compound 8 was reacted with NaHCO3 (1.39 g, 16.6 mmol, 7.00 eq) and FMOC-OSU (760 mg, 2.25 mmol, 0.95 eq) in THF/H2O (11 mL/11 mL) at 25 °C for 1 h (Scheme S.4.). After acidification, extraction, prep-HPLC purification, and a second purification by column chromatography, Int_9870 was obtained as a white solid (436 mg, 751 μmol, 31.6% yield, 94.9% purity), and its structure was confirmed by LCMS, HPLC, SFC, and 1H NMR.

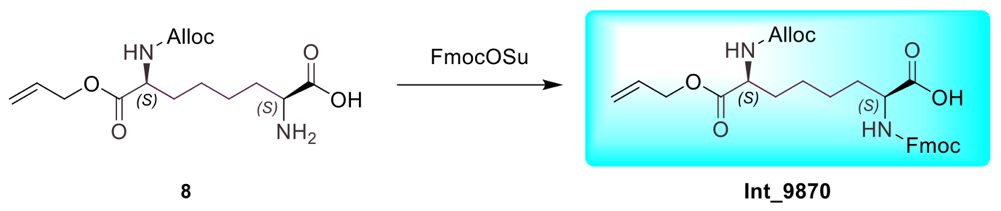

### Scheme 4: Fmoc protection of compound 8 into Int_9870

### Late Stage functionalization of peptides with meta-dichloroxylene (MCX) linker

This is a representative procedure for the synthesis of MCX cyclized peptides 8414-8418 and 8921-8923 (supplementary Fig 15b). In a vial, a 494 µL solution (5-10% ACN) of 2 mM peptide, 0.6 mM TCEP, 3.4 mM MCX linker, and 100 mM sodium bicarbonate buffer (pH 8.5) was made. The reaction was left for 2 h at room temperature before purifying on CombiFlash (C18, 12 g column, 214 nm). Isolated fractions were analyzed with LCMS to verify purity and product identity.

### Synthesis of reference compounds

Derivatives of WL-12 and BMS-986229 compounds (Scheme 5) have been synthesized by WuXi AppTec and the identity and purity of the compounds have been confirmed by LCMS and HPLC. The K_D_ of the synthesized compounds have been confirmed by SPR to be as follows: K_D_=0.9 nM for 8450; K_D_=1-2 nM for WL-12 as depicted in Fig. 1d, and employed in Fig. 7b for labeling with ^68^Ga; K_D_=0.003 nM for 9929 (Supplementary Fig. 1c). K_D_=0.06 nM for 8329 with non-radioactive Ga.

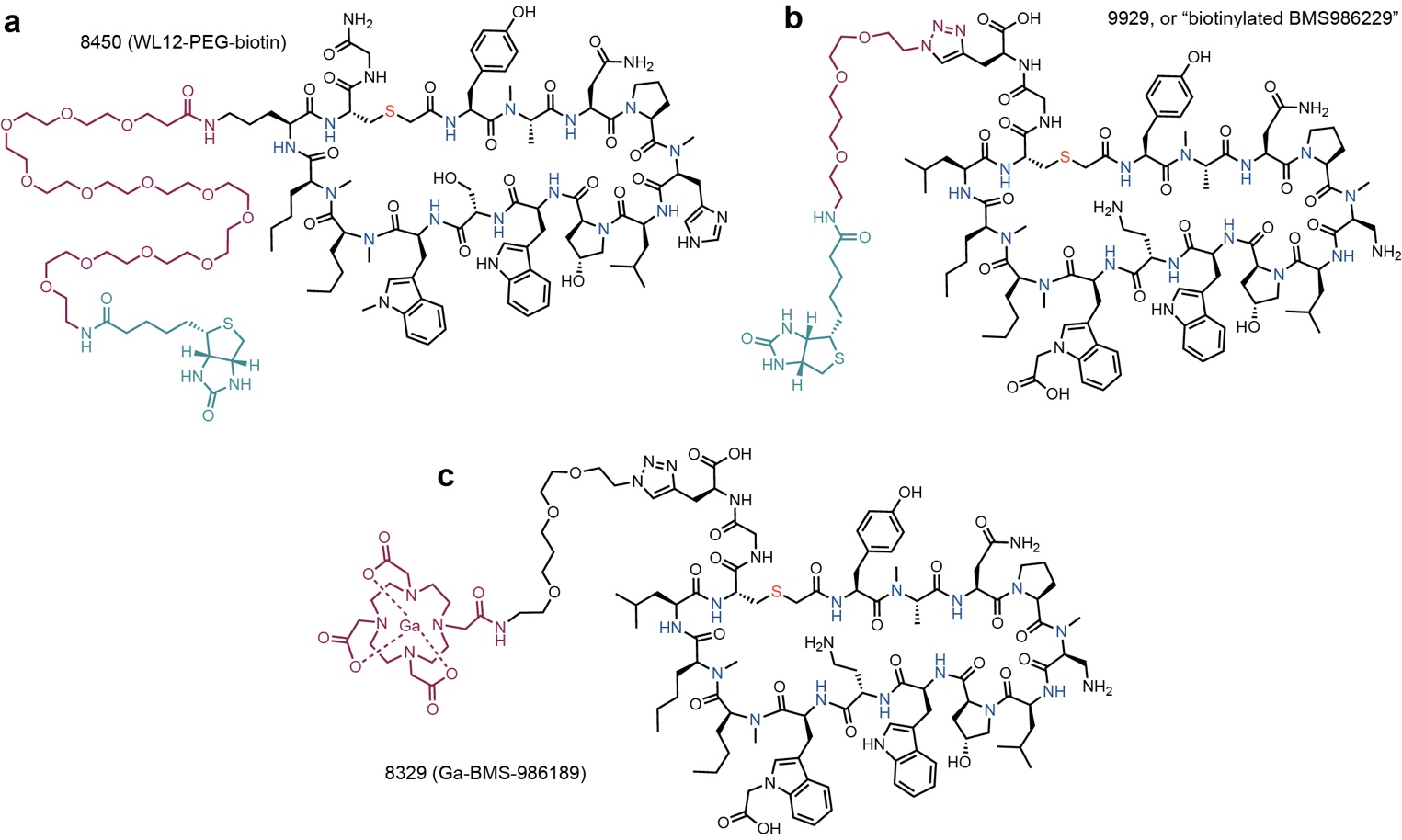

### Scheme 5: Structures of BMS-macrocycles used as references in this publication

**a,** Biotinylated derivative of WL-12 (Fig. 1d), compounds 8450 in this report, was employed as positive control in FACS measurements in Supplementary Fig. 1f. **b,** Biotinylated derivative of BMS-986229, compound 9929 in this report, was employed as positive control in SPR and FACS measurements in Supplementary Fig. 1c. **c,** Derivative of BMS-986229 that contains DOTA-chelator in place of fluorine tag in BMS-986229 (Fig 1c); the radioactive version of this compound with ^68^Ga has been used for animal PET imaging (Fig 7d).

### Stability of Peptide in previously Frozen Mouse Serum

The following procedure has been performed by WuXi AppTec. Frozen CD-1 mouse serum (WuXi, Cat#: WuXi_MsS_20250211, male). Analytical grade inorganic salts were purchased from Sinopharm Chemical Reagent Co. (Shanghai, China). HPLC grade organic solvents were purchased from Sigma-Aldrich, USA. Distilled water, prepared from demineralised water, was used throughout the study. Atenolol (Cat#: A7655-1G) and Tamoxifen (Cat#: T5648) were purchased from Sigma (St. Louis, MO, USA), and Ketoconazole (Cat#: K0045) was purchased from TCI (Tokyo Chemical Industry, China). Sodium Phosphate dibasic anhydrous was purchased from SINOPHARM (20040618-500g), Sodium dihydrogen Phosphate dihydrate was purchased from SINOPHARM (20040718-500g), and dimethyl sulfoxide was purchased from AMRESCO (0231-500mL). Distilled water was prepared with Millipore Direct-Q water purification system. Analytical grade inorganic salts were purchased from Sinopharm Chemical Reagent Co. (Shanghai, China). Human plasma (male:female = 1:1, v/v) was purchased from Bioreclamation (Cat# BRH470404). CD-1 mouse plasma (male:female = 1:1, v/v) was obtained from HDB (Cat# HDB_MsP_20240410). The HTD 96b Complete Unit (Cat# 1006), HTD 96a/b Dialysis Membrane Strips (Cat# 1101), and HTD 96a/b Adhesive Sealing Film (Cat# 1102) were purchased from HTDialysis.

### Stability of Peptide in previously Frozen Human Serum

The following procedure has been performed by WuXi AppTec. Stock solution of test compounds were prepared in 100% DMSO. The stock solution for each compound was diluted into 400 μM with ACN/H2O, diluted into frozen human serum to achieve a final concentration of 2.0 μM in duplicate, and incubated at 37 °C. Aliquots of 50 μL sample were collected at 0, 0.25, 0.5, 1 and 2 h and mixed with 300 μL of ice-cold 100% acetonitrile containing internal standard. After centrifuging the plate at 4000 rpm for 20 min, 50 μL of supernatants were added to 100 μL H2O, mixed well and analyzed with LC-MS/MS.

### Protein Binding study

The following procedure has been performed by WuXi AppTec. 4 membrane strips were soaked in water for 60 min. *Note that each membrane strip consists of two membrane sheets.* Membranes were then taken out of the water, separated, and soaked in 20% ethanol overnight. Membranes were rinsed with water three times to remove traces of alcohol before soaking in RPMI 1640 Medium without protein until ready to assemble. Extra membranes were stored in 20% EtOH at 4ºC until ready to use. Compound stocks were prepared at 200 μM: 10 μL of 10000 μM diluted into 200 μM with mixture of 50% acetonitrile. 4 μL of 200 μM compound stock solutions were added to 796 μL plasma in glass vials and mixed thoroughly. The final compound concentration in spiked plasma is 1.0 μM for each compound. In an assembled HTDialysis 96-well plate, each well consisted of an upper section (sample) and a lower section (dialysate) separated by the membranes. 150 μL of PBS was added into the lower section (n=3). Then 150 μL of spiked plasma was added into the upper (n=3). The plasma HTD plates were sealed with sealing films and incubated in 37 ºC for 5 hours in 5% CO2 incubator. Aliquots of 50 μL spiked plasma into a 96 wells plates in 3 replicates were used as T0 h. After incubation, 50 μL of PBS sample and 50 μL of plasma samples from the lower and upper compartment, respectively, and matrix matched with blank plasma or blank PBS. Matrix-matched samples (100 μL) were quenched with 400 μL internal standard in 100% ACN and shaken on a Thermomixer (750 rpm for 5 min). after shaking, the plate was centrifuged (4000 rpm, 30 min, 4º C). 100 μL from all samples were then transferred into a daughter plate containing 400 μL ddH2O, mixed, and analyzed using LC-MS/MS.

### Kinetic Solubility Study

The following procedure has been performed by WuXi AppTec. Buffer was prepared at pH 7.4 by mixing 40.5 mL of solution A (0.2 M Na_2_HPO_4_) and 9.5 mL of solution B (0.2 M NaH_2_PO_4_.2H_2_O) into 50 mL distilled H2O, adjusted pH to 7.4. Stock solution of each test compound was prepared by dissolving the pure reference compounds individually in DMSO to yield a final concentration of 10 mM. An aliquot of 10 μL stock solution was spiked into 490 μL of phosphate buffer (pH 7.4) to get 200 μM test solution in triplicate. The final concentration of DMSO was 2% (v/v). The test solutions were vortexed for 5 min, shaken for 1 h, and then stood still at room temperature for another 2 h. Then, the test solutions were centrifuged at 14000 rpm for 10 min. An aliquot of 10 μL stock solution was spiked into 490 μL of 50% acetonitrile to get 200 μM working solution. Then, the working solution was diluted with 50% acetonitrile to get calibration standards at concentrations of 100, 50, 30, 10, 3, 1, 0.3 and 0.1 µM. Aliquots of 10 μL above calibration standards at different concentrations were added into 90 μL phosphate buffer generate series calibration standards (10.0 μM, 5.0 μM, 3.0 μM, 1.0 μM, 0.3 μM, 0.1 μM, 0.03 μM, 0.01 μM). All samples were then analyzed with LC-MS/MS.

### Measurement of LogD of peptide macrocycles

The following procedure has been performed by WuXi AppTec. Test compounds were weighed and dissolved in 100% DMSO to get 10 mM stock solution. 10 μL of 10 mM stock solutions were dispensed into a 96-well plate and then 300 μL of octanol was added to the same well. The sealed plate was then agitated on a plate shaker for 5 minutes. After centrifuging the plate at 2000 rpm for 5 minutes, 600 μL of potassium phosphate buffer (pH 7.4) was added to achieve a final concentration of 110 μM. The two phases were then mixed on the plate shaker vigorously for 1 h at 25°C and centrifuged (2000 rpm, 5 minutes). 10 μL of water phase samples from the water phase plate were mixed with 10 μL of 50% ethanol (1:2 dilution). 180 μL of 50% ethanol was added into the water phase sample plate (1:10 dilution) to achieve 1:20 dilution. 30 μL from the octanol phase (the upper phase) were then transferred into a new 96-well plate with 90 μL 50% Ethanol (1:4 dilution). To prepare samples for analysis, 50 μL of 1:20 water phase sample was mixed with 100 μL of 50% Ethanol (containing IS) into the water phase sample plate to achieve 1:60 dilution. In a new 96-well plate, 120 μL of 1:4 diluted octanol samples were mixed with 60 μL 50% Ethanol with IS to achieve 1:6 dilution. Samples were mixed well and analyzed with LC-MS/MS.

### Determination of K_D_ by SPR

Surface Plasmon Resonance (SPR) measurements were conducted using a Biacore 1K Instrument (Cytiva) at 25 °C in 1x PBS-P+ (20 mM phosphate buffer, 2.7 mM KCl, 137 mM NaCl, 0.05% Surfactant P20, pH 7.4), prepared by diluting 10x PBS-P+ stock (Cytiva, CAT# 28995084) and supplementing with 1% DMSO and 0.05% BSA. The recombinant human PD-L1-His ligand protein (ACROBiosystems, CAT# PD1-H5229) was immobilized onto two types of Series S Sensor Chips, NTA (Cytiva, CAT# BR100532) or CM5 (Cytiva, CAT# BR100530). For HIS capture on the NTA chip, standard immobilization conditions were applied using the NTA Reagent Kit (Cytiva, CAT# 28995043). Covalent immobilization on the CM5 chip was carried out at pH 5.5 through standard amine coupling using 1-ethyl-3-(3-dimethylaminopropyl)carbodiimide (EDC) and N-hydroxysuccinimide (NHS) followed by Ethanolamine blocking.

Following immobilization, peptide affinity measurements were made with the serially diluted solution (ranging from 0.5 nM – 10000 nM, based on K_D_) in assay buffer via single-cycle or multi-cycle kinetics. The association and dissociation phases were monitored at flow rate of 30 µL/min. The HIS tag captured surface was regenerated in assay buffer with 350 mM EDTA at flow rate 30 µL/min for 60 s followed by 0.5 mM NiCl_2_ and covalently immobilized surface was regenerated in assay buffer with 1 M MgCl_2_. Obtained data through double referenced subtraction was analyzed with the Biacore™ Insight Evaluation Software (Cytiva) and K_D_ values were estimated from 1:1 binding kinetics fit model.

### Determination of EC_50_ by ELISA

Enzyme-Linked Immunosorbent Assay (ELISA) measurements were performed on Pierce Nickel Coated Plate (Clear, 96-Well, Thermo Fisher, CAT# 15442) at room temperature. The plates were coated with recombinant human PD-L1-His (Acro Biosystem, CAT# PD1-H5229) in 100 µL (0.5 µg/well) PBS (pH 7.4) for 1 hour at room temperature. Following immobilization, coated plates were washed three times with 300 µL of PBS and subsequently blocked with 200 µL SuperBlock (Thermo Fisher, CAT# 37515) for 10 min. Plates were then washed three times with PBS with 0.05% Tween-20 (PBST). Biotinylated-peptide solutions (100 µL/well) were serially diluted across a desired concentration range in assay buffer (PBS, 0.05% Tween-20 & 0.5% BSA, pH 7.4), were added and incubated for 1 hour. For subsequent steps, ELISA plates were washed with PBST (300 µL, three times) in between the steps. For detection, Streptavidin-HRP (1:2000, STN-NH913, Acro Biosystem) was added to the plates and incubated for 45 minutes, followed by TMB substrate. The reaction was quenched with 1 M phosphoric acid stop solution. Absorbance was measured at 450 nm using a Multiskan SkyHigh Microplate Spectrophotometer (Thermo Fisher). Data analysis and EC50 values were calculated using GraphPad Prism (v.9.2; GraphPad Software, San Diego, CA).

### Determination of IC_50_ values by Biolayer Interferometry

Bilayer Interferometry (BLI) competitive inhibition experiments were performed at 30°C using a Gator Prime instrument (Gator Bio, Inc.) and 96-well black microplate. The anti-his biosensors (SKU 160009, Gator Bio, Inc) were pre-equilibrated in the assay buffer (PBS with 0.05% Tween-20, pH 7.4) for 120 s. The natural ligand of PD-L1, PD1-Avi/His (PD1-H82E4, ACRO Biosystems) was immobilized on the biosensor at a concentration of 100 nM. Following a wash step in assay buffer to establish a baseline, the ligand-loaded biosensors were dipped into competitive binding sample solutions. The solutions contained a fixed concentration (20 nM) of human PD-L1-Fc (ACRO Biosystems, PD1-H5258) pre-mixed with serially diluted peptide solutions at varying concentrations, depending on the potency of the peptides. The association and dissociation phases were monitored for 400 s and 600 s, respectively. Raw data was buffer subtracted using the parallel reference biosensors dipped in buffer alone. Data processing was executed using the Gator Analysis Software and GraphPad Prism (v.9.2; GraphPad Software, San Diego, CA) for IC_50_ estimation.

### Measurement of abundance with CaR

Peptide library screening by CaR-nMS was performed in positive-ion mode using a Q Exactive UHMR Orbitrap mass spectrometer (Thermo Fisher Scientific) equipped with a modified nanoflow electrospray ionization (nanoESI) source. The capillary temperature and S-lens RF level were set to 160 °C and 100, respectively. An automatic gain control target of 5×10^5^ and a maximum injection time of 200 ms were used. The resolving power was set to 25,000 for both full-MS and higher-energy collisional dissociation (HCD) spectra. HCD spectra were acquired using collision energies of 10–80 V. Data acquisition and preprocessing were performed using Xcalibur version 4.1. NanoESI emitters with an outer diameter of approximately 5 μm were pulled from borosilicate glass capillaries (1.0 mm outer diameter, 0.78 mm inner diameter) using a P-1000 micropipette puller (Sutter Instruments). A platinum wire was inserted into each tip and placed in contact with the sample solution. A voltage of approximately 0.8–1.0 kV was applied to the wire to generate the electrospray.

Screening solutions were prepared in aqueous ammonium acetate (200 mM, pH 7.2) and contained PDL1 (0.5 μM) and a library of 37 peptides, each at a concentration of 8 nM. PDL1–peptide complex ions with m/z values of 2,500–4,200 were isolated and subjected to HCD using nitrogen as the collision gas. Peptide release was first observed at a collision energy of 20 V and reached a maximum at 50 V. Accordingly, a collision energy of 50 V was used for peptide-library screening.

### Determination of IC_50_ values by VICA

Protein-peptide mixtures were pre-incubated for 30 minutes at 23 °C. Each solution (3.5 µL; n = 3) was applied to a fresh 3.9-mm VICA biosensor spot and incubated for a further 30 minutes. Solutions were removed by pipetting, slides were washed in PBS (pH 7.2) for 3 minutes, rinsed with distilled water, and dried with compressed air. Slides were imaged with the VICA lightbox and analyzed using proprietary VICA software (www.pavonis.app). dC quantified binding-induced changes at the sensor surface. Controls included buffer blank, PD-1 alone, PD-L1-His alone, and PD-1 + PD-L1 without peptide. PD-1 and PD-L1 were each used at 1.54 µM, with blank and single-protein controls prepared in parallel.

Replicate dC values were summarized as mean ± SD at each concentration. Dose-response curves used a logarithmic concentration axis and a four-parameter logistic model, in which the no-peptide response defined the upper signal, the high-dose plateau defined the lower signal, IC_50_ defined the midpoint, and the Hill term defined slope. Fit quality was assessed using R² values and residual inspection in Origin.

The shared controls provided a large signal window and separated surface background from PD-L1 capture and PD-1/PD-L1 complex formation. Measurements were technical replicates collected across multiple slides; consequently, the results quantify assay-level reproducibility but do not replace independent biological replication.

Although the two peptides exhibited an approximately four-fold difference in potency by BLI (IC_50_s of 39 & 175 nM respectively), produced nearly overlapping VICA inhibition curves with higher apparent IC_50_ values (Supplementary Fig. 10). Such assay-dependent differences are not unexpected and illustrate how peptide panels characterized across established orthogonal assays can be used to benchmark the dynamic range and performance of emerging analytical platforms.

### Determination of cell binding EC_50_ by flow-cytometry

Lyophilized biotinylated peptide stocks were reconstituted in DMSO to a concentration of 10 mM and serially diluted in staining buffer (SB) (CM-HBS 5% FBS) to generate a range of working concentrations (e.g., 0–20,000 nM) tailored to bracket the predicted EC_50_. Cultured cells were harvested, washed twice with 10 mL of wash buffer (WB) (CM-HBS 1% FBS) via centrifugation at 400×*g* for 7 min, and resuspended in SB. Cells were then aliquoted into 1.7 mL tubes at a density of 1x10^6^ cells per sample. After pelleting the cells and removing the supernatant, cell pellets were resuspended in 200 µL of the respective peptide dilutions and incubated for 1 hour at 4°C with periodic agitation every 15–20 minutes.

Following the primary incubation, cells were washed twice with 500 µL of SB. To detect bound peptide, cells were resuspended and incubated with Streptavidin, Alexa Fluor 488 (1.5 µg/sample) in 100–200 µL of SB for 1 hour at 4°C. During this secondary incubation and all subsequent steps, samples were protected from light using aluminum foil and agitated every 15–20 minutes. Cells were then washed twice with 500 µL of WB, and with a final wash of 200 µL of SB to ensure complete removal of the supernatant with a P200 pipette.

Final cell pellets were resuspended in 300 µL of SB, and fluorescence intensity was acquired on a BD Accuri C6 Plus Flow Cytometer at the medium flow rate for 100,000 events. Data files were exported to FlowJo software, where debris and dead cells were excluded to gate the single-cell population. To estimate EC_50_ values, a fluorescence intensity threshold was established using the 0 nM negative control (cells incubated with Streptavidin-Alexa Fluor 488 alone), defining a baseline where <3% of control events fell above the threshold. The percentage of the cell population shifting above this background threshold was calculated for each concentration, and the resulting dose-response curves were analyzed using GraphPad Prism 9.3.1 (GraphPad Software, San Diego, CA) for EC_50_ determination.

### Radiolabeling with ^68^Ga

DOTA conjugated peptides WL12, 9735, 9682, and BMS-986189 were radiolabeled with gallium-68 (^68^Ga). Gallium-68 was obtained from a 50 mCi ^68^Ge/^68^Ga generator using an automated GRP synthesis module (Scintomics GmbH, Fuerstenfeldbruck, Germany). The generator eluate was trapped on a CHROMAFIX PS-H+ (M) cartridge, and the retained ^68^Ga was eluted with 770 µL of 5 M sodium chloride solution.

A 750 µL aliquot of the ^68^Ga solution was mixed with 250 µL of 2.7 M HEPES buffer (pH 10.5), resulting in a final reaction pH of 4.0-4.5. Subsequently, 500 µL of this mixture was added to 25 µg of DOTA-peptides and heated at 90 °C for 10 min in a LoBind Eppendorf tube.

Radiolabeling efficiency was monitored by radio-TLC using silica gel 60 F254 plates and 0.1 M citric acid as the mobile phase. After labeling, the reaction mixture was diluted with 9 mL of deionized water and purified by solid-phase extraction on a Sep-Pak tC18 Light cartridge (Waters Corporation, Milford, MA, USA), which had been preconditioned with acetonitrile and water. The radiopeptides were eluted with 1.1 mL of a 1:1 ethanol/water mixture. The solvent was removed under reduced pressure, and the products were reformulated in 200 µL saline. The resulting radiopeptides were obtained with radiochemical purities greater than 95-98%, radiochemical conversions above 95%, and radiochemical yields ranging from 48-76%.

### *In vivo* PET experiments

**Animal models:** All animal studies were conducted according to the guidelines of the Canadian Council on Animal Care (CCAC) and approved by the Cross Cancer Institute Animal Care Committee (ACC) with protocol number AC 25277. *In vivo* PET experiments were performed using NIH-III mice (body weight: 20-24 g, Charles River Laboratories, Saint-Constant, QC, Canada). For tumor xenografts, approximately 5 × 10^6^ JIMT-1 cells (ATCC CRL-3793) suspended in 100 µL of a 1:1 mixture of phosphate-buffered saline (PBS) and Matrigel (Corning, Tewksbury, MA, USA) were injected subcutaneously into the upper left shoulder. JIMT-1 is a HER2-overexpressing, trastuzumab-resistant human breast cancer cell line; as it does not require estrogen receptor (ER) stimulation for growth, no hormonal supplementation was necessary. Tumor-bearing mice were used for PET experiments approximately 3-5 weeks after cell injection, once tumors reached volumes of approximately 250-300 mm^3^.

**Dynamic PET imaging:** PET imaging of the ^68^Ga-labeled radiopeptides was performed on an INVEON® PET/CT scanner (Siemens Preclinical Solutions, Knoxville, TN, USA). Prior to radiotracer injection, mice were anesthetized by inhalation of isoflurane in 40% oxygen/60% nitrogen (gas flow 1 L/min), and body temperature was maintained at 37 °C. Mice were positioned prone in the center of the field of view. A transmission scan for attenuation correction was not acquired.

Mice were injected intravenously via a tail vein catheter with 4-8 MBq of the respective radiopeptide in 100-150 μL saline. For blocking studies, 200 µg (8 mg/kg) of BMS-986189 was administered intravenously ~5 min before injection of the radiopeptide. Data acquisition was performed over 60 min in 3D list mode. The dynamic list-mode data were sorted into sinograms with 54 time frames (10 × 2 s, 8 × 5 s, 6 × 10 s, 6 × 20 s, 8 × 60 s, 10 × 120 s, and 6 × 300 s). Images were reconstructed using the maximum a posteriori (MAP) algorithm. No correction for partial-volume effects was applied.

Image files were processed using ROVER v2.0.51 software (ABX GmbH, Radeberg, Germany). Three-dimensional regions of interest (ROIs) were defined by thresholding and encompassed the entire visible tumor volume. Thresholds were defined at 50% of the maximum radioactivity uptake within the tumor tissue. Mean standardized uptake values [SUVmean = (activity/mL tissue)/(injected activity/body weight), mL/g] were calculated for each ROI, and time–activity curves (TACs) were generated using GraphPad Prism 9.3.1 (GraphPad Software, San Diego, CA). All semi-quantitative PET data are presented as mean ± SEM from n measured mice.

### Expression and purification of PD-L1 for crystallization

Human PD-L1 (residues 19–239) was codon-optimized for expression in Escherichia coli and cloned into pET303/CT-His (GeneArt) between the XbaI and XhoI restriction sites. The construct was transformed into One Shot BL21 Star (DE3) cells and expressed in 10 × 1 L cultures of Terrific Broth supplemented with 100 μg/mL carbenicillin and antifoam 204. Cultures were grown at 37 °C to an OD600 of ~2.0, induced with 0.5 mM IPTG, and incubated at 20 °C for 18 h. Cells were harvested by centrifugation and stored at −80 °C.

Cell pellets were resuspended in lysis buffer containing 50 mM Tris-HCl (pH 8.0), 100 mM NaCl, 0.5% Triton X-100, 0.5% sodium deoxycholate, 10 mM β-mercaptoethanol, 1 mM EDTA, and 12.5% glycerol, supplemented with EDTA-free protease inhibitors, lysozyme, and Benzonase. Cells were lysed by sonication and clarified by centrifugation at 27,000 × g for 15 min. Inclusion bodies were collected, washed with 50 mM Tris-HCl (pH 8.5), 200 mM NaCl, 10 mM EDTA, and 10 mM β-mercaptoethanol, and subsequently solubilized in the same buffer containing 6 M guanidine hydrochloride. Insoluble material was removed by centrifugation at 75,000 × g for 1 h.

The denatured protein was refolded by dropwise dilution into refolding buffer containing 100 mM Tris-HCl (pH 8.5), 1 M L-arginine hydrochloride, 0.25 mM oxidized glutathione, and 0.25 mM reduced glutathione, followed by overnight incubation at 4 °C. The refolded sample was concentrated using a 5 kDa MWCO Vivaflow 200 tangential-flow filtration cassette and dialyzed overnight against 10 mM Tris-HCl (pH 8.5) and 20 mM NaCl.

Refolded PD-L1 was purified by anion-exchange chromatography on Capto DEAE resin using a 20 mM–1 M NaCl gradient, followed by size-exclusion chromatography on a Superdex 200 26/900 column equilibrated in 20 mM Tris-HCl (pH 8.5) and 100 mM NaCl. Peak fractions were pooled, concentrated to 15.4 mg/mL, flash-frozen, and stored at −80 °C until use.

### Crystallization and structure determination of PD-L1 in complex with 10002

For co-crystallization, PD-L1 was mixed with compound 10002 at final concentrations of 7.5 mg/mL and 1.5 mM, respectively (1:5 protein-to-ligand molar ratio). The mixture was incubated overnight at 4 °C and clarified by centrifugation prior to crystallization.

Crystallization trials were performed at 20 °C using the sitting-drop vapor diffusion method. Initial crystals were obtained in Morpheus III (Molecular Dimensions) condition A5 containing 1.6% dipeptide mix, 0.1 M Buffer System 2 (pH 7.5), and 30% Precipitant Mix 3. Crystals appeared after 8–12 weeks and diffracted to >6 Å resolution. Crystal quality was improved through optimization of precipitant concentration and pH, followed by supplementation with Silver Bullets (Hampton Research) B9 additive solution containing 0.025% (w/v) hexamminecobalt(III) chloride, salicylamide, sulfanilamide, and vanillic acid, and 2 mM HEPES (pH 6.8).

Crystals were transferred to reservoir solution supplemented with 25% glycerol and 1.5 mM compound 10002, and flash-cooled in liquid nitrogen. X-ray diffraction data were collected at the FMX beamline at NSLS-II using an Eiger X 9M detector. The PD-L1–10002 complex crystallized in space group C222_1_ and diffracted to 2.78 Å resolution.

Diffraction data were processed with XDS. Initial phases were obtained by molecular replacement using MoRDa (CCP4) with the PD-L1 structure (PDB ID: 4Z18) as the search model. The resulting structure was refined through iterative cycles of model building and refinement using PHENIX.

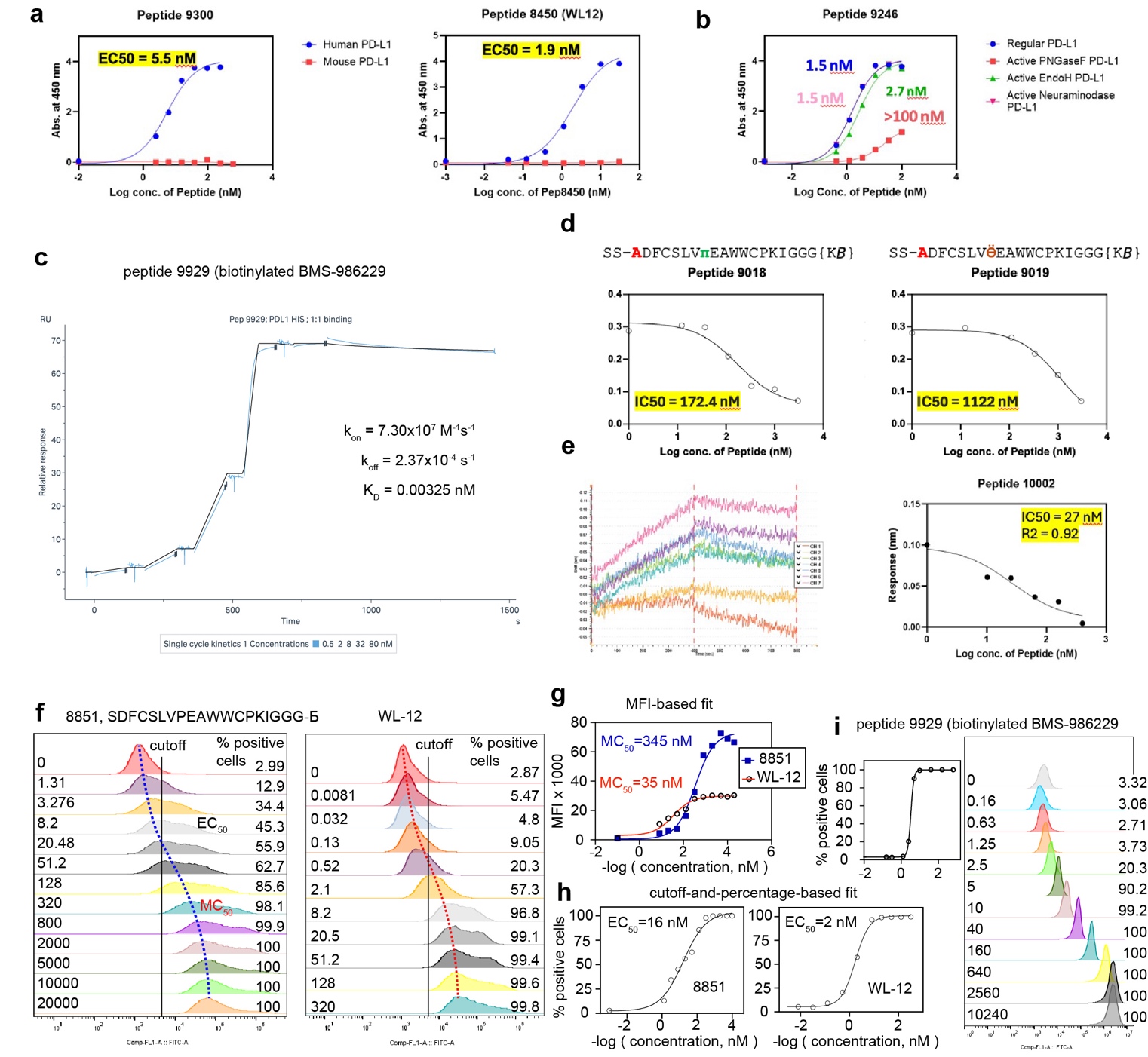

### Supplementary Figure 1: Summary of four major assays employed in this report

**a,** Representative curves from ELISA experiment which employed binding of macrocycles with C-terminal biotin to PD-L1 immobilized on the surface of microtiter well plates. These two experiments measure the binding of a representative macrocycle 9300 ADFCSLVπEAWWCQEIGGGK(ε−biotin) and a control compound, biotinylated WL-12, to plates coated with human PD-L1 and mouse PD-L1 (n=4 for each concentration, average shown). Based on this experiment we concluded that VPEA-family, just like WL-12 have exclusive binding to PD-L1 and no cross-reactivity with mouse homologue.

**b,** We employed ELISA assay with biotinylated macrocycle 9246 to test whether glycosylation of PD-L1 protein is critical for binding to VPEA-family macrocycles. We observed that complete removal of N-linked glycans by PNGaseF treatment was detrimental to binding whereas treatment with neuraminidase and EndoH had no effect (*continued next page*)

**c,** Screenshot of a calibration experiment describing surface plasmon (SPR) measurement of the kinetics of binding of compound 9929, biotinylated BMS-986229 macrocycle, to PD-L1 immobilized on the surface of a CM5 chip.

**d,** Examples of a readout from biolayer interferometry (BLI, GatorBio) instrument to measure the ability of soluble peptide macrocycles to block the interactions between PD-L1 immobilized on the BLI tip and its native ligand PD-1 present at a constant concentration (50 nM). Decrease in the response units with increasing concentration of the competitor in solution indicates blocking of all PD-L1 proteins by the macrocycle. The illustrated example shows the effect of simple epimerization leading to factor of 6.5 improvement in IC_50_ values for compound **9018** with (4S)-4-fluoro-ʟ-proline in position 8 when compared to the epimer compounds **9019** with (4R)-4-fluoro-ʟ-proline in the same position.

**e,** We used the BLI instrument to measure the binding of soluble PD-L1 protein to biotinylated peptide immobilized on the BLI tip. For example, this assay describes binding of PD-L1 protein used for crystallisation studies to biotinylated WL-12 peptide immobilized on the BLI tip (red curve). Titration of soluble macrocycle **10002** employed in crystallization studies progressively diminished the signal. This experiment indicated that **10002** binds to PD-L1 in the same binding pocket as WL-12.

**f,** flow cytometry and dose-response titration of macrocycles with C-terminal biotin to PD-L1(+) cells incubated on ice, followed by the readout of the signal with fluorescently labeled streptavidin. Measurement of the labeled population by flow-cytometry (FACS) produces dose-response curves that can be processed using two different analyses. One strategy employs tracking of the median or mean fluorescent intensity (MFI) of the population illustrated by blue and red dashed lines. The second strategy employs measurement of fraction of fluorescently labeled cells above constant cutoff.

**g,** Fit of “MFI vs. concentration” scatter to a hyperbolic curve can determine the concentration at which the intensity reaches the half-maximum response (MC_50_). The problem with this analysis is the instability of the hyperbolic fit with increasing peptide concentration. The titration of peptides on cells rarely reaches clear saturation: increased concentration always leads to increased signal leading to the drift of the upper bound of the hyperbolic curve. As a result, the MC_50_ values from hyperbolic fit also drift as the upper bound of the concentration series in the titration steadily increases.

**h,** Illustration of the EC_50_ calculation method based on a constant threshold (black line in **f**) and determining the % of cells above this threshold at various concentrations. The hyperbolic fit has a clear, stable saturation plateau at 100% and it indicates that at 16 and 2 nM concentrations of **8851** and **WL-12**, 50% of the cells are labeled by these ligands above the threshold. This fit is stable for any number of datapoints because has a well-defined upper bound (100% at C🡪 infinity) and predetermined lowed bound (here 3% as C🡪0). In this publication we ubiquitously employ the EC_50_ in place of MC_50_. For side-by-side comparison of EC_~~50~~_ and MC_50_ see Supplementary Fig 5e, h.

**a**
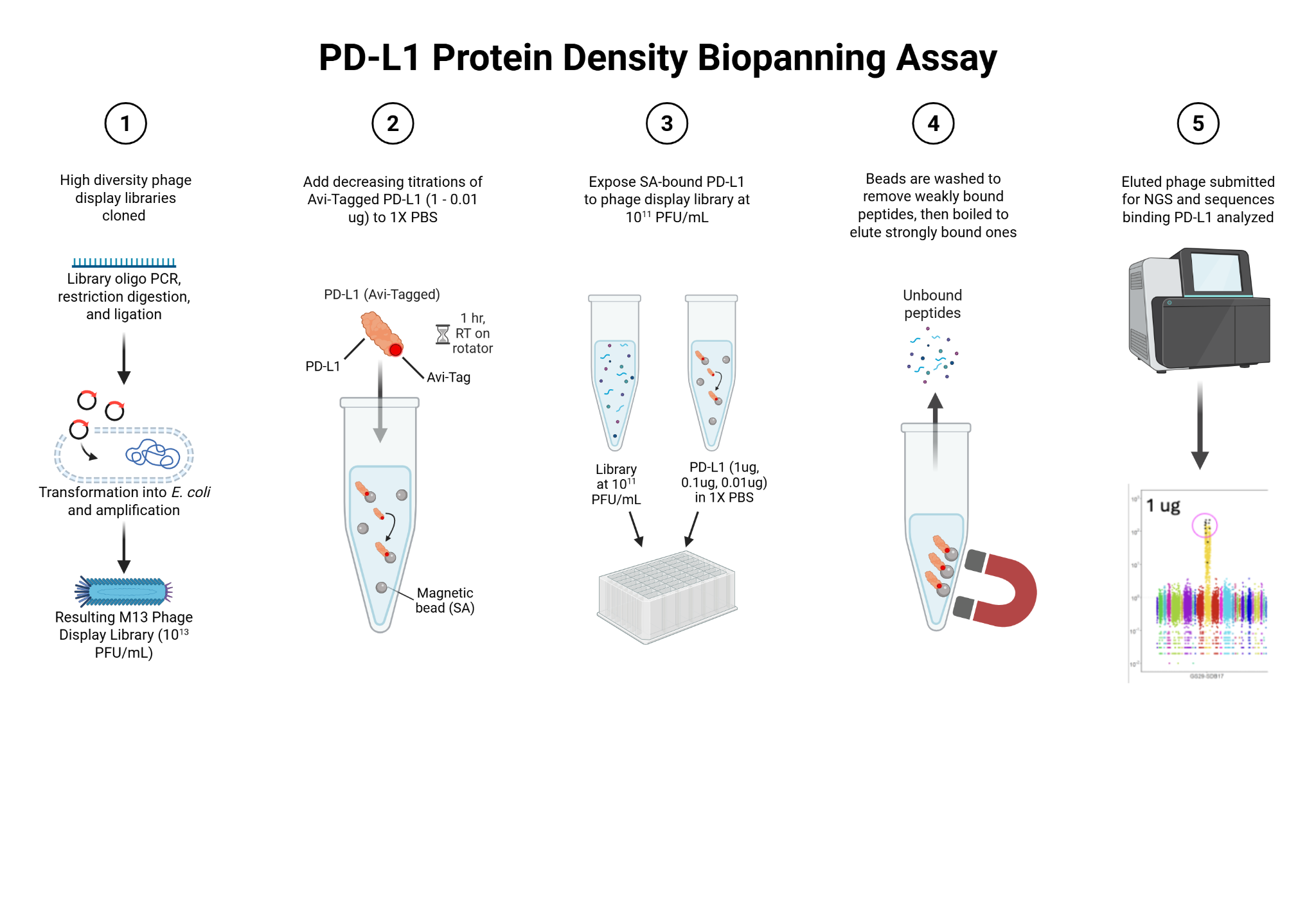

**b**

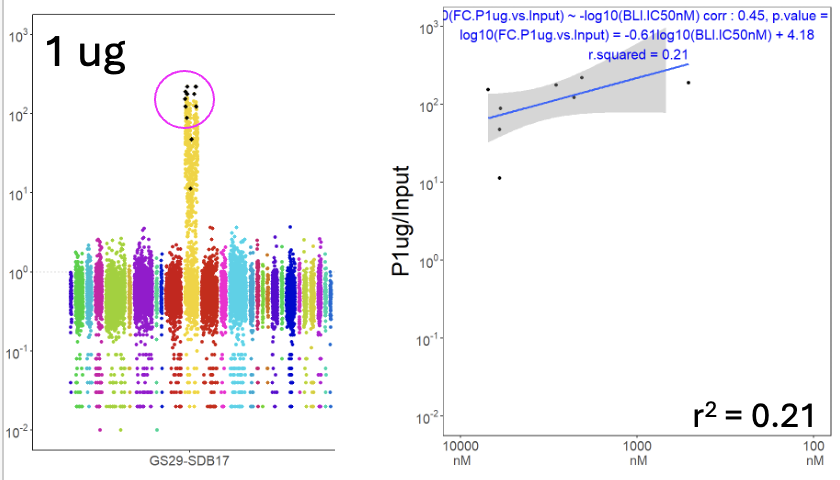

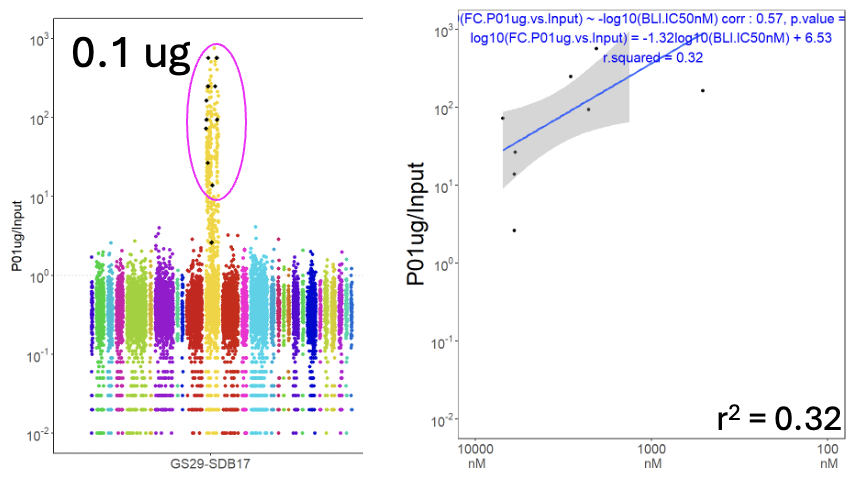

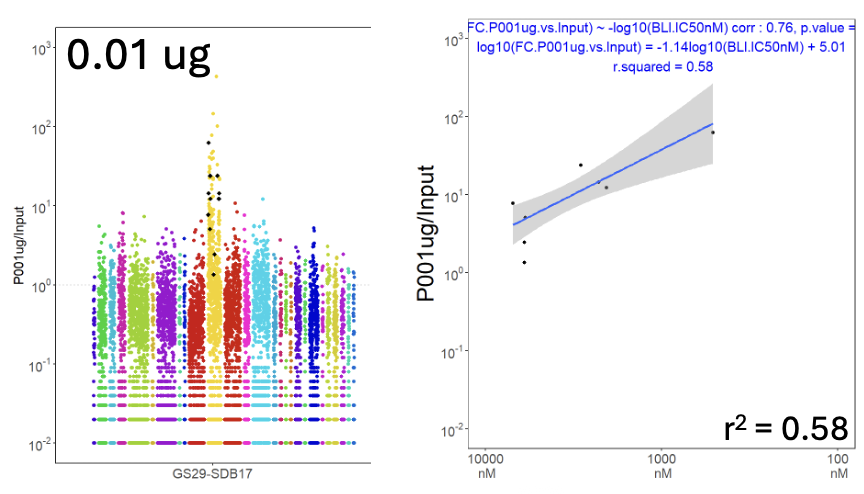

### Supplementary Figure 2: Production and panning of focused libraries

**a,** Phage display library was generated from SC1966-HF92:GenTitan Hi-Fi 92k single stranded DNA (ssDNA) pool that contains anywhere from 8,000 to 50,000 unique DNA sequences. PCR amplification of this ssDNA yields dsDNA for cloning (step 1). Subsequent restriction digestion, ligation into phage vector DNA and transformation into electrocompetent ER2738 *E. coli* yielded phage library which was amplified in *E. coli* and purified by PEG/NaCl precipitation. Step 2: In a typical focused library experiment, avi-tagged PD-L1 protein was diluted to a desired amount, 1ug, 0.1 ug, or 0.01 ug per “tube”, and all the protein was captured on streptavidin coated magnetic beads (SMB). Step 3: SMB-bound PD-L1 was mixed with focused library diluted to 10^11^ PFU/mL. The numbers 1, 0.1 and 0.01 ug represent the total amount of protein captured by a fixed number of SMB. Step 4: Bead-protein-phage mixture was washed by automated magnetic bead washer (KingFisher) to remove unbound phage. Boiling the beads in nuclease-free water elutes ssDNA of the phage. Step 5: Eluted ssDNA is converted by PCR to an Illumina-compatible dsDNA for next generation sequencing (NGS). Differential Enrichment (DE) analysis of the Illumina-NGS data yields fold-change enrichment factors (FC) for 8,000 to 50,000 peptides. **b,** Correlation between FC values observed in NGS analysis of the panning of focused library GS29 on PD-L1:SMB and IC_50_ of synthetic peptides measured by BLI. Higher PD-L1 density on SMB offers a wider dynamic range for FC. Lowest coating density of 0.01 ug total protein results in poor dynamic range of FC but the correlation between FC and IC_50_ has the widest dynamic range.

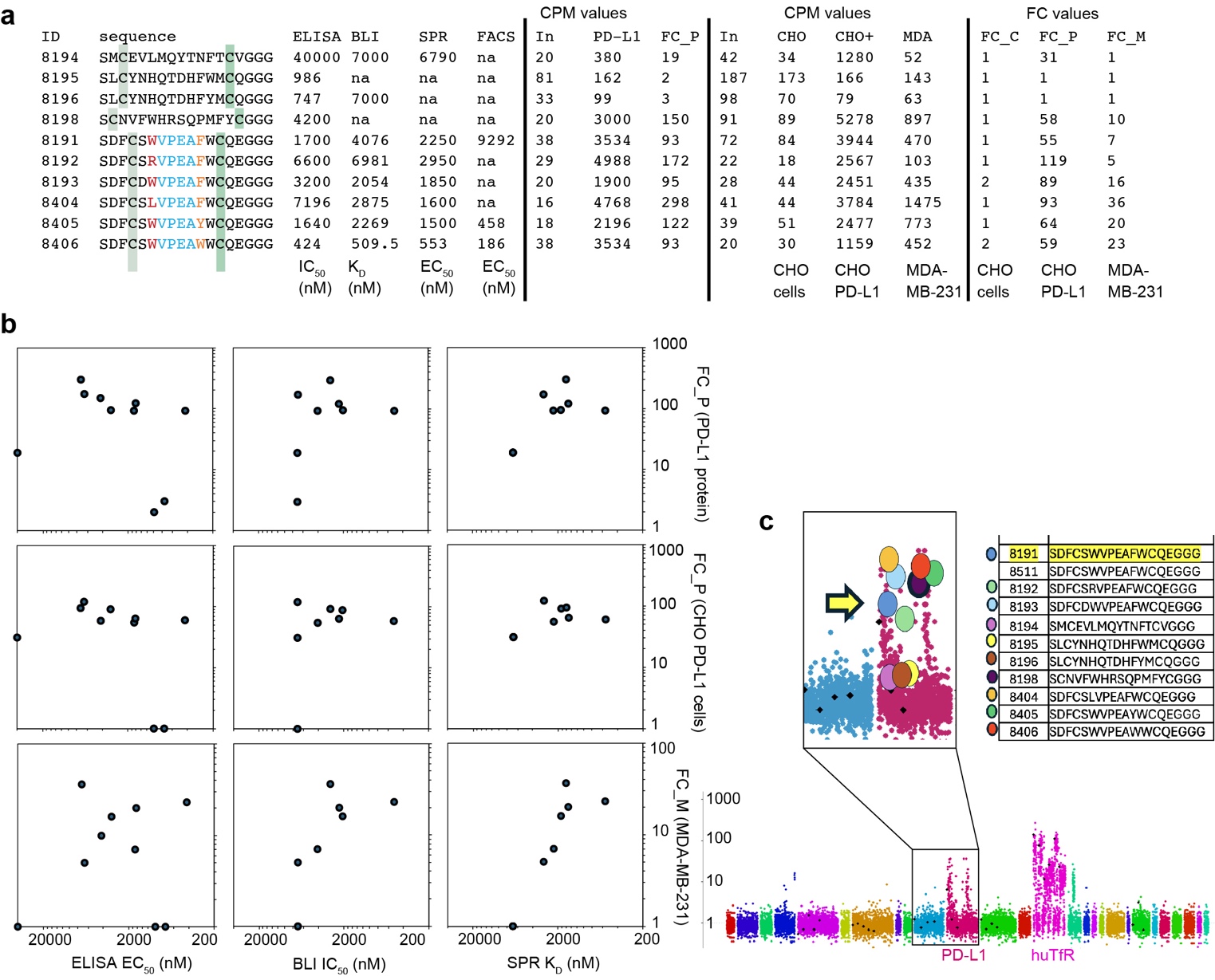

### Supplementary Figure 3: Calibration of focused library GS29 in panning on protein or cells

**a,** Summary of measurements of affinities and inhibitory activities for 10 synthetic macrocycle peptides across four assays—ELISA, BLI, SPR and FACS. The same table contains normalized NGS data—counts per million (CPM) values in the input and output of various panning experiments—and calculated fold-change enrichment (FC) for the same peptides displayed on phage, as part of focused library GS29 panned on four different targets. The panning targets were purified PD-L1 protein immobilized on streptavidin-coated magnetic beads (SMB:PD-L1, see Supplementary Fig. 1), negative CHO cells or CHO PD-L1(+) cells, and MDA-MB-231 cells. **b,** Scatter plots showing correlation between various values in table (a); FC values from panning on MDA-MB-231 cells show the best correlation with K_D_ values measured by SPR and IC_50_ values measured by BLI. In contrast, panning on CHO PD-L1(+) cells and SMB:PD-L1 correlate poorly with outcomes in all assays and show little distinction between low-micromolar and mid-nanomolar binders. **c,** Manhattan plot describing panning of GS29 on MDA-MB-231 cells. The peptides from (**a**) are highlighted in the zoomed in portion of the plot. Peptides in a segment labeled “huTfR” are from panning on human transferrin receptor.

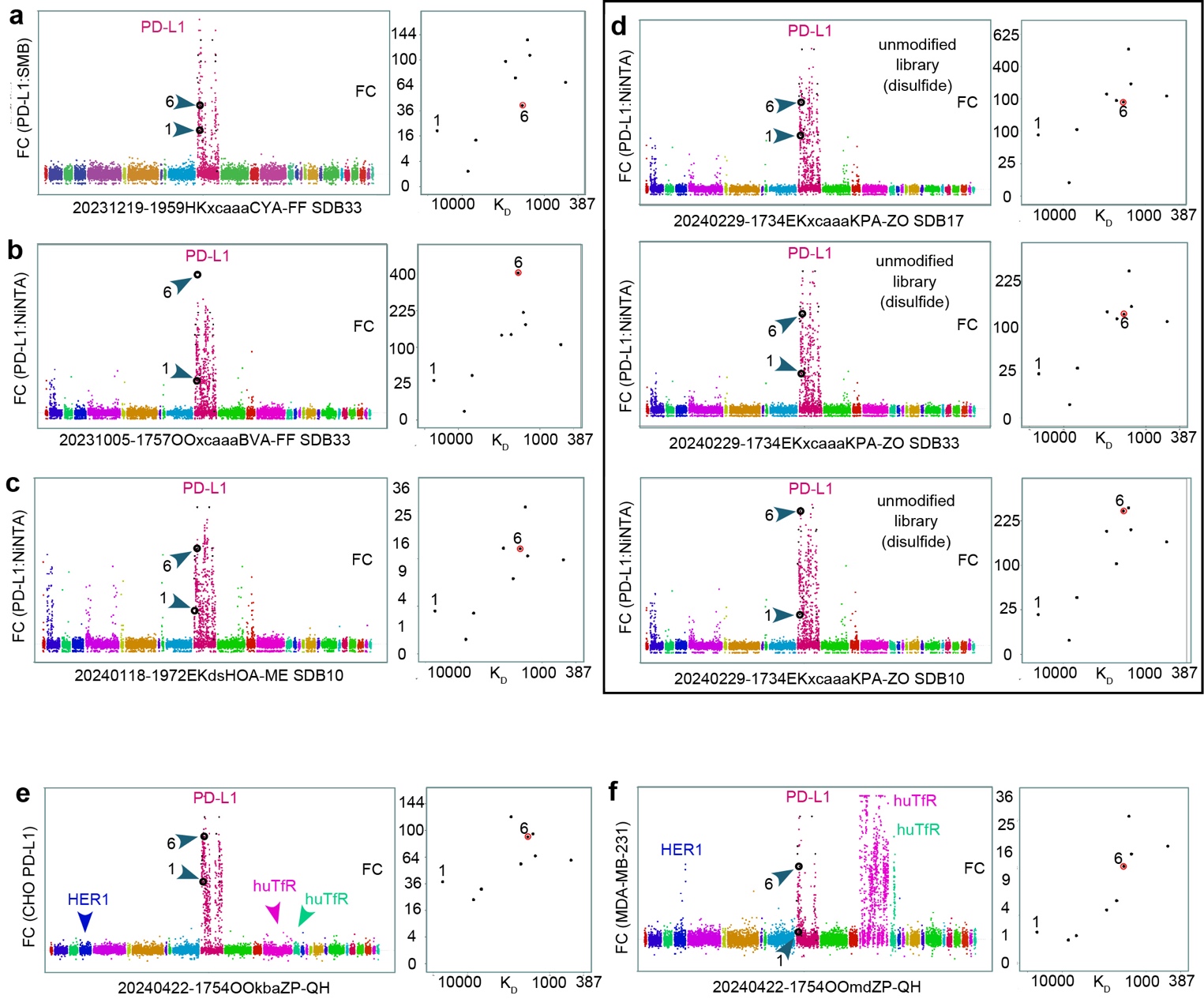

### Supplementary Figure 4: Reproducibility of calibration of GS29 in panning on protein and cells

**a,** Manhattan plot showing enrichment of GS29-SDB33 library in panning on SMB coated by PD-L1 protein with tandem hexa-histidine and Avi-tag (AcroBio: PD-L1 His/Avi; Cat# PD1-H82E5, Lot BV1044-96HF1-11S). The red scatter contains PD-L1 binding peptides whereas other segments of other color contain peptides that bind to other classes of proteins described in our previous report^3^. Unique alphanumeric ID for the NGS data is listed below the Manhattan plot and more information for this ID is available at <https://48hd.cloud/> . Manhattan plot is juxtaposed to a scatter plot with Y-axis synchronized to Manhattan plot and X-axis containing the K_D_ values for nine synthetic peptides as determined by SPR. Throughout this figure, we track two peptide sequences denoted as “1” and “6” with the highest and 6^th^ highest K_D_ value. **b,** Repeat of panning of the same GS29-SDB33 library against the same batch of HisAvi-PD-L1 protein immobilized on Ni:NTA beads shows noticeable binding of peptides from non-PD-L1 segments because these peptides have affinity for Ni:NTA beads. **c,** Repeat of the exact panning conditions on PD-L1:NiNTA as in **b** but performed three months later, by a different user, using a different batch of GS29 cloned into SDB10 vector. **d,** Panning of a mixture of three GS29 libraries encoded by SDB10, SDB11 and SDB33 silent codons against PD-L1 on Ni:NTA beads. The Manhattan plots and calibration curves are similar however performances of individual peptides (e.g., 6) are fluctuating. Comparison within **d** shows fluctuations between different batches of library in the same solution, in the same experiment, whereas comparison between **b**, **e**, and horizontally aligned Manhattan plots in **d** offers comparison of the same batches of GS29 in panning experiments performed on different days, by three different users, and sequenced in different NGS batches. Overall, the GS29 panning on PD-L1 protein is stable with respect to user, library age, library batch, and sequencing batch. Minor fluctuations are unavoidable even when chemically identical libraries are panned at the same time, in the same solution and sequenced in the same batch.

**e,** Panning of GS29 library against CHO cells that overexpress PD-L1 protein and **f,** panning of the same library against MDA-MB-231 cells that have lower level of expression of PD-L1. Cells with lower level of PD-L1 expression offer a calibration curve with a wider dynamic range. Peptides in the purple and green segments labeled “huTfR” are from panning on human transferrin receptor, which is expressed on MDA-MB-231 cells but not on CHO cells. Similarly, peptides from blue segment labeled HER1 are from panning on HER1/EGFR protein and five of the peptides from this segment bind to HER1 expressed on MDA-MB-231 cells; these peptides do not bind to CHO cells. Note that peptides in blue (HER1), purple and green (huTfR) segments are silent in all panning experiments that use purified PD-L1 protein (**a**-**d**)

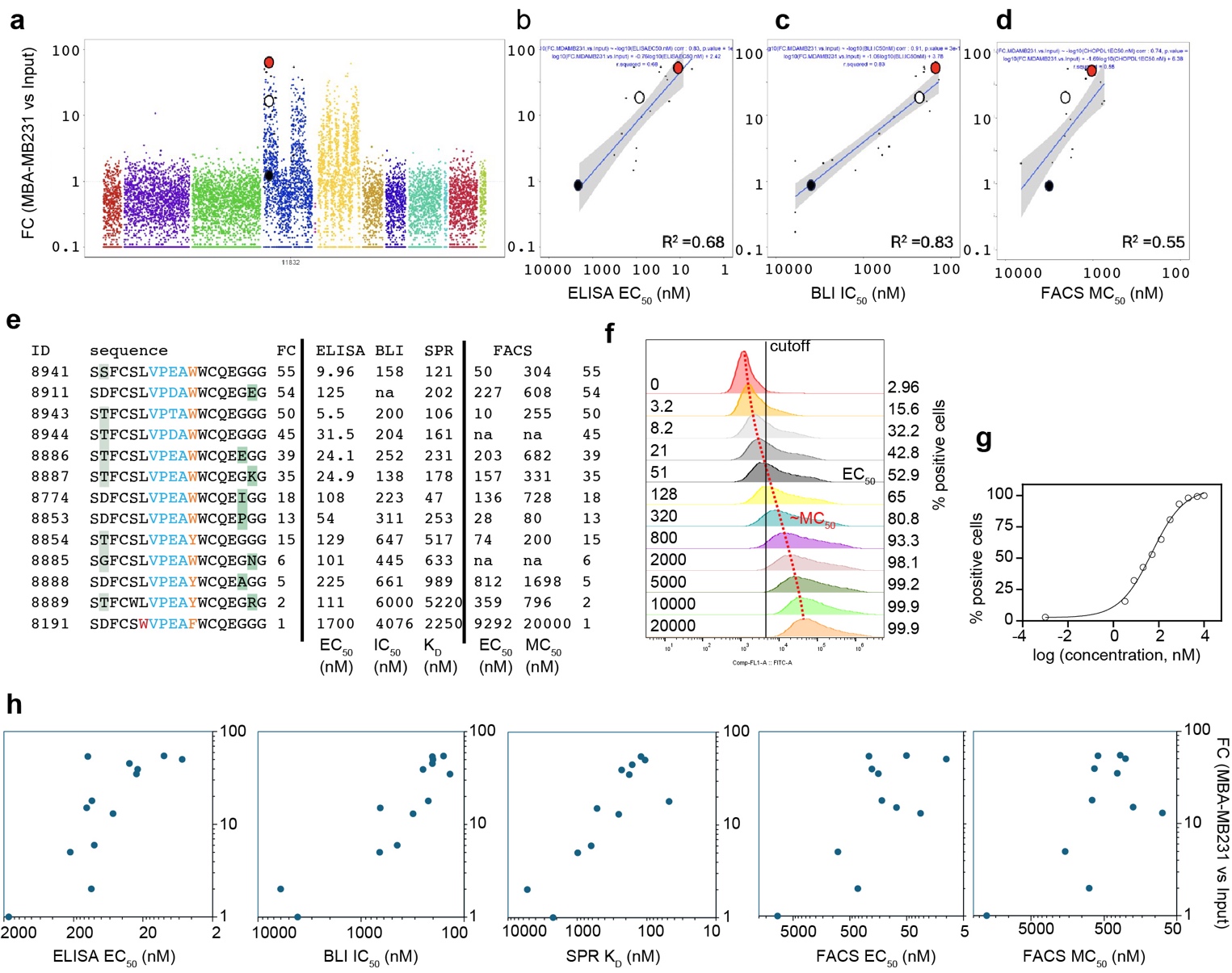

### Supplementary Figure 5: Calibration of focused library GS32 panned on MDA-MB-231 cells

**a,** GS32 library displayed on phage contains a parent sequence SDFCSWVPEAFWCQE (8191) and scans and mutations in positions 2, 5, 6, 9, 11, 16, 17, a total of 1857 PD-L1 sequences and 9965 sequences not related to PD-L1 for a total of 11,822 sequence in the library. The library was panned on MDA-MB-231 cells and the eluted ssDNA was analyzed by Illumina NGS and DE-analysis to yield FC values for the 11,822 sequences (each dot on the Manhattan plot represents a unique peptide sequence) **b-d,** Based on GS32 results, 15 peptides were made and tested in three different assays. Plots describe the correlation of FC-values with EC50 values measured by ELISA (**b**), IC50 values measured by BLI (**c**) and FACS MC50 the half-saturation values measured by flow-cytometry (**d**). e, A summary of 13 peptides, their FC values and activities. f, Flow cytometry data describing binding of peptide 8941 to CHO PD-L1 cells.

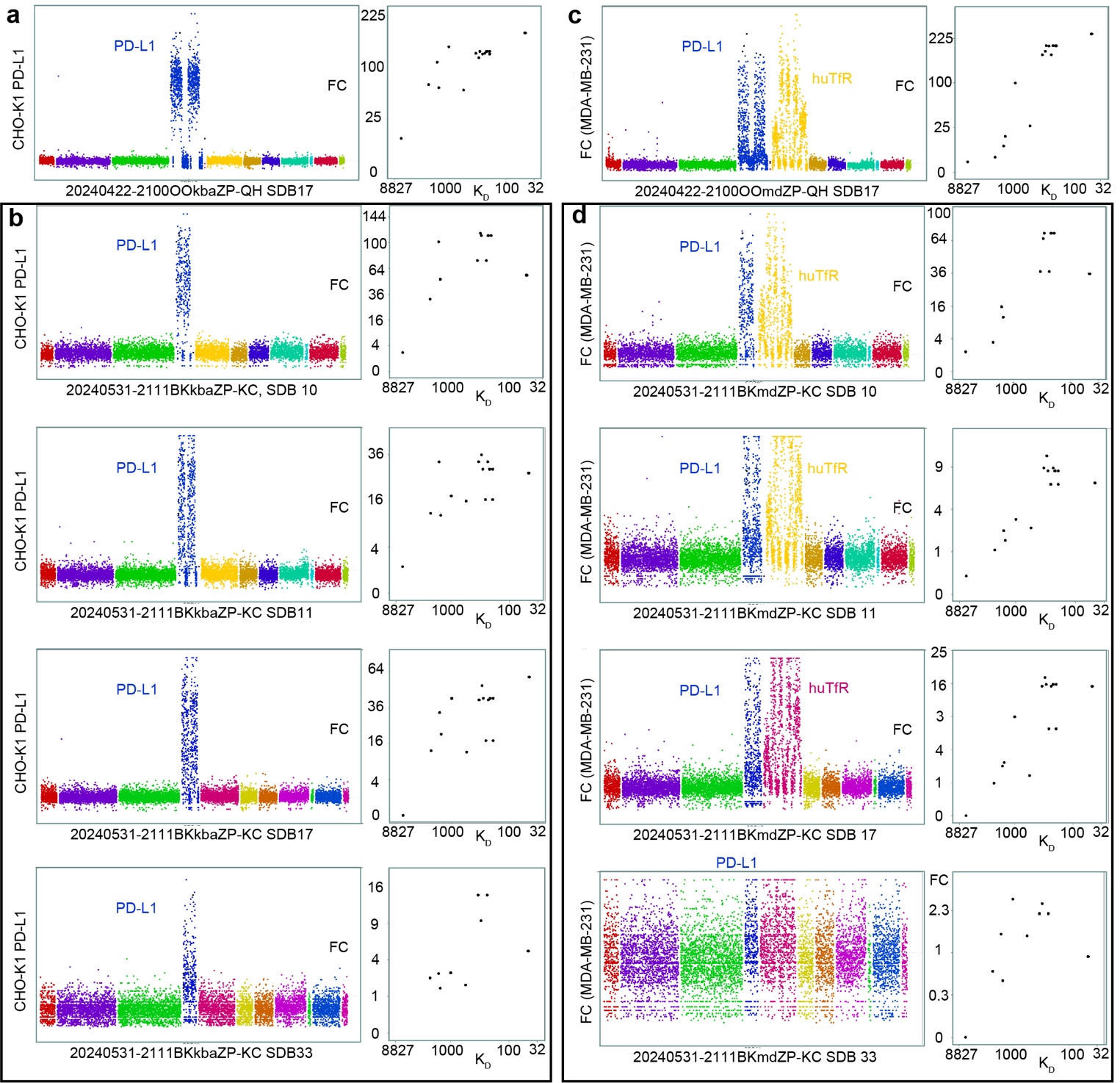

### Supplementary Figure 6: Reproducibility of calibration of phage library GS32 in panning on cells

**a,** Manhattan plot showing fold change (FC) enrichment of GS32 in panning on PD-L1(+) CHO cells and its calibration using K_D_ of synthetic peptides determined by SPR. **b**, Four Manhattan plots from panning on PD-L1(+) CHO cells performed on a different day by a different user (*rmistry*) using a mixture of four batches of GS32 library encoded by SDB10, SDB11, SDB17 and SDB33 silent barcodes. **c,** Panning of the same GS32 batch performed by the same user as in **a** (*gsharma*) but using MDA-MB-231 cells. As in other experiments (Supplementary Fig. 4f), we observed that peptides in a segment originating from panning on human transferrin receptor (huTfR) bind to MDA-MB-231 cells; huTfR is expressed on MDA-MB-231 cells but not on CHO cells. **d**, Four Manhattan plots from the same MDA-MB-231 cells experiment performed on a different day by a different user (*rmistry*) using a mixture of four batches of GS32 library encoded by SDB10, SDB11, SDB17 and SDB33 silent barcodes. Horizontal comparison between a, b and c, d shows the outcomes from the same batches of GS32 in two parallel experiments performed by the same user, on the same day on two different cell lines.

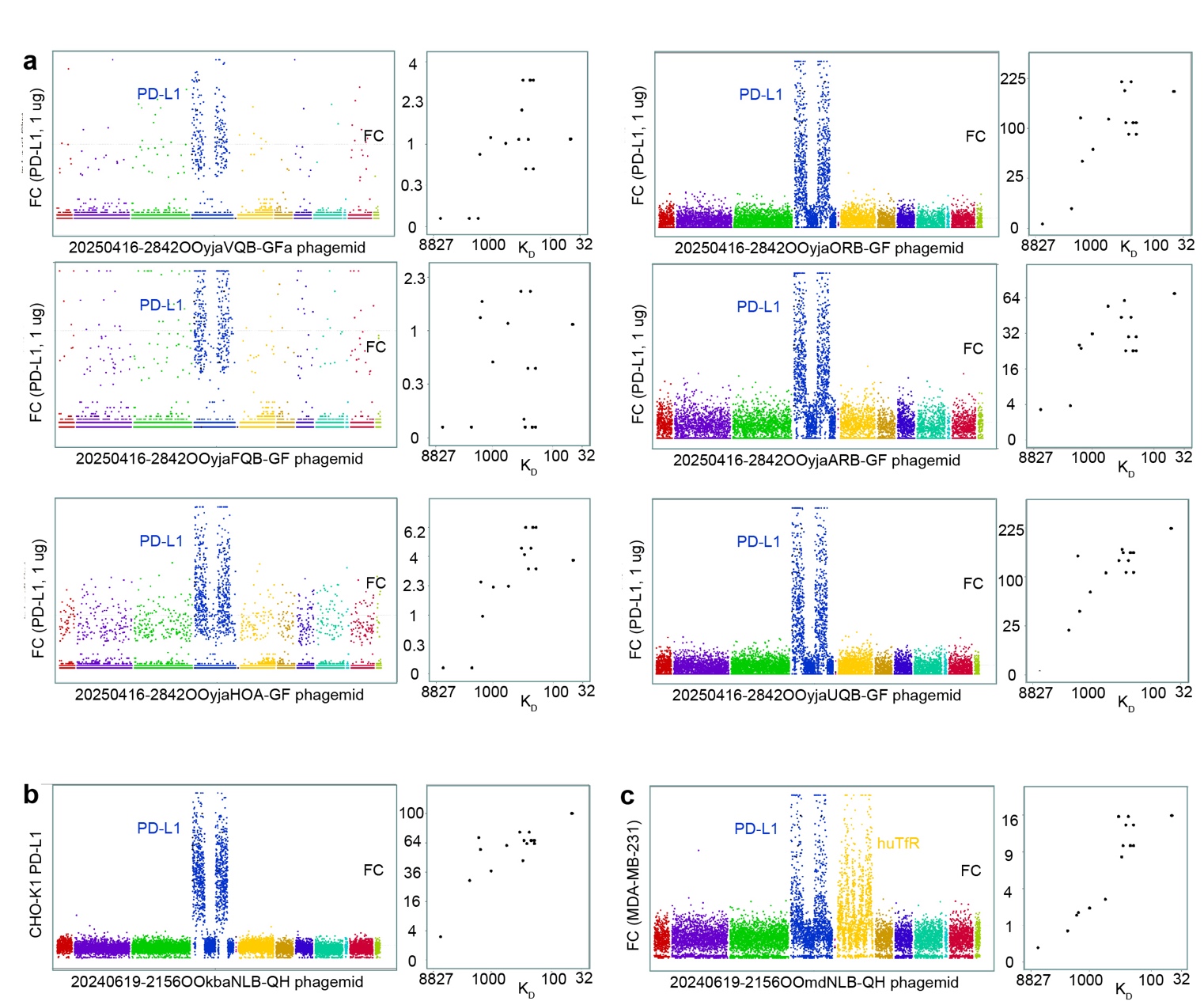

### Supplementary Figure 7: Calibration of phagemid library GS32 in panning on PD-L1 and cells

**a,** Manhattan plot showing fold change (FC) enrichment of GS32 phagemid library in six different panning procedures that employ purified PD-L1 protein and its calibration using K_D_ of synthetic peptides determined by SPR. Experiments were performed by the same user in six different panning conditions that altered the PD-L1 binding time, number and duration of washes and other variables. **b,** Manhattan plot showing FC in panning on PD-L1(+) CHO cells and its calibration using SPR-K_D_. **c,** Manhattan plot showing FC in panning performed by the same user as in **b** (*gsharma*) using MDA-MB-231 cells and its calibration using SPR-K_D_. As in other experiments (Supplementary Fig. 4f, 6c-d), we observed that peptides in a yellow segment originating from panning on human transferrin receptor (huTfR) bind to MDA-MB-231 cells because huTfR is expressed on MDA-MB-231 cells but not on CHO cells.

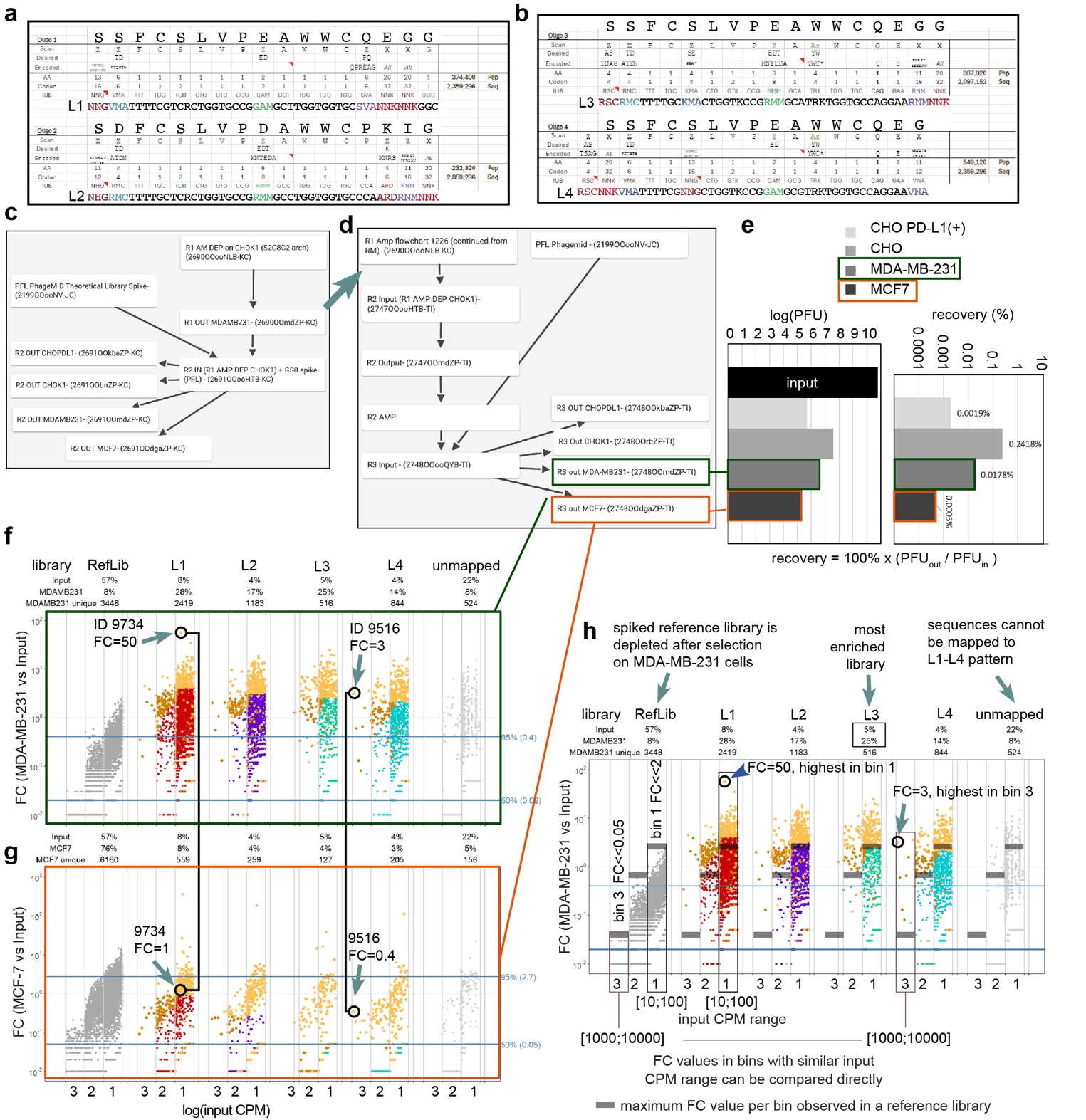

### Supplementary Figure 8: Design and cell-based panning of affinity maturation libraries

**a,** Design of oligonucleotides L1 and L2 based on two parent sequences 8915 and 8851. Colored trinucleotides in positions 1, 2, 9, 14, 15, 16 encode a predetermined subset of amino acids. **b,** Oligonucleotides L3 and L4 further randomize parent sequence 8915 using two additional randomization strategies. L1-L4 were designed to contain ~2 million unique nucleotide sequences and 250-500K unique peptide sequences. The oligonucleotides L1-L4 were cloned into the phagemid vector to yield affinity maturation (AffMat) library. **c,** Flow chart of the 2-round AffMat campaign that starts from panning on MDA-MB-231 cells, followed by panning on MDA-MB-231, MCF-7, CHO PD-L1(+) and control CHO cell lines. **d,** Flow chart of 3-round AffMat campaign which recycles R1 output from the previous campaign followed by two additional rounds of selection. Each box in the flow-chart represents an NGS dataset. Arrows indicate relations between these NGS datasets. Datasets in red and green frames are analyzed in panels (**e-g**).

**e,** Quantification of the recovery of the phage at the round 3 of the selection (see **d**) was performed by qPCR analysis of the input and output phagemid. There is 10-100x enrichment on PD-L1(+) cels when compared to a PD-L1(-) control.

**f-g**, Manhattan plot describing fold change (FC) enrichment of sequences in panning of R3 library on MDA-MB-231 cells (**f**) or MCF-7 cells (**g**). FC was calculated using differential enrichment (DE) pipeline (see methods) from output CPM values observed in n = 4 independent experiment using input CPM values as reference. Each dot represents a unique peptide sequence. Sequences are grouped by library type: L1-L4, spiked reference library and unmapped sequences. Within each library, the sequences are sorted horizontally based on their normalized input (CPM). Top of the plot contains the number of unique sequences observed in the input of each library, and fraction of each library in the output and input. Gold and copper color designate auto-generated list of hit sequences. Two hit sequences are highlighted by black circle; they correspond to synthetic peptides 9734 and 9516. The 9516 is one of the most abundant sequences and it exhibits a modest enrichment of FC=3 whereas 9734 is significantly lower abundance and it exhibits the highest enrichment FC=50. Vertical lines in **f** and **g**, illustrate that both sequences have higher enrichment on MDA-MB-231 cells when compared to MCF7 cells. To calibrate the FC values for sequences of high and low abundance, we employ enrichments of the reference library as a baseline.

**h,** Expanded NGS analysis that compares enrichment of each sequence to the baseline-level enrichment of the sequences in the reference libraries. Binning of the sequences in each library based on their input values creates 3-4 distinct bins. Sequences that have similar normalized input frequency (CPM) can be compared across different libraries. Grey bars denote the maximum FC values observed in each reference bin; the same values are mirrored to 5 other libraries. Presence of baseline clearly illustrates that enrichment of sequence 9516 (FC=3) is substantial because the baseline enrichment in the same bin is FC<0.05. For low abundance sequence 9734 the FC=50 the enrichment should be compared to a baseline response of FC=2. The cell-binding activity of synthetic 9516 and 9734 are EC_50_=4 nM and EC_50_=8 nM respectively. Subsequent figure expands this analysis to two AffMat campaigns and 4 cells and it connects the AffMat abundances and enrichments to the activity measurements for synthetic macrocycles

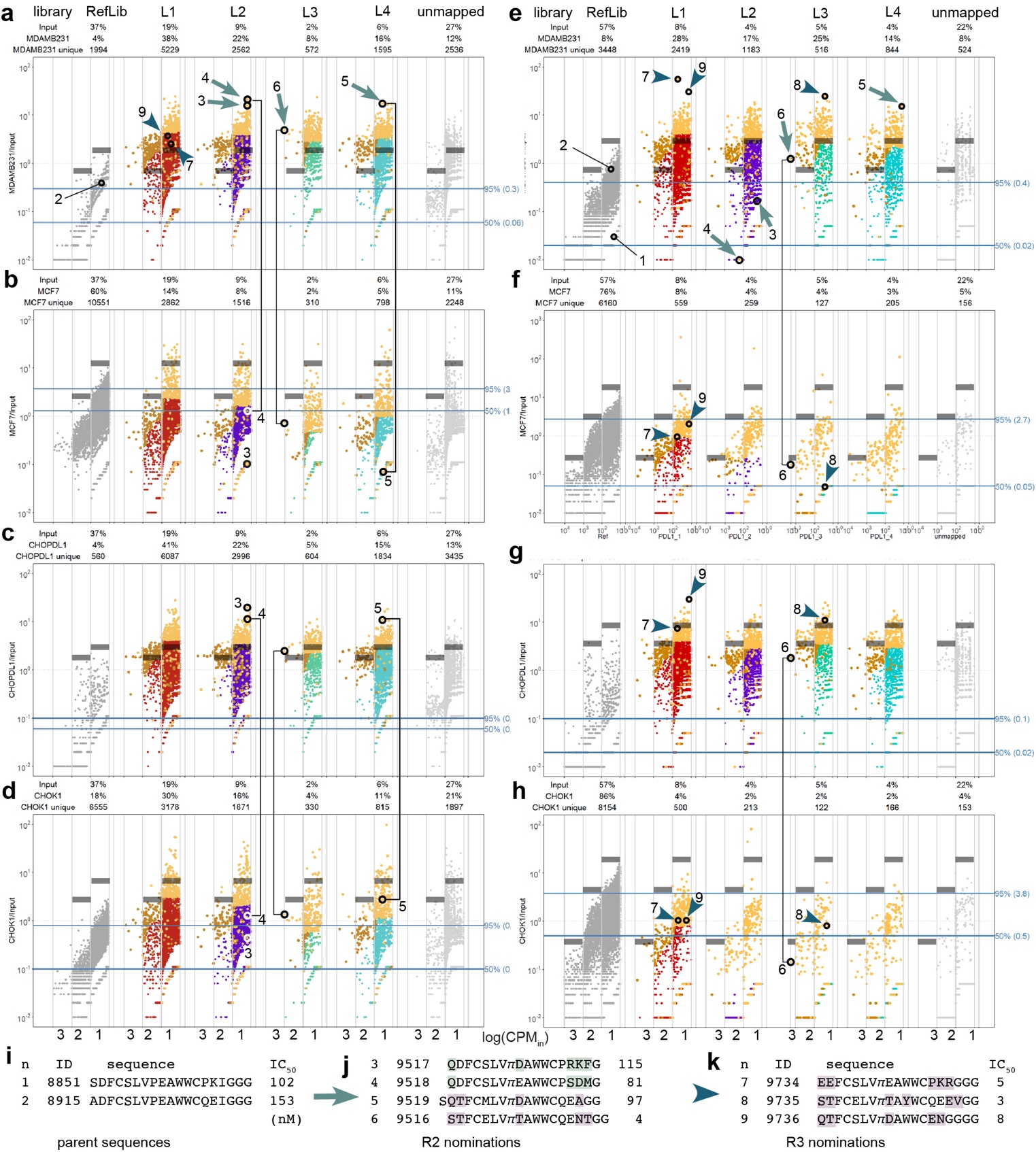

### Supplementary Figure 9: Concerted NGS analysis of eight AffMat datasets

**a-d,** Manhattan plots from 2-round AffMat campaign describing enrichment on MDA-MB-231 (**a**), MCF7 (**b**), CHO PD-L1(+) cells (**c**), and CHO cells (**d**) in the second round. **e-h,** Manhattan plots from 3-round AffMat campaign describing enrichment on MDA-MB-231 (**e**), MCF7 (**f**), CHO PD-L1(+) cells (**g**), and CHO cells (**h**) in the third round. The Manhattan plots in **a** and **e** track the enrichment of the two parent sequences numbered 1 and 2 (see panels **i** for sequences), four daughter sequences numbered 3-6 nominated in the R2 (see **j** for sequences), and three daughter sequences numbered 7-9 nominated in the R3 (see **k** for sequences). Sequences 3-6 are further tracked in plots b-d, and sequences 6-9 are tracked in plots **f**-**h**. As detailed in Supplementary Fig. 8, the grey bars denote the baseline-level enrichment of the control library spiked into a maturation library. Daughters nominated based on round 2 NGS analysis are marked by turquoise arrows and those nominated from round 3 NGS analysis are marked by teal arrowheads.

**i-k**, Sequences of the parent and daughter sequences as well as their cell-binding potency in flow-cytometry titration experiment (see main text for details). Changes in daughter sequences from PKI-family (3 and 4) are highlighted green; changes in daughters from QE-family are highlighted in purple.

Note that in **a** and **e**, the parent sequences do not enrich above the baseline population whereas all seven selected daughters exhibit above-the-baseline enrichment on PD-L1(+) cells and baseline-level enrichment on PD-L1(-) cells. Only daughter 6 exhibits consistent enrichment in rounds 2 and 3. Daughters 3-5 are not enriched in round 3 whereas daughters 7-9 are enriched above the baseline but their enrichment is only modest in round 2. Nominations based on round 3 appear to be more effective yielding single digit nanomolar cell-binding potency in 3 out of 3 tested cases, as opposed to 1 in 4 sequences from round 2 yielding similar performance. We note that the number of tested sequences is not sufficient to draw a statistically significant conclusion.

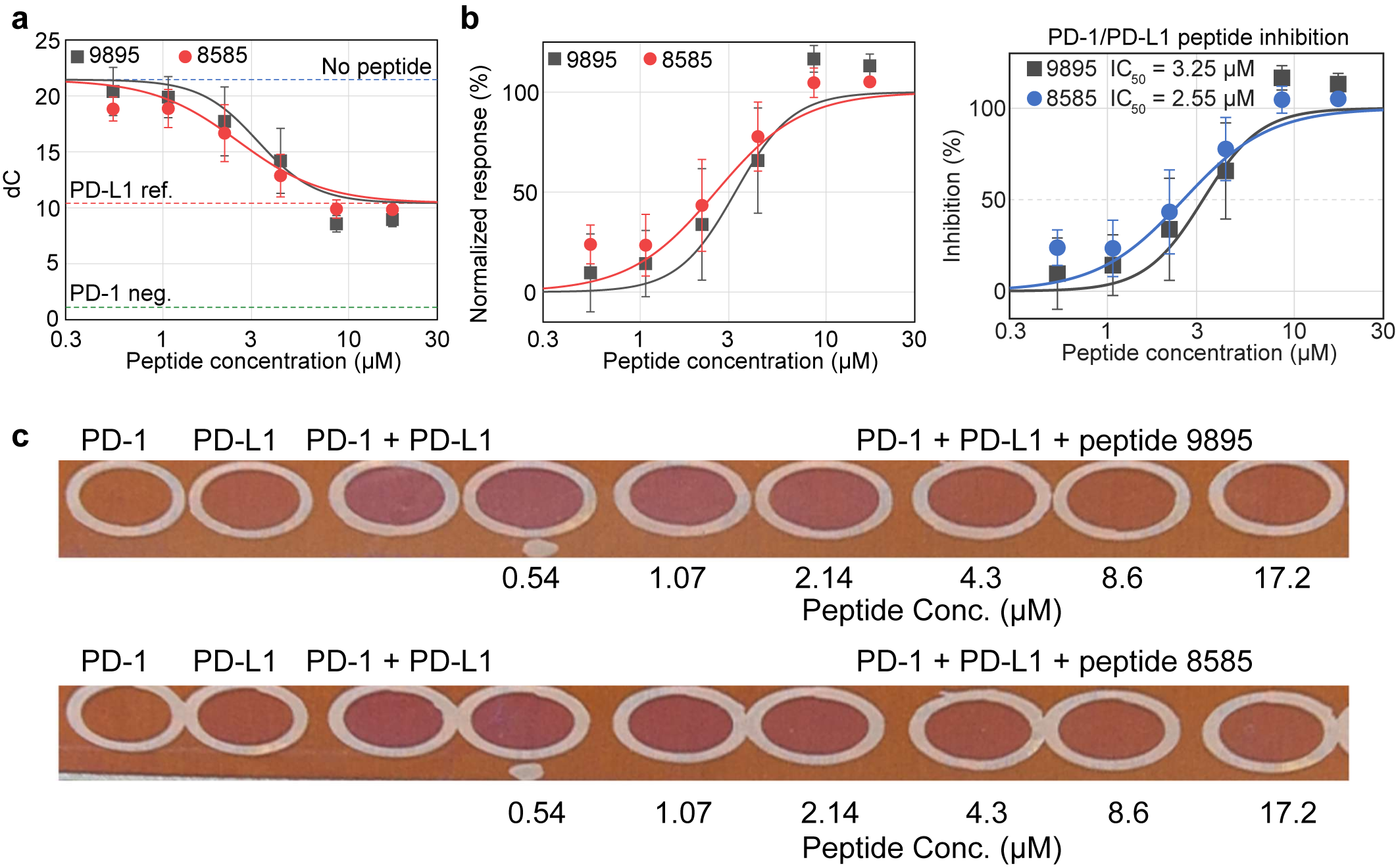

### Supplementary Figure 10: Inhibition of PD-1 and PD-L1 using VICA

**a,** Dose-dependent inhibition in solution measured as differential colour response (dC) of the inhibition of PD-1 and PD-L1 with peptides **9895** & **8585** at increasing concentrations. **b**, Normalized dose-response curves for peptides **9895** & **8585** as well as their relative IC_50_s. **c**, VICA biosensor colour response for PD-1/PD-L1 controls and increasing concentrations of peptides **9895** & **8585**.

**a	b**

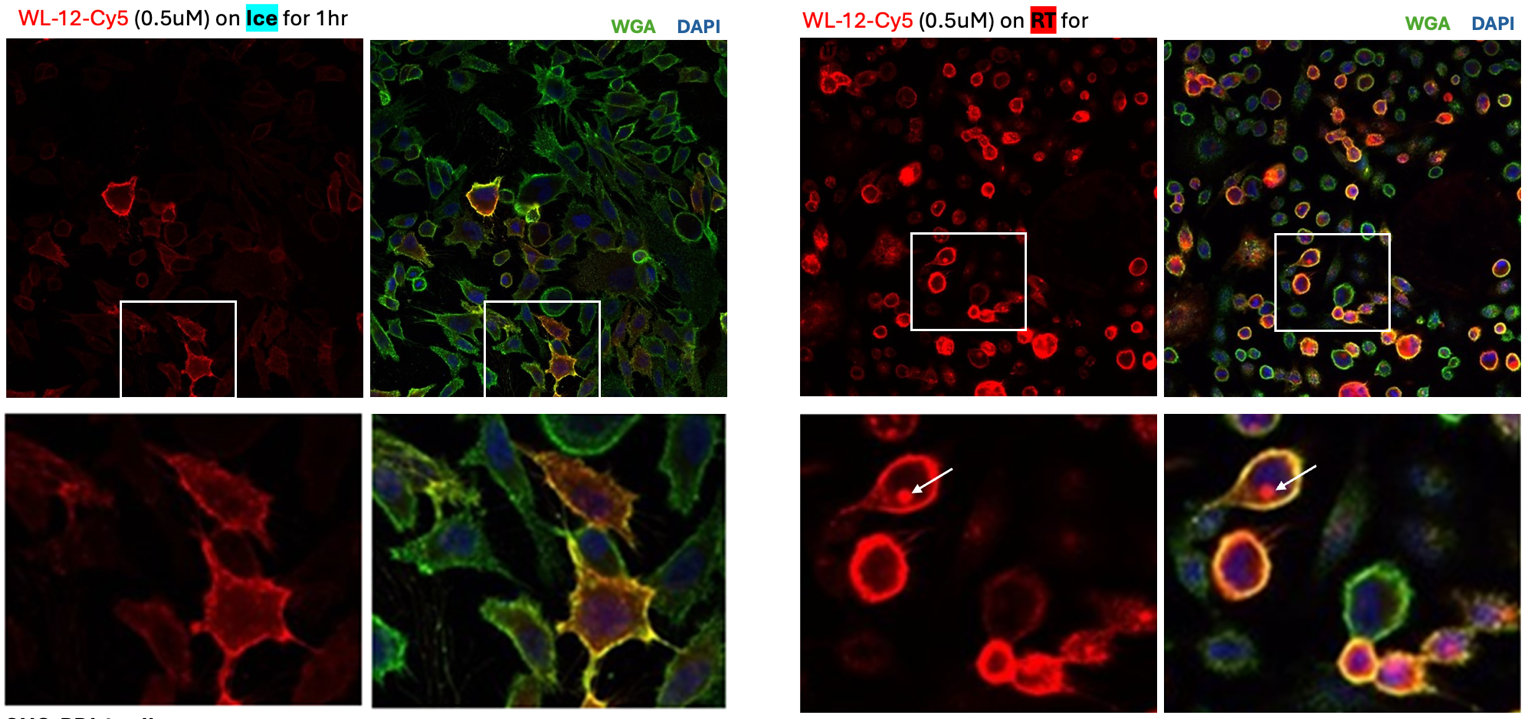

**c	d**

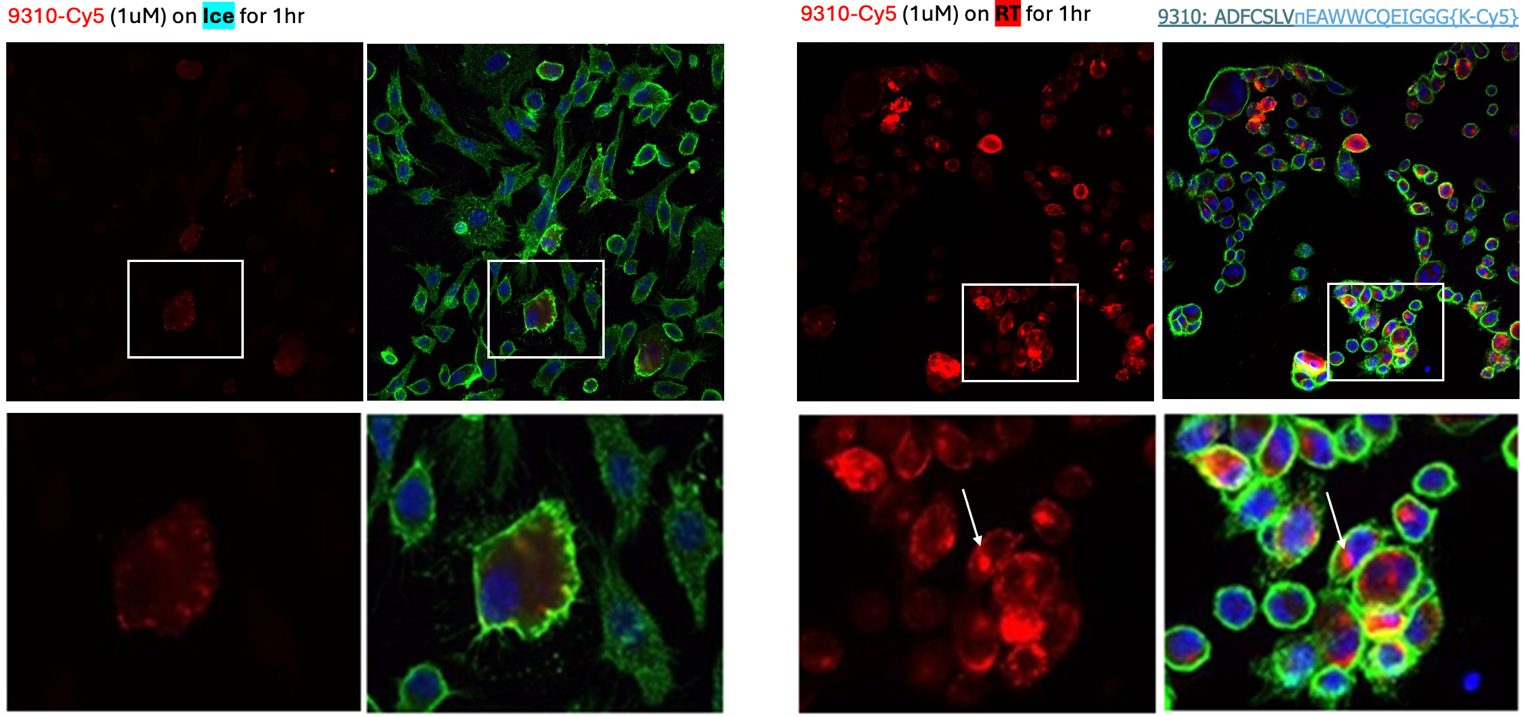

### Supplementary Figure 11: Internalization of PD-L1 targeting macrocycles

We employed PD-L1-overexpressing CHO cells with WL-12-Cy5 as a positive control (A,B) and macrocycle 9310 ADFCSLVπEAWWCQEIGGG[K-Cy5] as a test (C,D). Incubations were performed in the presence of 500 nM peptide at room temperature (B,D) or on ice (A,C). Cells were counterstained with DAPI and Wheat Germ Agglutinin (WGA) and imaged by laser-scanning confocal microscopy.

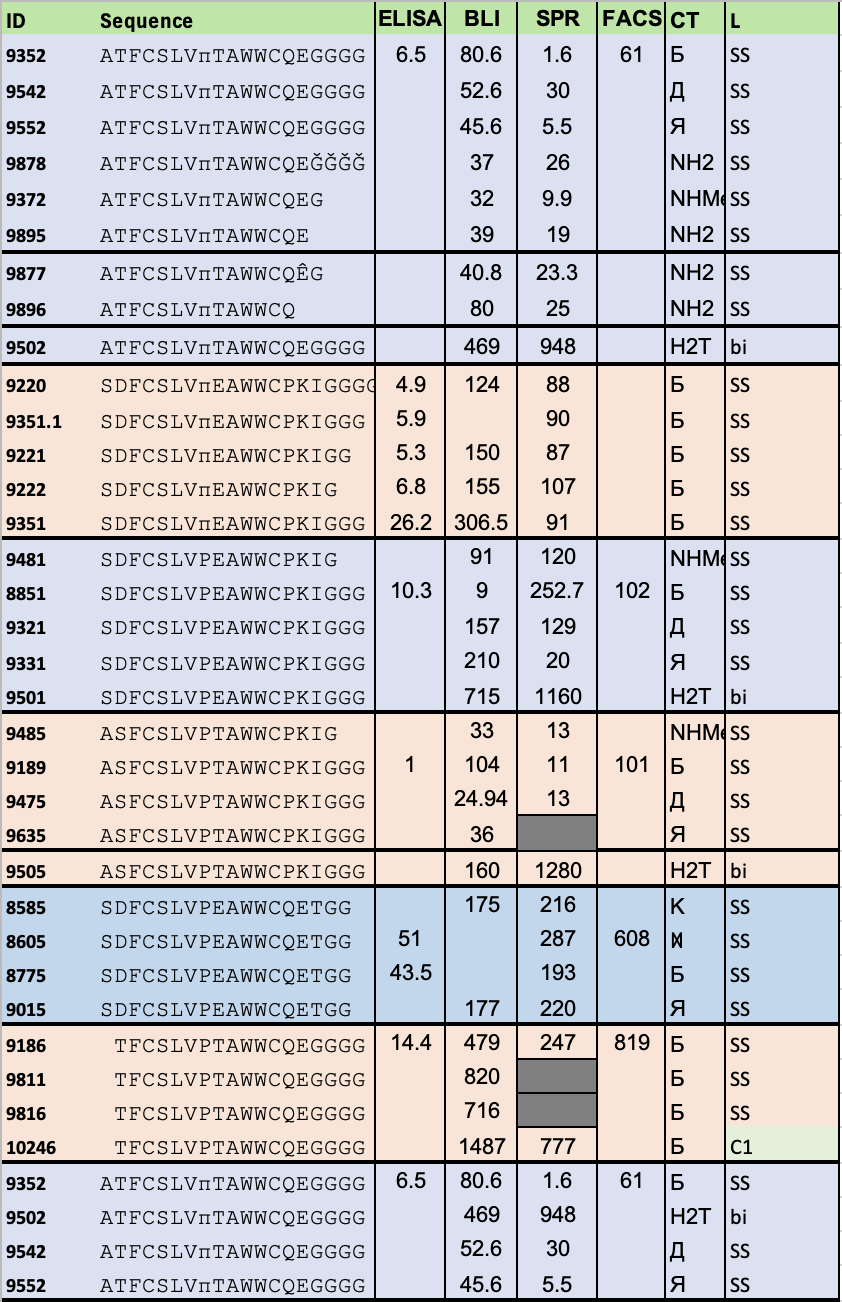

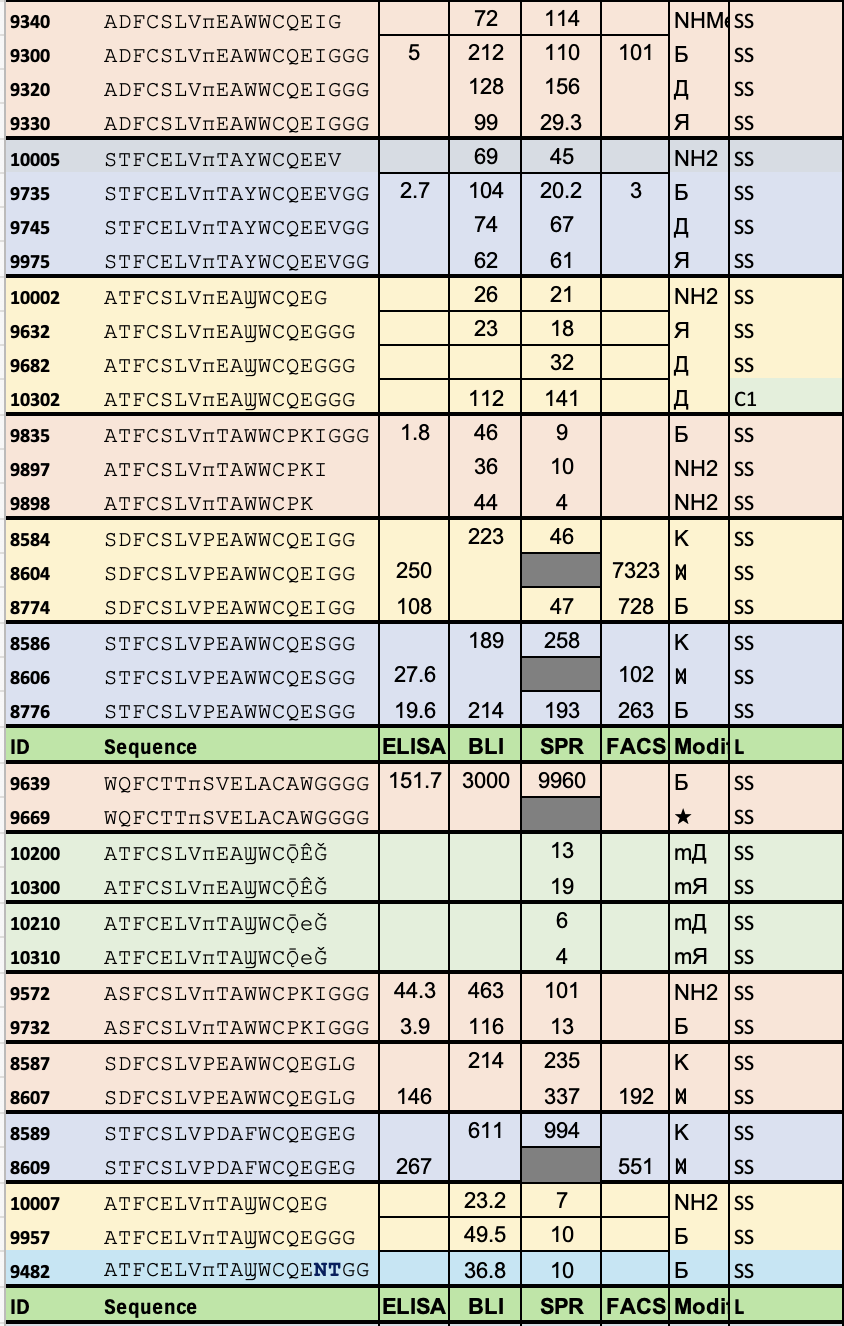

### Supplementary Figure 12: Effect of C-terminal modifications on binding of macrocycles

The tables summarize the peptides with identical amino acid sequences and two or more modifications to peptide macrocycles at the C-terminus, as well as the effect of these modifications on measurements in 4 different assays. The last two entries 9957 and 9482 illustrate a curious case where GG to NT replacement in C-terminus had no effect on peptide activity. Empty cells or grey cells denote missing measurements, “-2” indicates that the compound has been measured by SPR, but no signal was observed. For abbreviations of the C-terminal modifications (CT), see main text Fig. 4g. “L” column denotes the cyclization strategy by disulfide (SS), methylene linchpin (C1, see Fig. 8d), or head-to-tail bicycle (bi, see Fig. 5c)

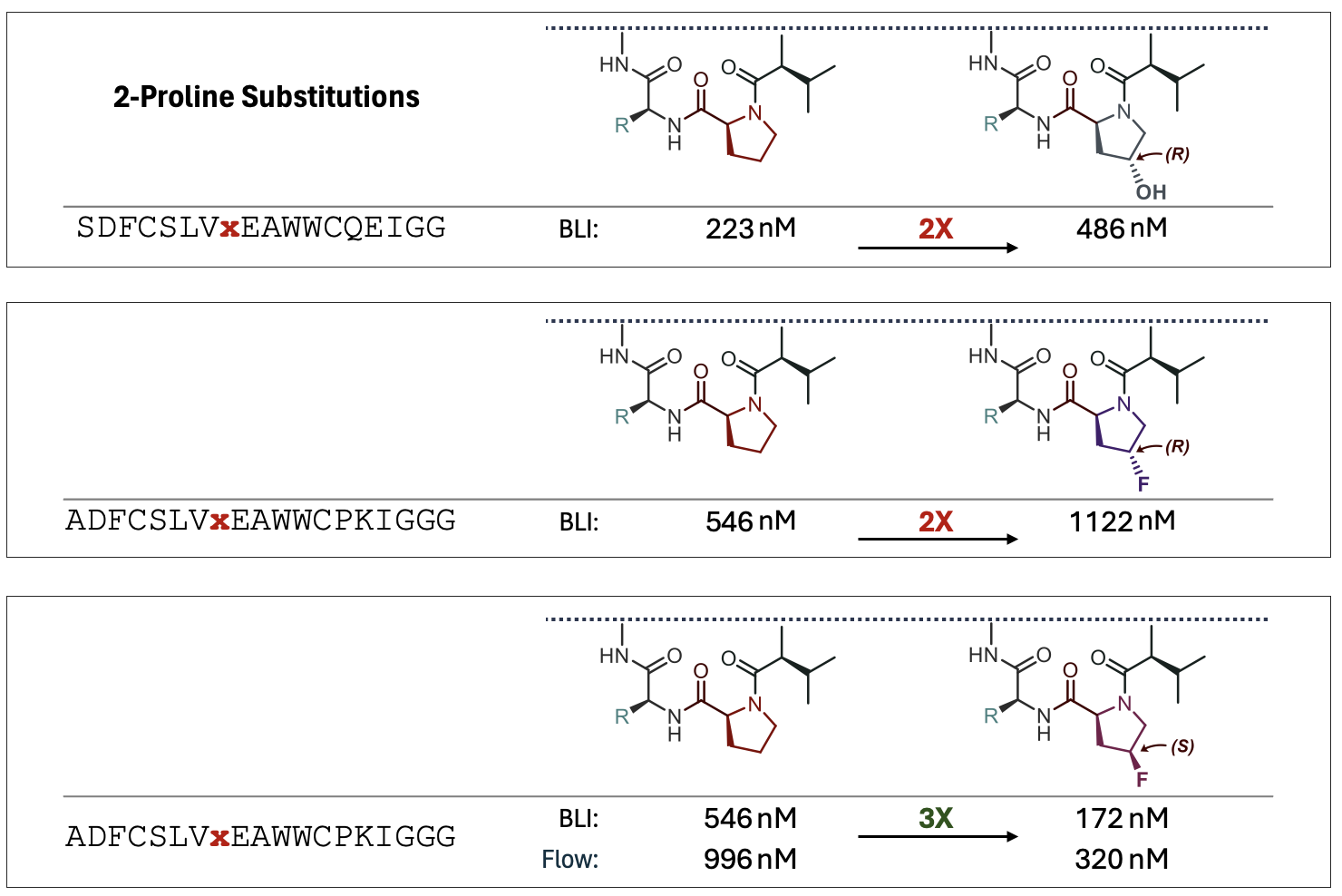

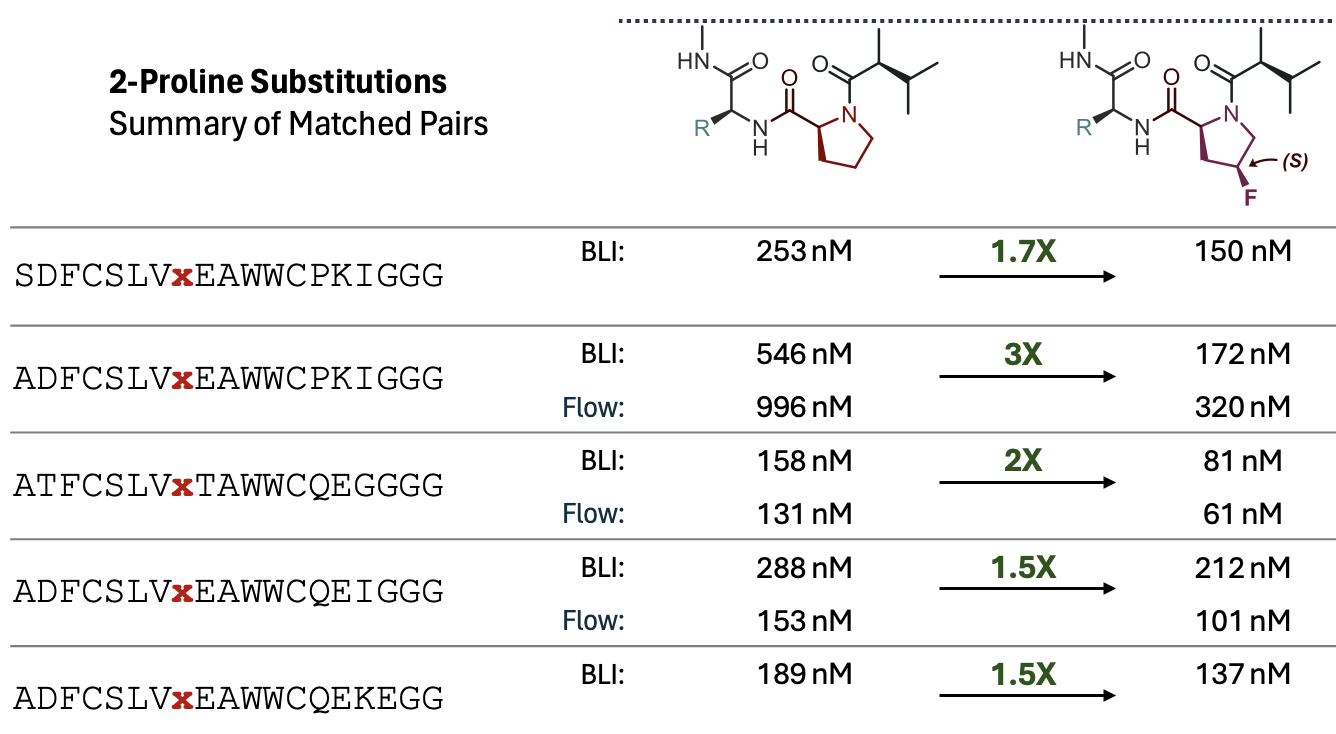

### Supplementary Figure 13: Effect of replacing Pro8 by proline analogs on PD-L1 binding activity

We tested replacement of Pro in six different peptides sequences and in one sequence we tested both R and S isomers of the fluoroproline. We observed that S isomer was generally favourable whereas R isomer had negative effect on the performance. The binding curve corresponding to two epimers of fluoroproline in the same peptide can also be found in Supplementary Fig. 1d.

**a**

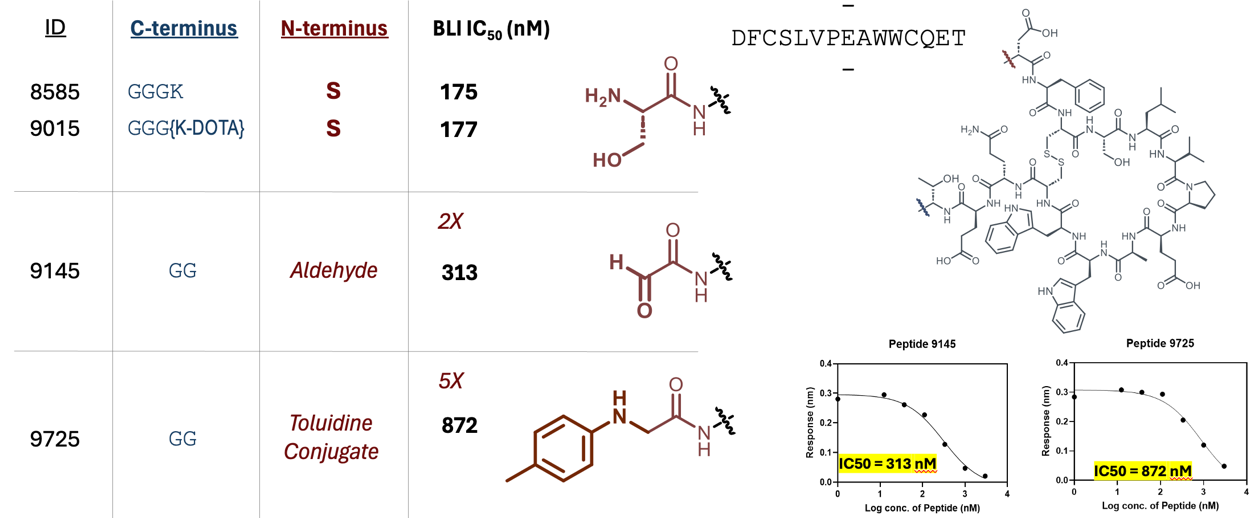

**b**

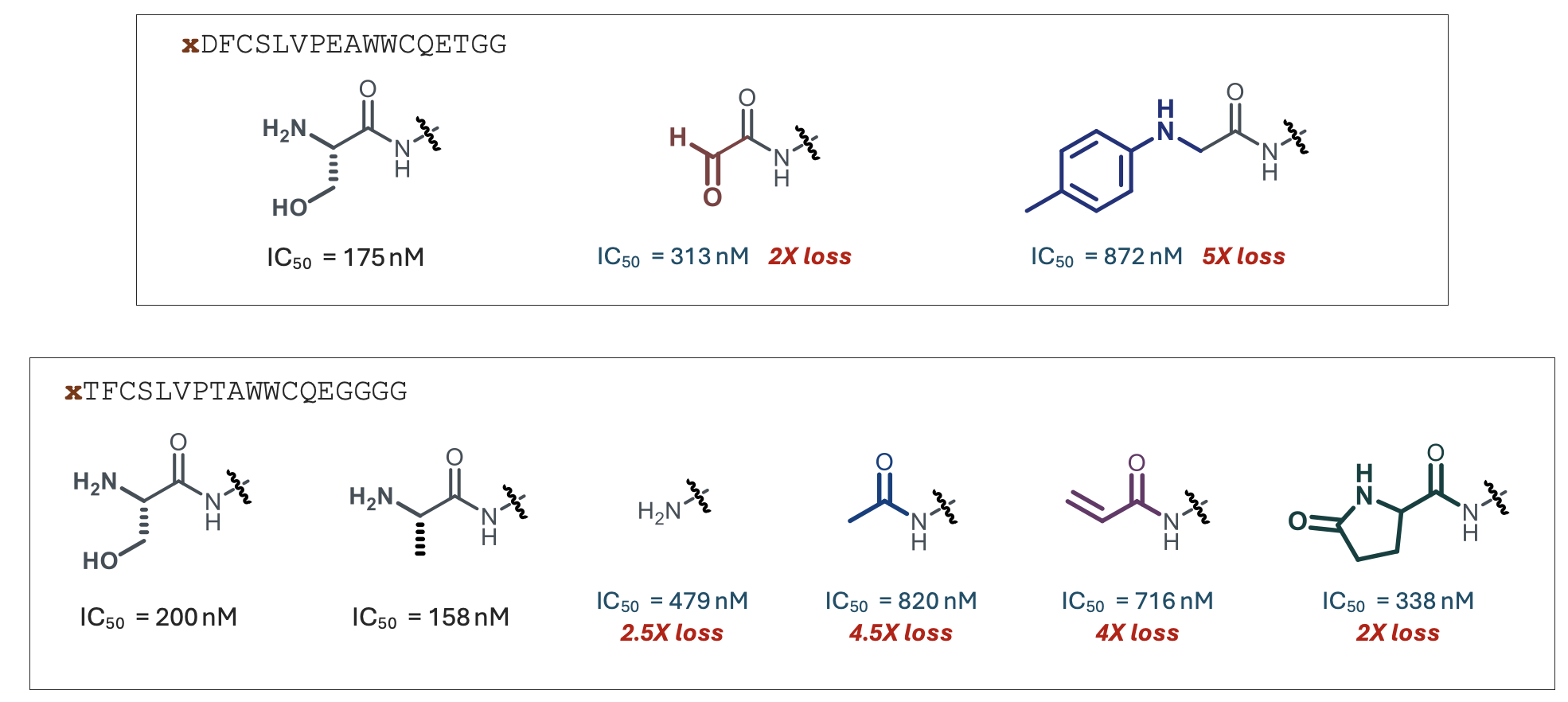

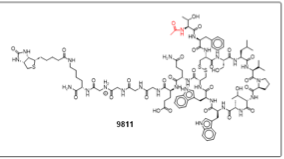

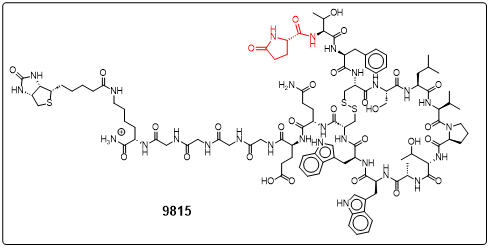

**c**

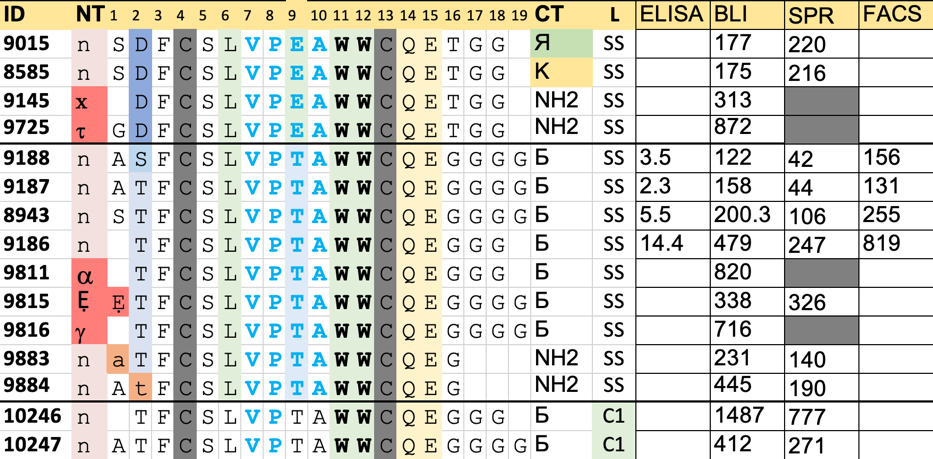

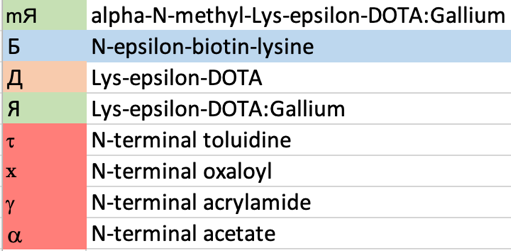

### Supplementary Figure 14: SAR of macrocycles with respect to N-terminal modifications

**a,** Changing the N-terminal Ser in **S**DFCSLVPEAWWCQET to x (oxaloyl) and t (N-toluidine-glycine) leads to 2-5 fold loss in the inhibitory potency. **b,** N-terminal changes in peptide **S**TFCSLVPTAWWCQEGGGGK(e-biotin) to replace the native N-terminal Serine (**8943**) with Alanine (9187) leads to modest enhancement. N-terminal deletion leads to 2.5x loss (**9186**), capping with N-terminal acetylation (a, **9811**), acrylamide (g, **9816**) and pyroglutamate (Ẹ, **9815**) gave rise to 2-4x loss in inhibitory potency. Inversion of the chirality of the N-terminal Ala and penultimate Thr gave rise to 4-5x loss in potency. **c,** summary of all changes in tabular form

**a** Predictions made in chemically-modified focused libraries.

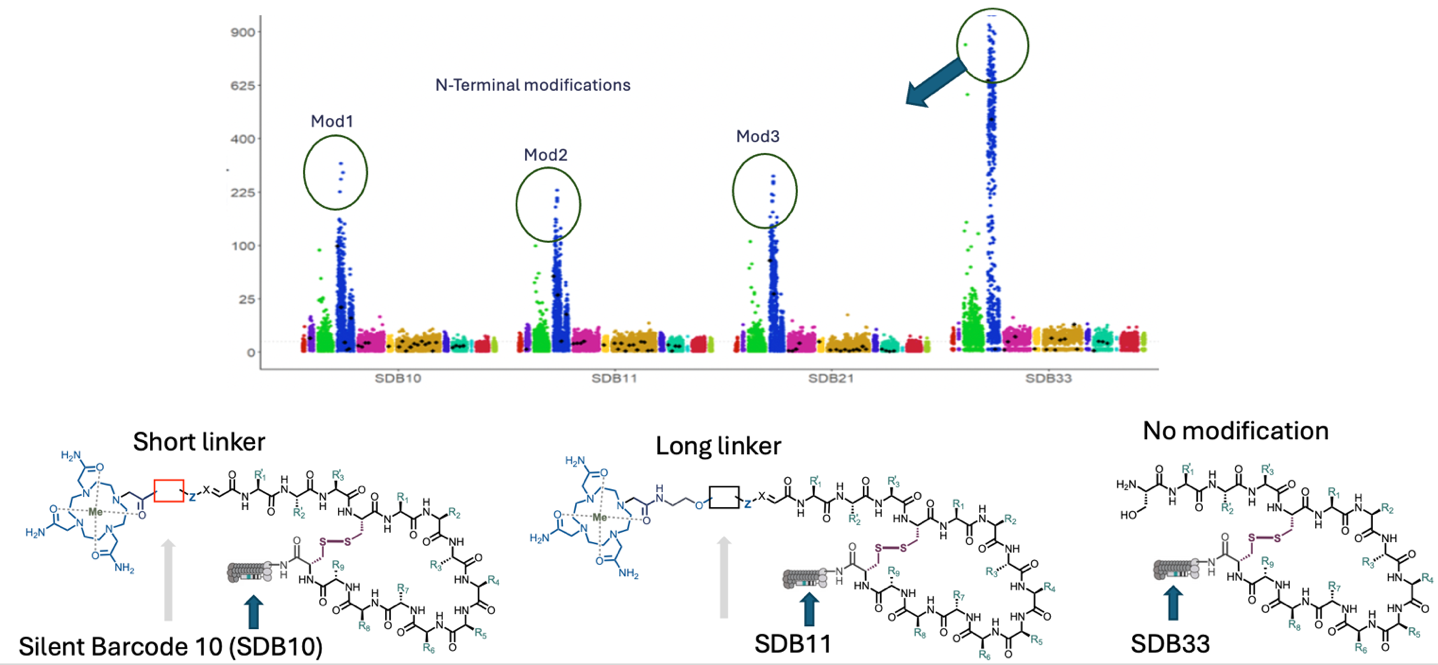

**b** Validation of the predictions in synthetic peptides

| 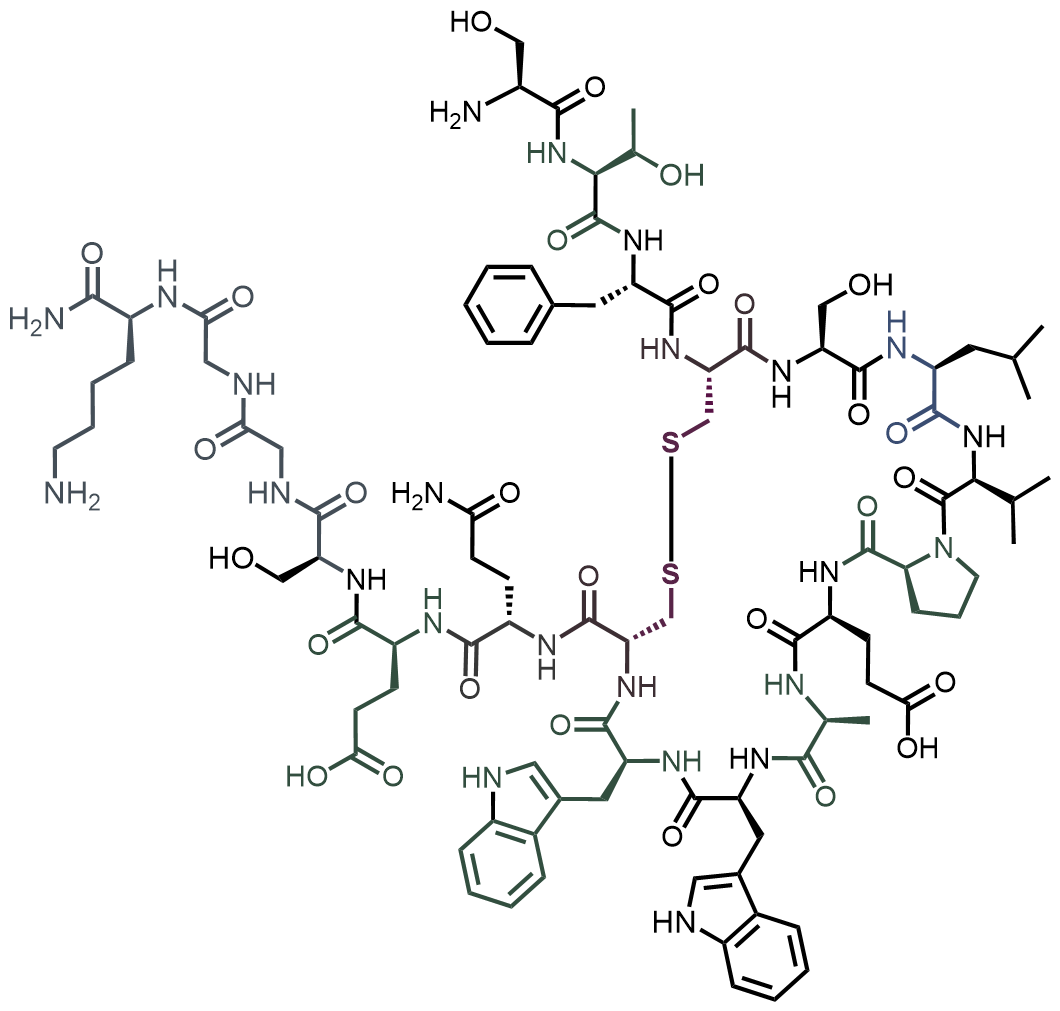 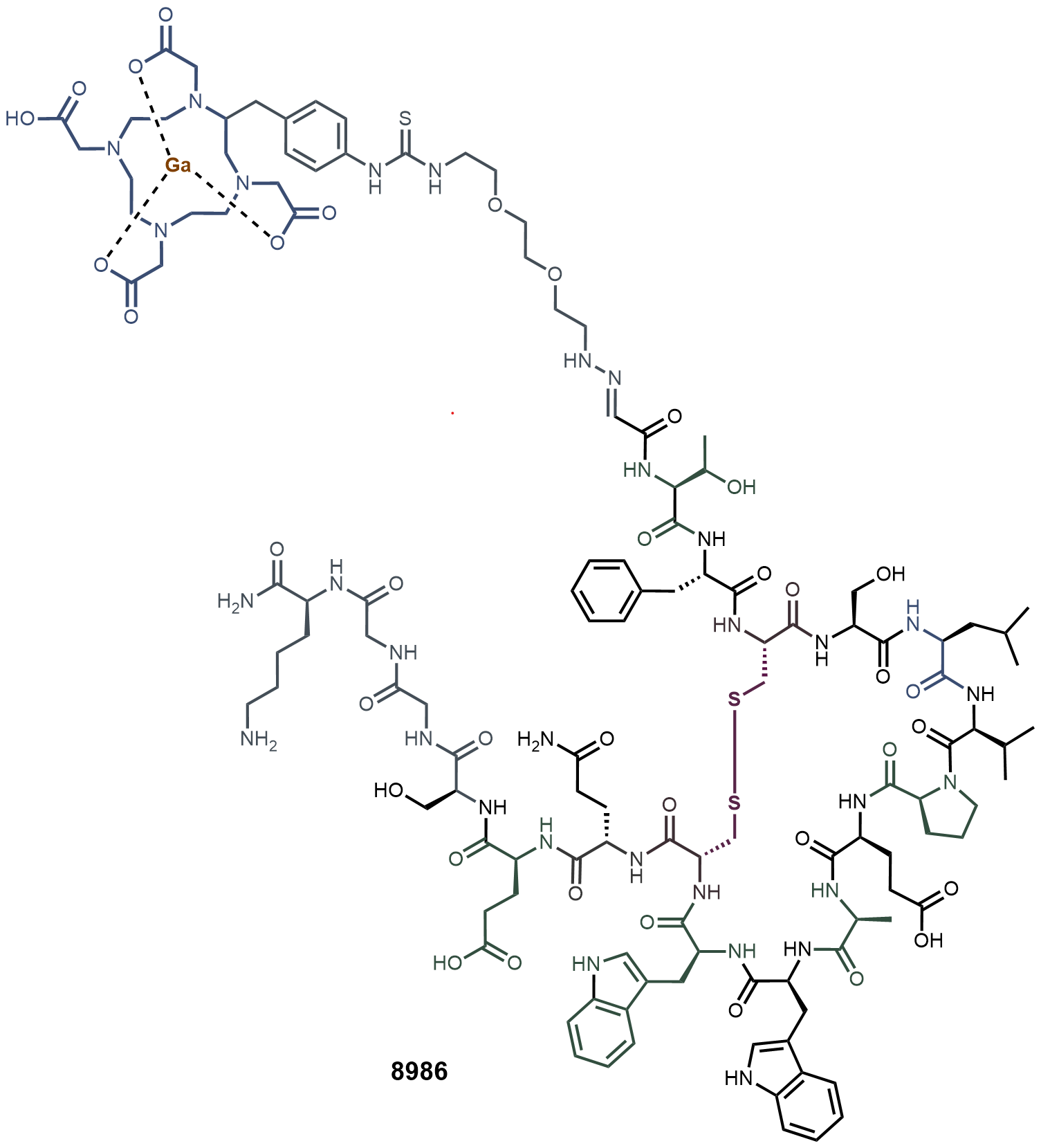  **8586** (IC_50_ = 187 nM ) **9896** (IC_50_ = 3098 nM ) | 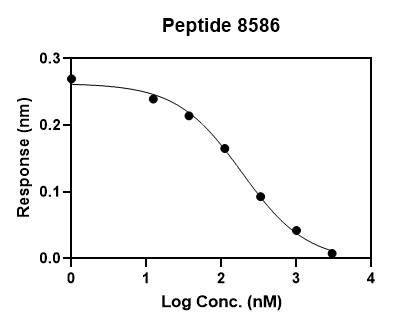  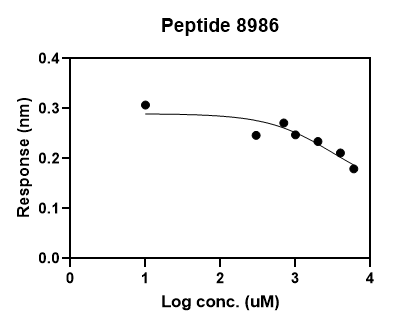 |
| --- | --- |
| **c** Effect of N-terminal biotinylation |  |

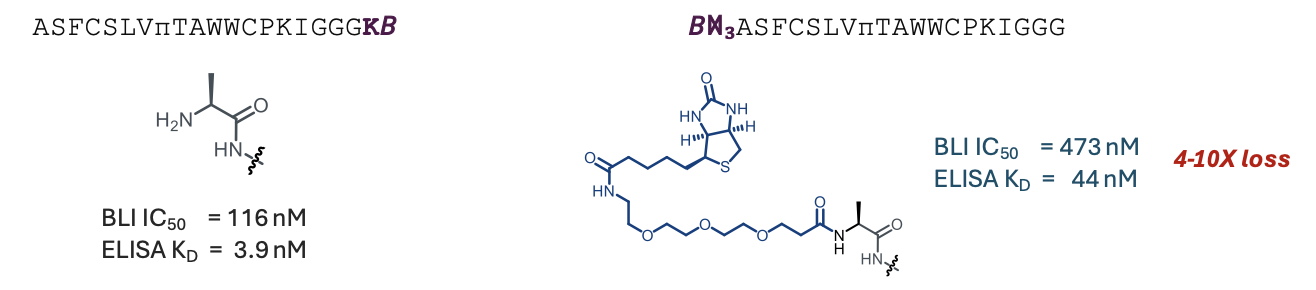

### Supplementary Figure 15: The effect of N-terminal ligation of DOTA-chelators or biotin

**a,** We cloned four focused libraries that encode the same amino acid compositions but employ different codons similarly to “silent DNA barcoding” reported in previous publications^4^. Each library has been treated with NaIO_4_ to convert N-terminal Ser to aldehyde, and subsequently DOTA-chelators with aldehyde reactive hydrazine groups (-Z-X= is -NH-N=) have been ligated to the N-terminus. We varied the length and the composition of the linker but overall, we observed that all N-terminal modifications with DOTA linkers significantly diminished the enrichment factor of nearly all peptides. Most peptides decreased their FC by factor of 5-10 upon N-terminal modification. Based on the calibration studies, such decrease pushes the activity of all such constructs to high nanomolar to low micromolar range. Between 4-10 peptides exhibited lesser penalty but even such intermediate decrease in FC can translate to 10x decrease in potency. **b**, Measurement of the inhibitory potency of the synthetic peptides with and without N-terminal chelator confirmed the detrimental effect of this modification predicted by SDB-GS libraries. **c**, N-terminal acylation with biotin decreased activity by factor of 3-10x.

**a**

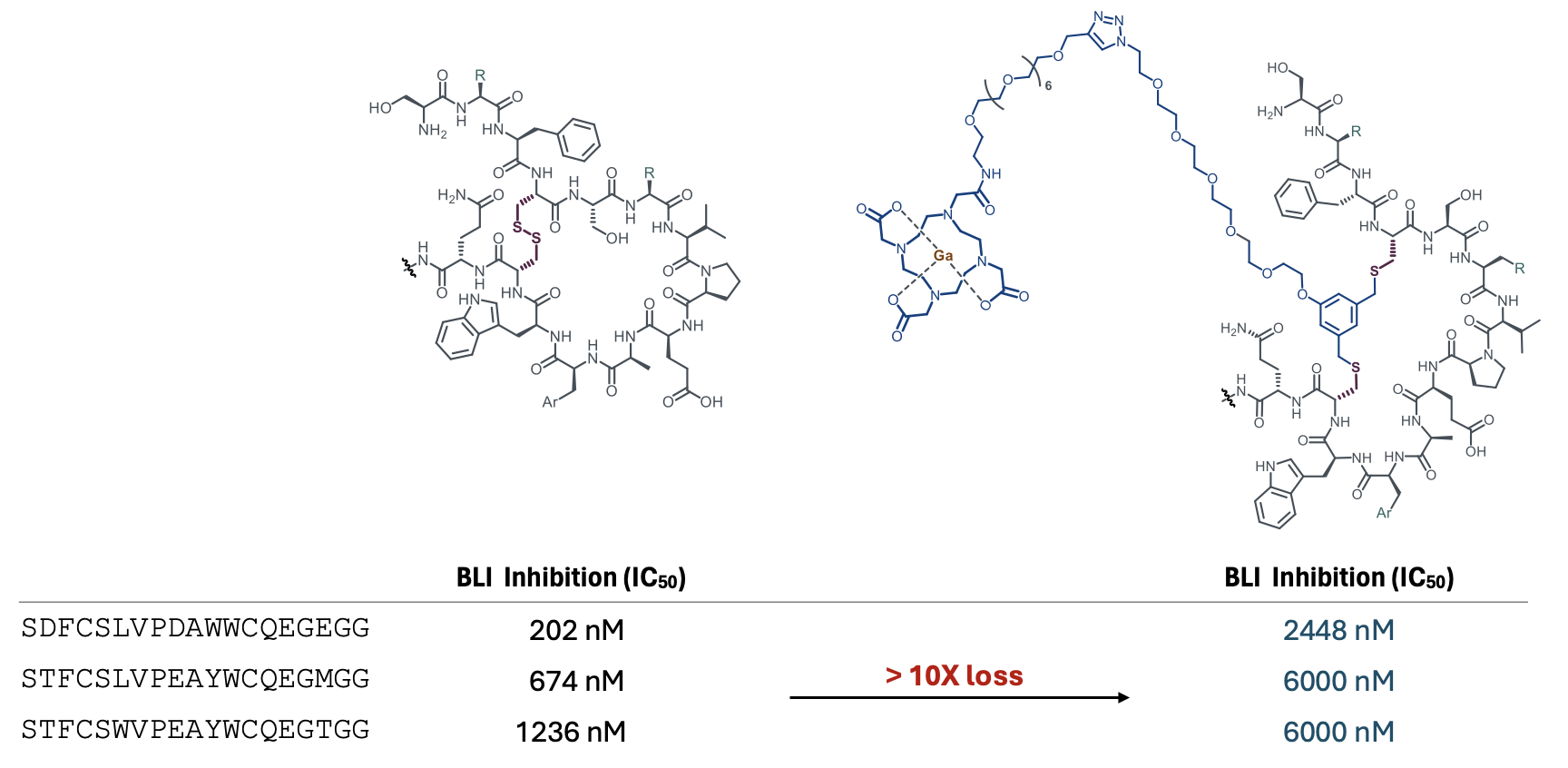

**b	c**

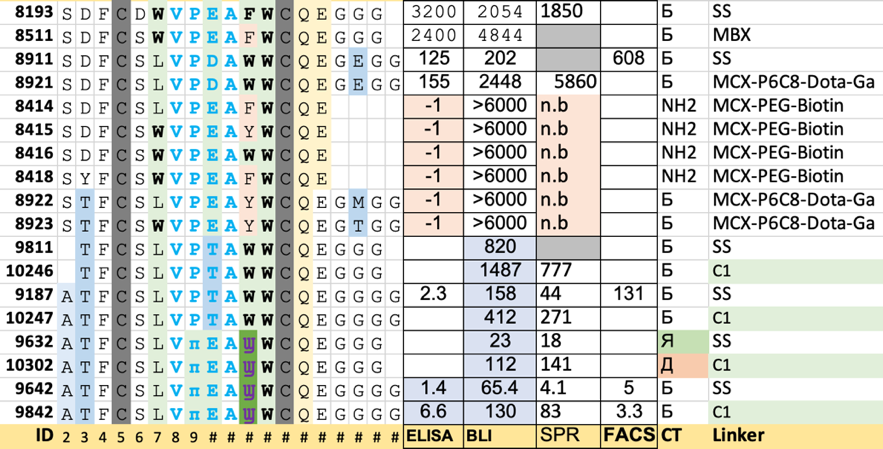

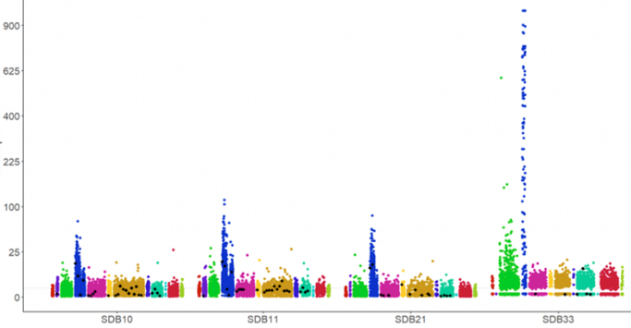

**d**
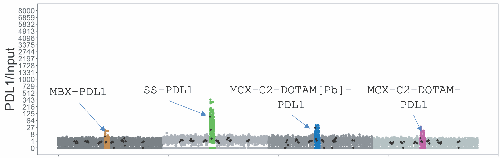

### Supplementary Figure 16: Modification of peptides by linchpins with DOTA-chelators

**a,** Representative structure of a synthetic peptide modified by metachloroxylene (MCX) with DOTA chelator complexes to gallium (MCX-P6C8-DOTA-Ga); the table below illustrates that this linchpin resulted in nearly complete loss of activity in three different peptide sequences. **b,** Summary of other attempted changes to the disulfide bridge using metabromoxylene (MBX), MCX-P6C8-DOTA-Ga, MCX-PEG-biotin or one-carbon linchpin (1C, see main text Fig. 8D). **c**, Effect of library-wide ring expansion using three different DOTA-containing linchpins that were used to modify three chemically identical focused libraries associated with silent barcoded populations SDB10, SDB11 and SDB21. The fourth unmodified library was associated with barcode SDB33. All linchpins resulted in >10x decrease in FC (enrichment) for all lead PD-L1-binding peptides present in these focused libraries when compared to unmodified disulfides (SDB33). **d,** We modified focused library GS29 by MBX or DOTAM-modified MCX linchpins in three silently barcoded sub-libraries, added disulfide-constrained GS29 library and panned the mixture in the same solution. As in c, all linchpins decreased FC in panning on PD-L1.

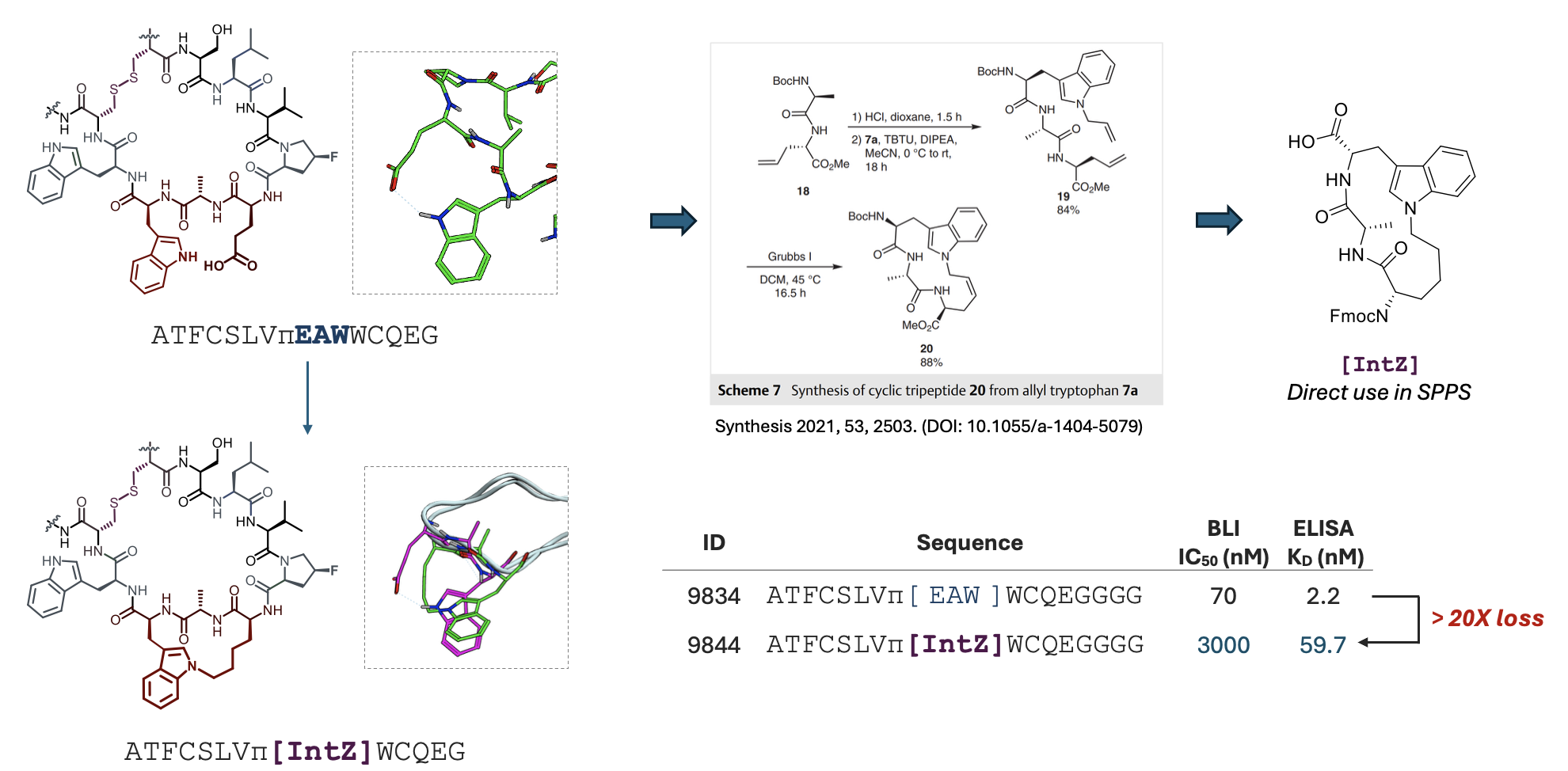

### Supplementary Figure 17: Bicyclization by cross-linking of i:i+2 side-chain residues

Modeling of the macrocycle architecture and its docking pose suggested proximity of E9 and W11 side chains. We devised a synthesis of the tripeptide scaffold following previously reported route and incorporated the resulting tripeptide into the macrocycle. Unfortunately, the resulting structure completely lost its inhibitory potency and exhibited >20x reduction of potency in ELISA assay. Given a critical role of W11 in other SAR experiments, this result is not entirely surprising. Most attempts to replace W11 with even structurally similar ncAA resulted in substantial decrease of activity. Future strategies for i:i+2 bicyclization should employ an X-ray structure, which was generated after the i:i+2-bicycles described in this figure have been synthesized and tested.

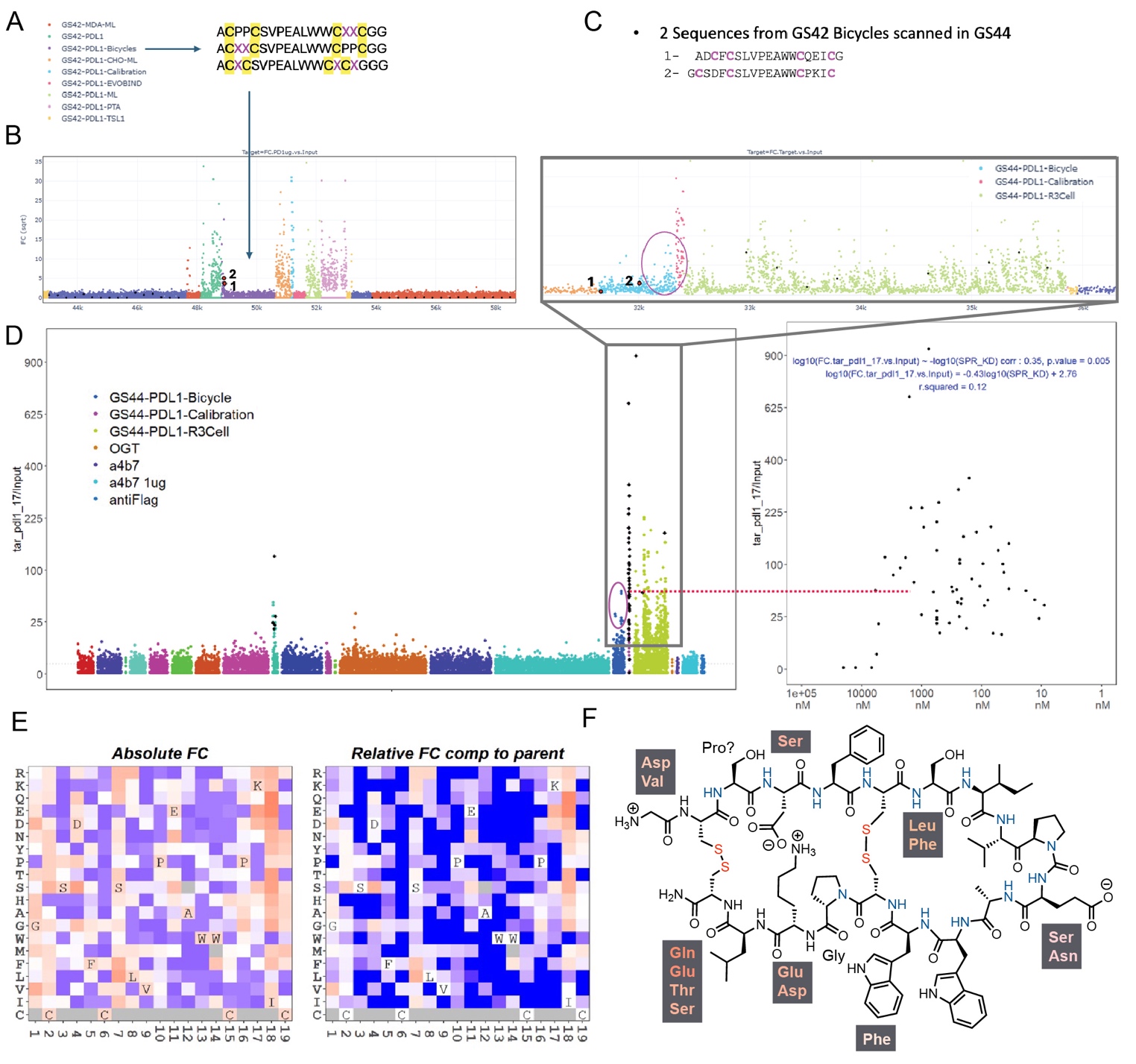

### Supplementary Figure 18: Attempted conversion of Cys-monocycles to tetra-Cys-bicycles

**a,** We introduced CPPC motifs at the C and N-terminal regions of the peptides and we produced a focused library GS42 containing several hundred derivatives, such as mutations, insertions and deletions. **b,** While other classes of PD-L1-binding sequences present in GS42 exhibited FC=5-30 enrichment above the baseline, nearly all tetracysteine bicycles in GS42 had undetectable PD-L1 binding (indistinguishable from baseline). Exceptions were two peptides (highlighted as 1 and 2) that had minor level of enrichment. **c-d,** Positional scan and insertion-deletion scan of these sequences identified 2 and 5 variants with enrichment above the baseline. **c,** is a zoomed-in view of full Manhattan plot in (**d**), along with a calibration of FC enrichment in binding to PD-L1 protein versus K_D_ of synthetic peptides as measured by SPR. **e,** Deep mutational scan of sequence 2, GCSDFCSLVPEAWWCPKIC suggested several improvements in several locations of the bicycle (**f**). Based on built-in calibration of GS44 (**d**), we anticipate these bicyclic derivatives to be of high nanomolar K_D_.
